# “It All Rolls Downstream: Upstream Control of Physical Activity Regulation”

**DOI:** 10.1101/2023.05.10.540028

**Authors:** Brianne M. Breidenbach, Liwen Liu, Troy La, Tatiana N. Castro-Padovani, Nathan Keller, Linda S Pescatello, Matthew M. Robinson, Scott A. Kelly, Kevin Gerrish, J. Timothy Lightfoot

## Abstract

Physical activity is regulated by a variety of genetic molecules. However, the pathways through which those molecules work to regulate activity is largely unknown. The purpose of this study was to gather the known genetic molecules that are associated with activity regulation and define overall upstream regulator pathways through which these molecules work. We conducted a systematic review to gather all available published datasets related to physical activity regulation, standardized the data for genomic location and species, and used this data, in an unbiased manner to create a dataset that was used: (1) to physically map and visualize all identified molecules to homologous chromosome locations and (2) as the dataset for which an Upstream Regulator Analysis (URA) was conducted using Qiagen Ingenuity Pathway Analysis (IPA) software. Our search resulted in 469 genetic molecules (e.g. genomic variant, transcript, protein, micro-RNA) that were split into brain (n=366) and muscle (n=345) sub-groups, which was our attempt to separate differences in central vs peripheral pathways. The brain and muscle data sets had several potential upstream regulators, the top-rated being β-estradiol as a regulator for 19.5% and 21% of the brain and muscle datasets respectively. To our knowledge, β-estradiol’s identification as a potential regulator, is the first evidence to link the well-known effects of sex hormones on physical activity with genetic regulation of physical activity. There were a variety of potential upstream regulators for the molecules collected in this review, but interestingly, three of the top five for both brain and muscle are nuclear receptor binding ligands; estradiol (estrogen receptor), dexamethasone (glucocorticoid receptor), and tretinoin (retinoic acid receptor), indicating a potential role of nuclear receptors in the regulation of physical activity. Selective nuclear receptor modulation may be an area of interest in future mechanistic studies of the genetic regulation of physical activity.

## Introduction

Physical activity has been broadly defined “as purposeful exercise or movement that expends a significant amount of energy”^1^, and more specifically defined as any bodily movement produced by the contraction of the skeletal muscles resulting in an increase in caloric requirements over resting energy expenditure^2^. Several published papers have established that daily physical activity levels have a significant genetic control element^3^, with observed heritability of physical activity control ranging from 20-92%^4,5^ depending upon the model, analysis method, and duration of data collection. Common environmental factors appear to play a small, if negligible role in controlling daily activity, while unique environmental factors play roles similar in magnitude to genetic factors^3^ (e.g. factors such as diet and the presence of environmental toxicants).

Overall, there are a wide-variety of genetically controlled physiological mechanisms that can influence physical activity -including various neurotransmitters^1,6–8^, muscle energy sources^9^, muscle calcium handling mechanisms^10^, and/or substrate availability^11,12^. It has been hypothesized^13^ and generally accepted^3^ that the genetic molecules that regulate physical activity function at the central level (i.e., brain/neural) and/or at the peripheral level (i.e., skeletal muscle). Data supports gene interference, gene silencing, and/or overexpression at either the central and peripheral levels can significantly influence daily activity^10,14–16^. While sex hormones have long been shown to influence daily activity^17–21^, it is unclear whether the actions of the sex hormones on physical activity are mediated through genetic mechanisms (e.g., genetic variation in hormone release/receptor activity) or through direct biological mechanisms (e.g., inhibition of a pathway). The literature establishing that genetics are a significant regulator of daily physical activity is extensive; however, a critical gap remains surrounding the integration of these molecules into multi-omics datasets and the identification of common upstream pathways controlling the regulation of daily physical activity.

The development of multi-omics datasets is most often accomplished through ‘omics’ data within the same model^22^ (e.g., genomic variation, proteomics, transcript expression variation). In the area of “genetic regulation of physical activity”, the challenge faced is the lack of ‘omics’ datasets in the same set of samples. Only one group to our knowledge has produced multiple omics datasets in the same model^10,15,23–25^. Thus, developing an ‘omics’ dataset of physical activity regulation requires the combining of datasets from various investigators, species, and models. Understanding the systems genetics network(s) involved in physical activity regulation will help the future synthesis and translation of a general mechanistic understanding of the upstream regulators that control physical activity. Given that the last summary of the known genomic variants associated with physical activity level was published in 2012^26^, and that further data regarding transcriptomic, proteomic, and other molecular variants associated with physical activity have been published since then; the purpose of this project was to (i) present an updated summary of the currently known manuscripts and molecules associated with, and mechanistically linked with daily physical activity regulation; (ii) provide a novel visualization of genomic-linked regulators based on chromosome location (mouse and human) and (iii) to identify potential common-upstream regulators of the identified molecules associated with physical activity.

## Methods

### Overview

We conducted a comprehensive systematic review to gather all available published datasets related to physical activity regulation, standardized the data for genomic location and species, and used this data to create a dataset that was used: (1) to physically map and visualize all identified molecules to chromosome locations and (2) as the dataset for which an Upstream Regulator Analysis (URA) was conducted using Qiagen Ingenuity Pathway Analysis (IPA) software (Qiagen Inc., https://digitalinsights.qiagen.com/IPA, Germantown, MD, USA).

### Literature Search

To create our omics dataset, the PubMed/MEDLINE database^27^ was utilized to identify manuscripts containing “omic” data (e.g. QTLs, eQTL, miRNA, proteomics) related to physical activity regulation in humans and mice. To build up the literature search in PubMed, we used the following MeSH-terms: Mice, Humans, Motor Activity / genetics, proteome, gene expression profiling, microRNAs, quantitative trait loci, and exercise. Using the same criteria, several additional databases were searched through the University-Library services to locate additional manuscripts (e.g. ERIC, ScienceDirect, Web of Science, Google Scholar). Author expertise and knowledge was also utilized to verify and confirm included manuscripts. Initial searches were completed on April 1, 2020 and subsequent verification of these searches was completed on October 12, 2020, returning a total of 251 potentially qualifying manuscripts. Two authors independently applied eligibility criteria, assessed the quality of the study, and extracted and coded data. We contacted study authors for more information where necessary. Studies were selected based on the following inclusion criteria: (1) original research papers with prospective or retrospective design, (2) physical activity was the primary dependent variable [e.g. wheel running, spontaneous cage activity, locomotion activity, accelerometry data, self-reported physical activity], (3) contained some variant of ‘omics’ data (e.g., genomic variation, proteomics, transcript expression variation) as an independent variable that were found to have either been associated with or causative of (i.e., directly affect) physical activity. Manuscripts showing up in multiple searches were counted only once, and review or summary papers were not retained. Studies were excluded when exercise/physical activity was the independent variable (e.g. study was not intended to identify genetic response to exercise). In total, 46 manuscripts qualified for data extraction (Table 1).

**Table 1.** Citations reporting genetic molecules associated with/causative of physical activity

| First Author / Year | Type of genetic molecule reported | Citation in Bibliography | First Author / Year | Type of genetic molecule reported | Citation in Bibliography |
| --- | --- | --- | --- | --- | --- |
| Bruneau 2018 | Genomic variant | 33 | Keeney 2012 | Proteomic | 34 |
| Caetano-Anolles, K, et al. 2016 | Transcriptomic | 35 | Klimentidis 2018 | Genomic variant | 36 |
| Cai 2006 | Genomic variant | 37 | Knab 2009 | Proteomic | 1 |
| Comuzzie 2012 | Genomic variant | 38 | Kostrzewa 2014 | Genomic variant | 39 |
| Dawes 2014 | Transcriptomic | 23 | Kumar 2010 | Transcriptomic | 40 |
| Dawes 2015 | miRNA | 24 | Leamy 2008 | Genomic variant | 41 |
| De Moor 2007 | Genomic variant | 42 | Leamy 2010 | Genomic variant | 43 |
| De Moor 2009 | Genomic variant | 44 | Leamy 2011 | Genomic variant | 45 |
| Doherty 2018 | Genomic variant | 46 | Lightfoot 2008 | Genomic variant | 25 |
| Dubreucq, 2012 | Proteomic | 16 | Lightfoot 2010 | Genomic variant | 47 |
| Eisener-Dorman 2010 | Transcriptomic | 48 | Lin 2018 | Genomic variant | 49 |
| Ferguson 2014 | Proteomic | 10 | Maestu 2013 | Genomic variant | 50 |
| Ferguson 2015 | Proteomic | 15 | Nehrenberg 2010 | Genomic variant | 51 |
| Furuse 2002 | Transcriptomic | 52 | Pistilli 2011 | Proteomic / Transcriptomic / Genomic Variant | 53 |
| Gielen 2014 | Genomic variant | 54 | Saul 2017 | Transcriptomic | 55 |
| Hartmann 2008 | Transcriptomic / miRNA | 56 | Simonen 2003 | Genomic variant | 57 |
| Hillis 2020 | Genomic variant | 58 | Steffan 2002 | Genomic variant | 59 |
| Jean 2012 | Proteomic | 60 | Tsao 2001 | Proteomic | 9 |
| Kas 2009 | Transcriptomic / Genomic Variant | 61 | Umemori 2009 | Transcriptomic | 62 |
| Kelly 2010 | Genomic variant | 63 | Wilkinson 2013 | Genomic variant | 64 |
| Kelly 2012 | Transcriptomic | 13 | Williams 2014 | Transcriptomic | 65 |
| Kelly 2014 | Transcriptomic | 66 | Yang 2009 | Transcriptomic | 67 |
| Keeney 2008 | Proteomic | 68 | Yang 2012 | Genomic variant | 69 |

### Data Extraction and Standardization Methods

From qualifying studies, 464 molecules were identified (Supplemental Table 1), and subdivided into muscle (n=344) and brain (n=361) subsets. Subset assignment was based on whether or not the tissue type was identified in the original manuscript. The majority of genomic variant related manuscripts did not indicate which tissue/sample was utilized for extraction; when this occurred, the molecule(s) were assigned to both datasets (e.g. brain/muscle; Supplemental Table 1). However, when the tissue type was indicated (e.g. soleus muscle vs striatum), the molecules were assigned to either the muscle or brain set. Our intention was to separate differences in central vs peripheral networks, but in reality, the two networks had a large amount of overlapping data points due to lack of disclosure of tissue extraction type for genomic variants.

Integration of omics data from the 46 articles required standardization of genomic location and cross-species comparisons. Genomic locations associated with physical activity from earlier papers [where microsatellites were used as genomic markers] were converted from ‘centimorgans’ to ‘base pairs’ utilizing a publicly available converter that was recently decommissioned (http://cgd.jax.org/mousemapconverter/). Genomic locations of the identified variants were updated to the build of the specific species current at the time of data collection for this systematic review (for human GRCh38.13, ref.^28^, for mice: GRCm39, ref.^29^). Cross-species comparisons of genetic molecules require translation into a common species to allow these analyses. Given that many of the genetic molecules associated with physical activity (miRNA, transcripts, proteome, and genomic variants) have been identified in mouse models, genomic locations for the molecules in humans (all genomic variants) were converted to the homologous genome location within the mouse chromosome. Ensembl (https://ensembl.org) was used to convert the human genetic molecules to the mouse homologs^30^.

Following extraction and standardization, and when possible, the molecules were: (1) mapped to their chromosome location using the online program MG2C^31^ (mg2c.iask.in) with the molecules only found in humans (i.e., not homologous) mapped to the human chromosome and (2) uploaded and linked to molecules in the IPA database for Upstream Regulator Analysis (URA) (Supplemental Table 3,4). Kramer et. al (2014) describes the algorithms, tools, and visualizations in IPA utilized for the URA analysis^32^. Briefly, URA determines likely upstream regulators that are connected to dataset genes through a set of direct or indirect relationships^32^. Our dataset did not include expression-fold change, expression ratios, or expression log ratios. Consequently, the URA is unable to predict activation status (activated or inhibited) but may be a useful tool to investigate mechanistic targets in future studies. In the results, the p-value of overlap indicated the most significant upstream regulators that influence target molecules from the dataset.

## Results

The resulting papers from our systematic literature review (Table 1) netted a total of 469 genetic molecules (Supplemental table 1) that have either been associated with or causative of (i.e., directly affect) physical activity. Genomic variants made up the majority of molecules (n=225, 48%) while other types of genetic molecules were less prominent in the literature (proteomic, n=61, 13%; transcript, n=139, 30%; and miRNA, n=44, 9%).

### Location Mapping

Of these molecules, 385 were mappable to the mouse chromosome map (Fig. 1) and an additional 74 were mappable only to the human chromosome map (Fig. 2). Ten of the molecules previously found (3 transcripts and 7 genomic variants) were unmappable due to outdated markers and locations that could not be standardized. Of the 116 human genomic variants associated with physical activity (Supplemental Table 1), only 40 had homologs on the mouse genome (indicated in red on Figs. 1 and 2) and two were not mappable due to outdated location markers that could not be standardized.

**Figure 1.**
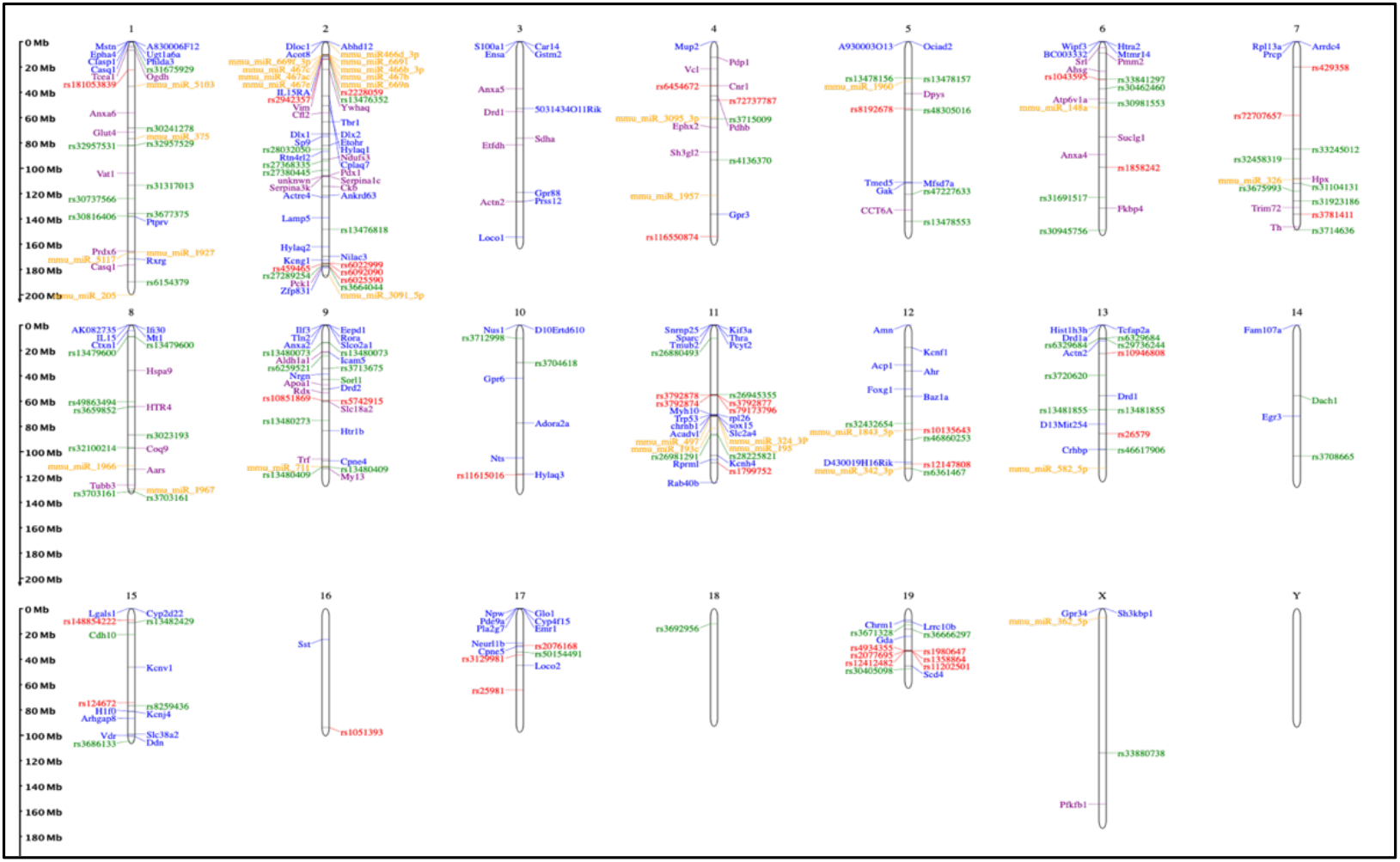
Chromosome Map for Omics Data Associated With/Causal for Physical Activity. Color indicates the type of “omic” data; with green- genomic, blue – transcriptomic, purple – proteomic, yellow – miRNA, red – homologous human genomic variant.

**Figure 2.**
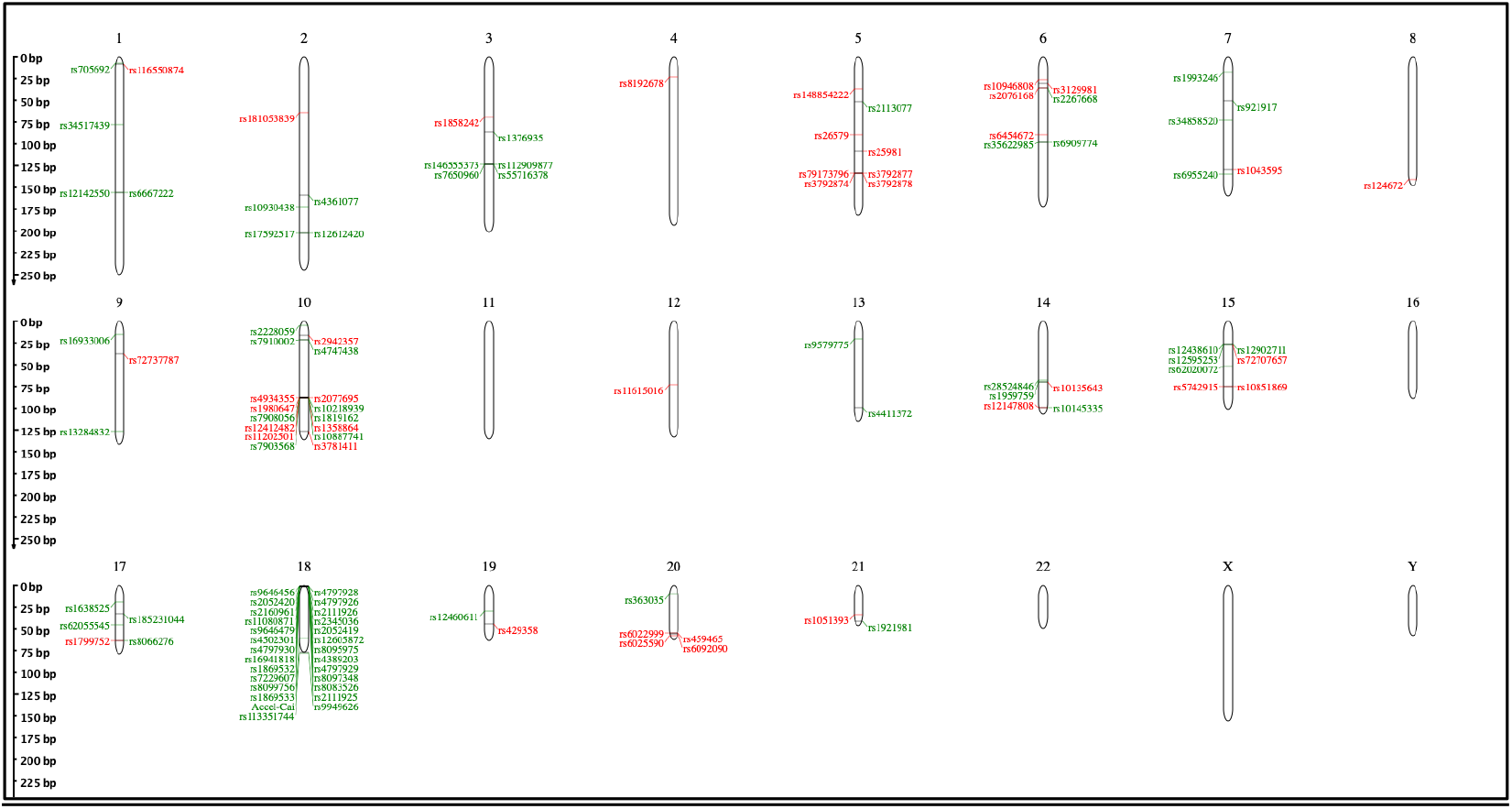
Map of Genetic Molecules Associated with Physical Activity - Human Genome

#### Potential Upstream Regulators

As detailed in the methods, subsets of data were created for brain and muscle. Not all the molecules from our brain or muscle datasets corresponded to molecules in the IPA database. Of the 366 molecules in the brain and 345 molecules in the muscle dataset, 226 (61.75%) and 195 molecules (56.52%) corresponded to molecules in the IPA database for brain and muscle respectively (Supplemental Table 2). The upstream regulator analysis (Table 2; Suppl Tables 3 and 4) revealed that β-estradiol was the highest predicted regulator for both data sets, with 44 molecules in the brain dataset affected (p=0.000000048) and 41 molecules in the muscle dataset (p=0.00000000179). This finding indicates that 19.5% of the brain data and 21% of the muscle data are potentially regulated upstream by β-estradiol. The other predicted regulators in the top five for both sets (Table 2) were also similar (e.g. TP53 ranked 5th in brain and 2nd in muscle), varying primarily in the number of molecules affected. Also unique to these data were the potential for exogenous chemical drugs – lipopolysaccharide (2^nd^ brain, 30 molecules / 3^rd^ muscle, 32 molecules) and dexamethasone (3^rd^ brain, 27 molecules / 5^th^ muscle, 27 molecules) -as potential regulators of physical activity. The amount of overlap between the two datasets (muscle vs brain subsets) was not surprising due to the overlap of data points discussed previously in the methods.

**Table 2.**
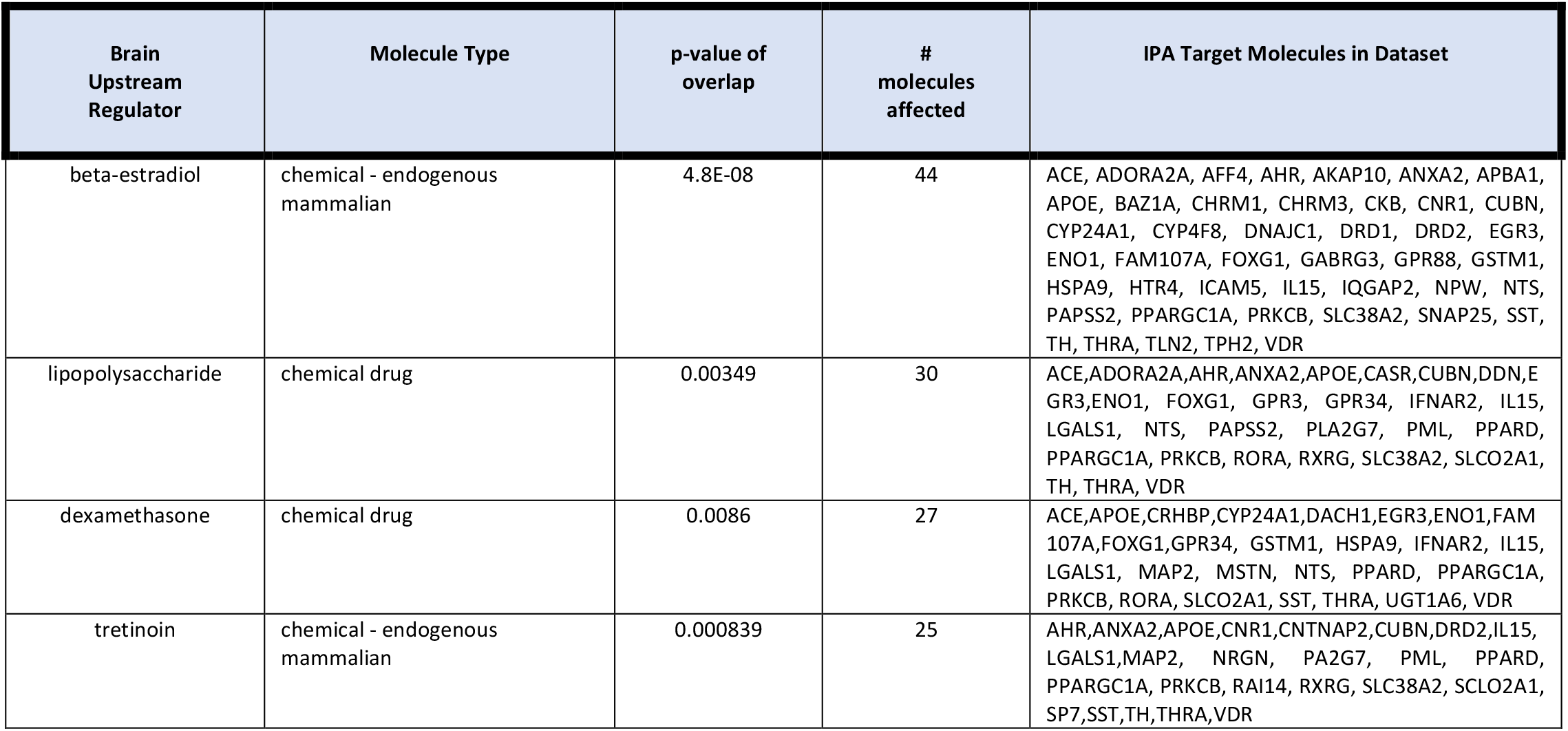
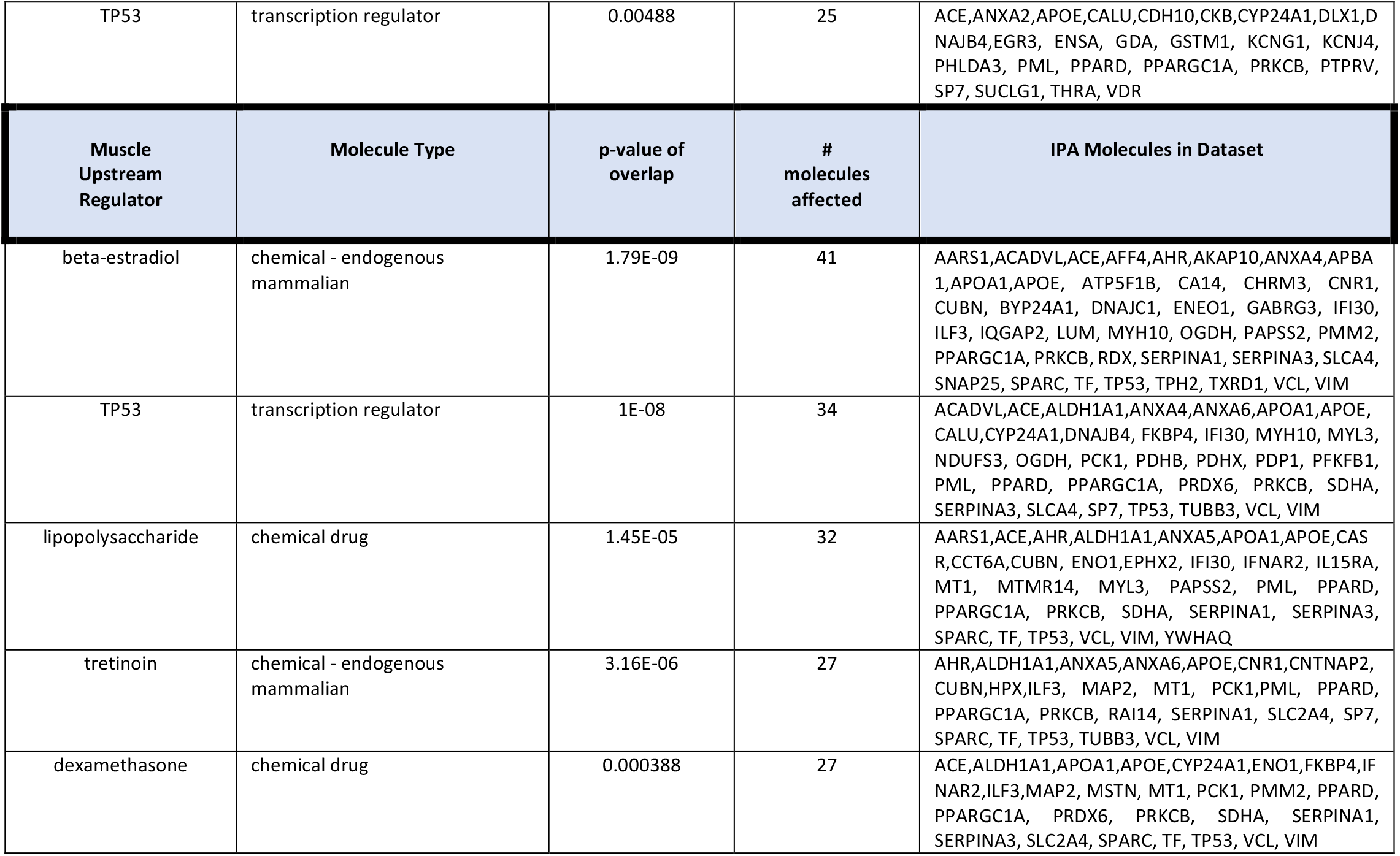
Top 5 Upstream Regulators for Brain and Muscle Subsets

## Discussion

The greatest number of genetic molecules in the mouse genome reside on Chrms 2, 1, 16, 9, 11, and 13 (n= 61, 36, 36, 34, 33, and 22, respectively, Fig. 1). When standardized for length of chromosome, Chrms. 8, 15, 7, 18, and Y (n=20, 15, 20, 1, 1, respectively), have the highest density of genetic molecules (Fig. 1). However, given the multi-chromosomal interaction of some of the molecules (e.g., miRNA) as well as the unmapped epistatic molecules^41^, it is unclear whether chromosomal density of these molecules is important. Interestingly, there were 74 genetic variants identified in humans that did not have homologous molecules in the mouse model, the majority on Chrm 18 (human). A major source of these molecules is De Moor and colleague’s findings^44^ that were associated with exercise participation. These non-homologous molecules may represent specific human-related mechanisms that regulate physical activity. To date, Kostrzewa, et al. is the only cross-species translation attempt in this literature^39^. Certainly, cross-species translation is an area that needs further investigation to determine if there are pathways specific to either the animal or human genetic control of physical activity (Fig. 2).

There are a variety of potential upstream regulators for the molecules collected in this dataset. Interestingly, three of the top five for both brain and muscle are nuclear receptor binding ligands; estradiol (estrogen receptor), dexamethasone (glucocorticoid receptor), and tretinoin (retinoic acid receptor), indicating a potential role of nuclear receptors in the regulation of physical activity. Unlike most intercellular messengers, these ligands can cross the plasma membrane and directly interact with nuclear receptors inside the cell verses being limited to cell surface interactions^70^. Activation of nuclear receptors subsequently causes transcription of genes controlling a variety of cellular processes like cell proliferation, development, metabolism, and reproduction; additionally, some nuclear receptors have also been found to regulate cellular functions within the cytoplasm (e.g. estrogens)^71^. Selective nuclear receptor modulation may be an area of interest for future studies.

The top ranked upstream regulator for both muscle and brain datasets identified in this review was β-estradiol. Since initial efforts in the 1920’s^20^, it has been well established that sex steroids – most prominently estrogen^19,21^, and/or testosterone^17,18^, markedly alter physical activity levels. For example, removal of endogenous sex steroids decreased wheel-running by 90% in male animals and 69% in female animals, with administration of exogenous testosterone recovering wheel running activity (103% of baseline in females, 90% of baseline in males)^18^. However, what has not been clear is the mechanism through which these sex steroids work; i.e., is the level of sex steroids – and thus activity – controlled by genetics, or is the effect of sex steroids merely as a ‘biological influencer’ [a detailed review of the mechanistic differences exists by Bjornstrom et al^72^]. There is evidential support for the capacity of estradiol to increase physical activity levels; however, the mechanisms directly linking estradiol to increased physical activity have not been tested. For example, both testosterone and 17β-estradiol have been shown to enhance insulin-stimulated glucose uptake in female myotubes and to increase palmitate oxidation in both male and female myotubes^73^ suggesting a role for estradiol in regulating substrate availability in the muscle. Further, there is extensive literature^74^ showing that expression of *Slc2a4* – also known as *Glut4* – is affected by estrogen triggered-mechanisms. Tsao, et al.^9^ clearly showed that overexpression of *Glut4* increased wheel-running activity in mice, suggesting that any mechanism through which *Glut4* expression was increased, would like-wise increase physical activity levels. Muscle biopsies obtained from men supplemented with 17β-estradiol demonstrated elevated medium-chain acyl-CoA dehydrogenase (MCAD), PGC1α mRNA, and reduced micro-mRNA, miR-29b (predicted to regulate PGC-1α) supporting enhanced fat oxidation during exercise^75^. Supporting central mechanisms of estradiol; Peterson, et al. demonstrated estradiol impacts behavioral and synaptic correlates of addiction in female rats through activation of cannabinoid 1 receptor (CB1R -CNR1 gene)^76^. Dubreucq, et al. has demonstrated decreased dopamine activity and subsequent decreased wheel running in CB1R deficient mice^16^ supporting a link between estradiol, cannabinoid receptors, and dopamine levels. In short, given the large number of targeted muscle molecules that are regulated in some manner by β-estradiol in both the brain and muscle, it is probable that β-estradiol exerts its physical activity regulating effects through both peripheral and central mechanisms.

Another potential regulator, TP53, a nuclear transcription regulator – was 2^nd^ and 5^th^ ranked potential upstream regulator of the muscle and brain molecules, respectively. TP53 is most often classified as a tumor suppressor gene and is activated by a variety of cellular stresses including oxidative stress at which time it becomes a transcription regulator (40). It has recently been suggested that moderate intensity exercise stimulates the cancer protective functions of TP53 (41). Thus, it is likely that physical activity can cause TP53 to accumulate in the stressed cell and regulate the various target genes in the physical activity dataset. However, since TP53 is not upregulated until physical activity is undertaken, TP53’s effects would be a product of exercise, rather than being a regulator of physical activity.

## Limitations

There are several limitations that must be considered when interpreting the results of this systematic review. First, while we worked to be as inclusive as possible, it is probable that there are studies that are missing from this summary. Certainly, there is no one set of key terms used by all the authors in this field; some agreed-upon set of terms would help similar future efforts.

Second, during the time-span of the considered papers, ranging from publication in the early 2000s to 2020, the methodology employed in genetic investigations changed dramatically. One significant example is the switch from using microsatellites to SNP as genomic markers. While we made the best efforts to standardize measurements, such as using the most recent builds (at time of data analysis) of the murine and human genomes, there were some results that could not be standardized^42^ and thus, were left out of the analysis. In a similar vein, the minimal epistatic data that are available^41,45^ was left out of the analysis primarily because of uncertainty of how to classify these data within the parameters of the network analysis. Removing these data from the analysis should not be construed to indicate they are not important – indeed the epistatic data are the only available data of their kind in the field – but rather are examples of the limits of our analysis paradigm.

Third, a significant limitation both in the composing of and interpretation of the analyses is that there is a tremendous amount of data available. While we recognize our limits for interpretation and the length limitations of a written study, we have provided not only the data-files used in the network analyses as supplemental information, but also, extensive supplementary tables to enable other investigators to add to the dataset and rerun the analysis, or to gather the interpretative-narrative in their own work. Fourth, it must be remembered that like much work in the genetic realm, the results are mostly associative with few causal studies available. While some of the data used in this study were causative^9,10,15,16,53^, much of the data are associative, especially the genomic variants we identified. Given there have been past difficulties noted in the literature in regards to defining mechanisms based on associative findings – especially genomic variant designs^77^ – inferences based on genomic variant data should be made cautiously. Further, a continuing limitation of designs focusing on genomic variation is the uncertainty of whether these variants translate into different functional molecules. The addition of proteomic, transcript, and microRNA data compliments the genomic variation data and can add support for those genomic variants. If anything, our results guide future studies that focus on causative mechanisms.

Last, the work is limited by the tools available. Even though tools such as IPA are powerful, these tools are still limited in their scope and the genetic molecules available in their databases. For example, between 40-45% of the target molecules identified from the literature are not identifiable in the IPA database and so it is possible that other critical networks have been missed in this report. Additionally, given the magnitude of potential targets of each miRNA, it is possible that IPA does not contain full network linkages for all the miRNAs in the dataset and as such may not be the best approach to investigating the potential role of miRNAs in regulating physical activity.

In conclusion, the current paper describes our approach to gather the significant physical activity genetic regulating molecules from the literature, and to integrate all the known possible data into common upstream regulators related to physical activity control. Given the established consensus that physical activity is regulated both at the central and peripheral levels, potential upstream regulators are presented for both central (e.g., brain) and peripheral (e.g., muscle) molecules. Based on these analyses, the upstream regulator analyses may be a great tool for future researchers to build research questions and investigate the broader scope of mechanisms directly linking upstream regulators to regulation of physical activity.

## Supporting information

Supplemental Table 1: All molecules associated with / causative of physical activity.

Supplemental Table 2: Mapped molecules in database.

Supplemental Table 3: All upstream regulators of brain molecules

Supplemental Table 4: All upstream regulators of muscle molecules

Supplemental Content Data File 1: All molecules associated with / causative of physical activity in the brain

Supplemental Content Data File 2: All molecules associated with / causative of physical activity in the muscle

## Acknowledgements

Funding for this study was provided by the Debbie and Mike Hilliard endowment. This research was supported [in part] by the Intramural Research Program of the NIH, National Institute of Environmental Health Sciences. We’d like to thank Dr. David Threadgill and the Texas A&M Center for Environmental Health Research in conjunction with the Texas A&M Vice-President for Research Office for providing access to Qiagen’s Integrated Pathway Analysis (IPA). We want to strongly recognize Dr. David Ferguson’s coining of the phrase ‘biological influencer’ and thank him for allowing us to use the phrase. We thank Dr. Ayland Letsinger for making the introductions that provided initial access to IPA and advice and guidance on its use by Drs. K. Gerrish and L. Liu of the Molecular Genomics Core Laboratory of the National Institute of Environmental Health Sciences. Lastly, the results of the current study do not constitute endorsement by the American College of Sports Medicine.

## Supplemental Content

Supplemental Table 1: All molecules associated with / causative of physical activity.

Supplemental Table 2: Mapped molecules in database.

Supplemental Table 3: All upstream regulators of brain molecules

Supplemental Table 4: All upstream regulators of muscle molecules

## Data Files

Supplemental Content Data File 1: All molecules associated with / causative of physical activity in the brain (.csv).

Supplemental Content Data File 2: All molecules associated with / causative of physical activity in the muscle (.csv).

