## Supplemental Table 1: All molecules associated with / causative of physical activity. for "“It All Rolls Downstream: Upstream Control of Physical Activity Regulation”"

**Supplemental Table 1: All Molecules Associated with / causative of Physical Activity in Current Literature**

| **Study ID #** | **Study** | **Molecule Type** | **Name / ID** | **Associated Gene (where applicable)** | **Tissue sub-analysis** | **Physical Activity Phenotype** | **P-Value** | **Chrm** | **Location (mbp)** | **Mouse Homolog Chrm (where available)** | **Mouse Homolog Location (where available)** |
| --- | --- | --- | --- | --- | --- | --- | --- | --- | --- | --- | --- |
| 1 | Ferguson 2014 | Protein | Calsequestrin 1 | Casq1 | Muscle | High Physical Activity | 0.000025 | 1 | 172.209894-172.219868 |  |  |
| 2 | Ferguson 2014 | Protein | Peroxiredoxin-6 | Prdx6 | Muscle | Low Physical Activity | 0.00079 | 1 | 161.240112-161.251210 |  |  |
| 3 | Ferguson 2015 | Protein | Transcription elongation factor A | Tcea1 | Brain | Low Physical Activity | 0.0078 | 1 | 4.857814-4.897909 |  |  |
| 4 | Ferguson 2014 | Protein | Phosphoenolpyruvate carboxykinase | Pck1 | Muscle | Low Physical Activity | 0.00042 | 2 | 173.153073-173.159274 |  |  |
| 5 | Ferguson 2014 | Protein | Vimentin | Vim | Muscle | Low Physical Activity | 0.007 | 2 | 13.574311-13.582826 |  |  |
| 6 | Ferguson 2014 | Protein | NADH dehydrogenase [ubiquinone] iron-sulfur protein 3 | Ndufs3 | Muscle | Low Physical Activity | 0.027 | 2 | 90.894627-90.904721 |  |  |
| 7 | Ferguson 2014 | Protein | Pyruvate dehydrogenase protein X component | Pdx1 | Muscle | High Physical Activity | 0.029 | 2 | 103.021075-103.073513 |  |  |
| 8 | Ferguson 2014 | Protein | Annexin A5 | Anxa5 | Muscle | Low Physical Activity | 0.036 | 3 | 36.448923-36.475887 |  |  |
| 9 | Ferguson 2014 | Protein | Electron transfer flavoprotein-ubiquinone oxidoreductase | Etfdh | Muscle | Low Physical Activity | 0.0016 | 3 | 79.603788-79.628767 |  |  |
| 10 | Ferguson 2014 | Protein | [Pyruvate dehydrogenase [acetyl-transferring]]-phosphatase 1 | Pdp1 | Muscle | High Physical Activity | 0.0022 | 4 | 11.958183-11.966450 |  |  |
| 11 | Ferguson 2015 | Protein | Endophilin | Sh3gl2 | Brain | High Physical Activity | 0.032 | 4 | 85.205456-85.389380 |  |  |
| 12 | Ferguson 2014 | Protein | T-complex protein 1 subunit zeta | CCT6A | Muscle | Low Physical Activity | 0.035 | 5 | 129.787356-129.846443 |  |  |
| 13 | Ferguson 2014 | Protein | Annexin A4 | Anxa4 | Muscle | High Physical Activity | 0.023 | 6 | 86.736840-86.793584 |  |  |
| 14 | Ferguson 2014 | Protein | FK506 binding protein 4 | Fkbp4 | Muscle | High Physical Activity | 0.029 | 6 | 128.430103-128.438656 |  |  |
| 15 | Ferguson 2015 | Protein | Succinyl-CoA ligase | Suclg1 | Brain | High Physical Activity | 0.032 | 6 | 73.248505-73.276907 |  |  |
| 16 | Ferguson 2014 | Protein | Hemopexin | Hpx | Muscle | Low Physical Activity | 0.018 | 7 | 105.591611-105.600116 |  |  |
| 17 | Ferguson 2014 | Protein | Tripartite motif-containing protein 72 | Trim72 | Muscle | High Physical Activity | 0.029 | 7 | 128.003949-128.011033 |  |  |
| 18 | Knab 2009 | Protein | Tyrosine hydrosylase | Th | Brain | High Physical Activity | 0.0008 | 7 | 142.892779-142.899995 |  |  |
| 19 | Ferguson 2014 | Protein | Tubulin beta-3 | Tubb3 | Muscle | Low Physical Activity | 0.007 | 8 | 123.411553-123.422015 |  |  |
| 20 | Ferguson 2014 | Protein | Alanyl-tRNA synthase | Aars | Muscle | Low Physical Activity | 0.031 | 8 | 111.033144-111.057664 |  |  |
| 21 | Ferguson 2014 | Protein | Ubiquinone biosynthesis protein COQ9 | Coq9 | Muscle | High Physical Activity | 0.042 | 8 | 94.838413-94.854895 |  |  |
| 22 | Ferguson 2014 | Protein | Radixin | Rdx | Muscle | Low Physical Activity | 0.00028 | 9 | 52.047150-52.088738 |  |  |
| 23 | Ferguson 2014 | Protein | Transferrin | Trf | Muscle | Low Physical Activity | 0.0026 | 9 | 103.208876-103.230286 |  |  |
| 24 | Ferguson 2014 | Protein | Myosin, light polypeptide 3 | My13 | Muscle | High Physical Activity | 0.0062 | 9 | 110.763678-110.769802 |  |  |
| 25 | Ferguson 2014 | Protein | Apolipoprotein A-I | Apoa1 | Muscle | High Physical Activity | 0.0062 | 9 | 46.228630-46.230469 |  |  |
| 26 | Ferguson 2014 | Protein | Apolipoprotein A-I | Apoa1 | Muscle | High Physical Activity | 0.0046 | 9 | 46.228630-46.230469 |  |  |
| 27 | Ferguson 2014 | Protein | Lumican | Lum | Muscle | High Physical Activity | 0.004 | 10 | 97.565501-97.572703 |  |  |
| 28 | Ferguson 2014 | Protein | Thioredoxin reductase 1 | Txnrd1 | Muscle | High Physical Activity | 0.0051 | 10 | 82.833951-82.897712 |  |  |
| 29 | Ferguson 2014 | Protein | Atp5b | Atp5b | Muscle | Low Physical Activity | 0.007 | 10 | 128.083307-128.090388 |  |  |
| 30 | Ferguson 2014 | Protein | Annexin A6 | Anxa6 | Muscle | High Physical Activity | 0.00031 | 11 | 54.979108-55.033445 |  |  |
| 31 | Ferguson 2014 | Protein | Synaptic vesicle membrane protein VAT-1 homolog | Vat1 | Muscle | High Physical Activity | 0.0046 | 11 | 101.458748-101.466199 |  |  |
| 32 | Ferguson 2014 | Protein | 2-Oxoglutarate dehydrogenase | Ogdh | Muscle | Low Physical Activity | 0.035 | 11 | 6.291633-6.356642 |  |  |
| 33 | Ferguson 2014 | Protein | Annexin A6 | Anxa6 | Muscle | High Physical Activity | 0.0016 | 11 | 54.979108-55.033445 |  |  |
| 34 | Tsao 2001 | Protein | Slc2a4 | Glut4 | Muscle | High Physical Activity | <0.05 | 11 | 69.942539-69.948188 |  |  |
| 35 | Ferguson 2014 | Protein | Serine protease inhibitor A3K | Serpina3k | Muscle | High Physical Activity | 0.005 | 12 | 104.338486-104.345739 |  |  |
| 36 | Ferguson 2014 | Protein | Cofilin2 | Cfl2 | Muscle | Low Physical Activity | 0.027 | 12 | 54.858815-54.862877 |  |  |
| 37 | Ferguson 2014 | Protein | Alpha-1-antitrypsin 1-4 |  | Muscle | Low Physical Activity |  | 12 | 103.763594-103.773592 |  |  |
| 38 | Ferguson 2014 | Protein | Serpina1c protein | Serpina1c | Muscle | Low Physical Activity | 0.031 | 12 | 103.894926-103.904887 |  |  |
| 39 | Ferguson 2014 | Protein | 14-3-3 Protein gamma subtype | Ywhaq | Muscle | High Physical Activity | 0.036 | 12 | 21.390329-21.417436 |  |  |
| 40 | Ferguson 2015 | Protein | Cluster of creatine kinase B | Ckb | Brain | High Physical Activity | 0.032 | 12 | 111.669361-111.672338 |  |  |
| 41 | Ferguson 2014 | Protein | Sdha | Sdha | Muscle | Low Physical Activity | 0.0016 | 13 | 74.322255-74.350240 |  |  |
| 42 | Ferguson 2014 | Protein | Alpha-actinin-2 | Actn2 | Muscle | Low Physical Activity | 0.0044 | 13 | 122.69426-123.40732 |  |  |
| 43 | Knab 2009 | Protein | Drd1 | Drd1 | Brain | High Physical Activity | 0.0001 | 13 | 54.051183-54.055658 |  |  |
| 44 | Ferguson 2014 | Protein | Epoxide hydrolase 2 | Ephx2 | Muscle | High Physical Activity | 0.0051 | 14 | 66.084374-66.124500 |  |  |
| 45 | Ferguson 2014 | Protein | Pyruvate dehydrogenase E1 component subunit beta | Pdhb | Muscle | High Physical Activity | 0.023 | 14 | 81.65996-81.730250 |  |  |
| 46 | Ferguson 2014 | Protein | Vinculin | Vcl | Muscle | Low Physical Activity | 0.028 | 14 | 20.929398-21.033676 |  |  |
| 47 | Ferguson 2015 | Protein | Dihydropryminidinase | Dpys | Brain | Low Physical Activity | 0.018 | 15 | 39.768487-39.857470 |  |  |
| 48 | Ferguson 2014 | Protein | Phosphomannomutase 2 | Pmm2 | Muscle | High Physical Activity | 0.018 | 16 | 8.637613-8.657608 |  |  |
| 49 | Ferguson 2014 | Protein | Alpha-2-HS-glycoprotein | Ahsg | Muscle | Low Physical Activity | 0.026 | 16 | 22.892015-22.899451 |  |  |
| 50 | Ferguson 2014 | Protein | Sarcalumenin | Srl | Muscle | High Physical Activity | 0.029 | 16 | 4.480216-4.541816 |  |  |
| 51 | Ferguson 2014 | Protein | V-type proton ATPase catalytic subunit A | Atp6v1a | Muscle | High Physical Activity | 0.0016 | 16 | 44.085402-44.139702 |  |  |
| 52 | Ferguson 2015 | Protein | V type proton ATPase catalytic subunit A | Atp6v1a | Brain | Low Physical Activity | 0.018 | 16 | 44.085402-44.139702 |  |  |
| 53 | Ferguson 2015 | Protein | Cluster of stress 70 protein (mitochondrial) | Hspa9 | Brain | Low Physical Activity | 0.018 | 18 | 34.937414-34.954351 |  |  |
| 54 | Ferguson 2014 | Protein | Aldehyde dehydrogenase 1 | Aldh1a1 | Muscle | Low Physical Activity | 0.031 | 19 | 20.492715-20.643465 |  |  |
| 55 | Ferguson 2013 | Protein | Vmat2 | Slc18a2 | Muscle | High Physical Activity | 0.0016 | 19 | 59.260896-59.296012 |  |  |
| 56 | Ferguson 2014 | Protein | 6-Phosphofructo-2-kinase/fructose-2,6-bisphosphatase 1 | Pfkfb1 | Muscle | Low Physical Activity | 0.031 | X | 150.588230-150.643878 |  |  |
| 57 | Dubreucq 2013 | Protein | cannabinoid receptor 1 | Cnr1 | Brain | High Physical Activity | <0.01 | 4 | 33.924593-33.948831 |  |  |
| 58 | Pistilli 2011 | Protein | interleukin 15 receptor, alpha chain | IL15RA | Muscle | high physical avitivy | <0.05 | 2 | 11.709992-11.738796 |  |  |
| 59 | Jean 2012 | Protein | 5 hydroxytryptamine (serotonin) receptor 4 | HTR4 | Brain | high physical activty | <0.05 | 18 | 62.457275-62.629648 |  |  |
| 60 | Kelly 2012 | Transcript | Clasp1 | Clasp1 | Brain | Running Time | <0.05 | 1 | 118.389058-118.612678 |  |  |
| 61 | Kelly 2012 | Transcript | Phlda3 | Phlda3 | Brain | Running Time | <0.05 | 1 | 135.766119-135.769136 |  |  |
| 62 | Kelly 2012 | Transcript | Ugt1a6a | Ugt1a6a | Brain | Average Running Speed | <0.05 | 1 | 88.134809-88.218997 |  |  |
| 63 | Kelly 2012 | Transcript | A830006F12 | A830006F12 | Brain | Maximum Running Speed | <0.05 | 1 | 70.725715-70.885397 |  |  |
| 64 | Kelly 2012 | Transcript | Abhd12 | Abhd12 | Brain | Running Distance | <0.05 | 2 | 150.832493-150.904741 |  |  |
| 65 | Kelly 2014 | Transcript | Acot8 | Acot8 | Muscle | Running Distance | <0.05 | 2 | 164.792765-164.804882 |  |  |
| 66 | Kelly 2012 | Transcript | Ensa | Ensa | Brain | Running Distance | <0.05 | 3 | 95.624993-95.632102 |  |  |
| 67 | Kelly 2012 | Transcript | Gstm2 | Gstm2 | Brain | Maximum Running Speed | <0.05 | 3 | 107.981702-107.986453 |  |  |
| 68 | Kelly 2014 | Transcript | Car14 | Car14 | Muscle | Running Distance and Average Running Speed | <0.05 | 3 | 95.897768-95.904691 |  |  |
| 69 | Kelly 2014 | Transcript | S100a1 | S100a1 | Muscle | Average Running Speed | <0.05 | 3 | 90.511034-90.514392 |  |  |
| 70 | Kelly 2012 | Transcript | Fam107a | Fam107a | Brain | Running Distance/Average Running Speed/ Running Time | <0.05 | 14 | 8.296274-8.318023 |  |  |
| 71 | Kelly 2014 | Transcript | Mup2 | Mup2 | Muscle | Running Time | <0.05 | 4 | 60.135932-60.154289 |  |  |
| 72 | Kelly 2012 | Transcript | A930003O13 | A930003O13 | Brain | Running Distance | <0.05 | 5 | 22.73891-22.746907 |  |  |
| 73 | Kelly 2012 | Transcript | Ociad2 | Ociad2 | Brain | Maximum Running Speed | <0.05 | 5 | NA |  |  |
| 74 | Kelly 2012 | Transcript | Wipf3 | Wipf3 | Brain | Average Running Speed | <0.05 | 6 | 54.429603-54.503768 |  |  |
| 75 | Kelly 2012 | Transcript | Htra2 | Htra2 | Brain | Maximum Running Speed | <0.05 | 6 | 83.051266-83.055273 |  |  |
| 76 | Kelly 2012 | Transcript | BC003332 | BC003332 | Brain | Maximum Running Speed | <0.05 | 6 | NA |  |  |
| 77 | Kelly 2014 | Transcript | Mtmr14 | Mtmr14 | Muscle | Running Time | <0.05 | 6 | NA |  |  |
| 78 | Kelly 2012 | Transcript | Prcp | Prcp | Brain | Running Distance and Duration | <0.05 | 7 | 92.87447-92.934583 |  |  |
| 79 | Kelly 2012 | Transcript | Arrdc4 | Arrdc4 | Brain | Running Duration | <0.05 | 7 | 68.736995-68.749241 |  |  |
| 80 | Kelly 2014 | Transcript | Rpl13a | Rpl13a | Muscle | Running Time | <0.05 | 7 | 45.125558-45.128761 |  |  |
| 81 | Kelly 2014 | Transcript | Prcp | Prcp | Muscle | Running Distance and Duration | <0.05 | 7 | 92.87447-92.934583 |  |  |
| 82 | Kelly 2012 | Transcript | IL15 | IL15 | Brain | Running Duration | <0.05 | 8 | 82.331632-82.403222 |  |  |
| 83 | Kelly 2014 | Transcript | Mt1 | Mt1 | Muscle | Running Distance and Average Running Speed | <0.05 | 8 | 94.179082-94.180327 |  |  |
| 84 | Kelly 2014 | Transcript | Ifi30 | Ifi30 | Muscle | Average Running Speed | <0.05 | 8 | 70.762774-70.766663 |  |  |
| 85 | Kelly 2014 | Transcript | AK082735 | AK082735 | Muscle | Maximum Running Speed | <0.05 | 8 | 69.132669-69.184225 |  |  |
| 86 | Kelly 2012 | Transcript | Slco2a1 | Slco2a1 | Brain | Running Distance/Average Running Speed | <0.05 | 9 | 102.988712-103.096002 |  |  |
| 87 | Kelly 2012 | Transcript | Tln2 | Tln2 | Brain | Running Distance | <0.05 | 9 | 67.217087-67.559703 |  |  |
| 88 | Kelly 2012 | Transcript | Anxa2 | Anxa2 | Brain | Maximum Running Speed | <0.05 | 9 | 69.45362-69.491795 |  |  |
| 89 | Kelly 2014 | Transcript | Eepd1 | Eepd1 | Muscle | Running Time | <0.05 | 9 | 25.481547-25.60411 |  |  |
| 90 | Kelly 2014 | Transcript | Ilf3 | Ilf3 | Muscle | Running Time | <0.05 | 9 | 21.367871-21.405361 |  |  |
| 91 | Kelly 2012 | Transcript | D10Ertd610 | D10Ertd610 | Brain | Maximum Running Speed | <0.05 | 10 | 127.182525-127.190083 |  |  |
| 92 | Kelly 2014 | Transcript | Nus1 | Nus1 | Muscle | Running Distance | <0.05 | 10 | 52.417547-52.440183 |  |  |
| 93 | Kelly 2012 | Transcript | Snrnp25 | Snrnp25 | Brain | Running Distance | <0.05 | 11 | 32.205415-32.208984 |  |  |
| 94 | Kelly 2012 | Transcript | Tmub2 | Tmub2 | Brain | Average Running Speed | <0.05 | 11 | 102.284931-102.289237 |  |  |
| 95 | Kelly 2012 | Transcript | Thra | Thra | Brain | Maximum Running Speed | <0.05 | 11 | 98.740638-98.769006 |  |  |
| 96 | Kelly 2012 | Transcript | Kif3a | Kif3a | Brain | Maximum Running Speed | <0.05 | 11 | 53.567379-53.601967 |  |  |
| 97 | Kelly 2012 | Transcript | Tmub2 | Tmub2 | Brain | Maximum Running Speed | <0.05 | 11 | 102.284931-102.289237 |  |  |
| 98 | Kelly 2014 | Transcript | Pcyt2 | Pcyt2 | Muscle | Maximum Running Speed | <0.05 | 11 | 120.610087-120.617936 |  |  |
| 99 | Kelly 2014 | Transcript | Sparc | Sparc | Muscle | max running speed | <0.05 | 11 | 55.3945-55.423183 |  |  |
| 100 | Kelly 2012 | Transcript | Amn | Amn | Brain | Maximum Running Speed | <0.05 | 12 | 111.271095-111.276426 |  |  |
| 101 | Kelly 2012 | Transcript | Hist1h3h | Hist1h3h | Brain | Running Distance/Running Time | <0.05 | 13 | 1.717659-1.718069 |  |  |
| 102 | Kelly 2012 | Transcript | Lgals1 | Lgals1 | Brain | Average Running Speed | <0.05 | 15 | - |  |  |
| 103 | Kelly 2012 | Transcript | Cyp2d22 | Cyp2d22 | Brain | Maximum Running Speed | <0.05 | 15 | 82.370527-82.38026 |  |  |
| 104 | Kelly 2012 | Transcript | Emr1 | Emr1 | Brain | Average Running Speed | <0.05 | 17 | 57.358691-57.483527 |  |  |
| 105 | Kelly 2012 | Transcript | Gpr34 | Gpr34 | Brain | Running Time | <0.05 | X | 13.632089-13.640858 |  |  |
| 106 | Kelly 2014 | Transcript | Sh3kbp1 | Sh3kbp1 | Muscle | Running Distance/ Average Running Speed/ Maximum Running Speed | <0.05 | X | 159.627272-159.978069 |  |  |
| 107 | Kelly 2012 | Transcript | DBY (Ddx3y) | DBY (Ddx3y) | Brain | Running Distance/ Running Time/ Maximum Running Speed | <0.05 | Y | 1.260771-1.286629 |  |  |
| 108 | Williams 2014 | Transcript | Acp1 | Acp1 | Brain/muscle | Lcomotion/Rearing | <0.05 | 12 | 30.893326-30.911589 |  |  |
| 109 | Williams 2014 | Transcript | Ahr | Ahr | Brain/muscle | Lcomotion/Rearing | <0.05 | 12 | 35.497979-35.534989 |  |  |
| 110 | Yang 2012 | Transcript | Tcfap2a | Tcfap2a | Brain | Wheel running and cage activity | <0.05 | 13 | 40.715302-40.738376 |  |  |
| 111 | Yang 2012 | Transcript | Drd1a | Drd1a | Brain | Wheel running and cage activity | <0.05 | 13 | 54.053494-54.053596 |  |  |
| 112 | Dawes 2014 | Transcript | Casq1 | Casq1 | Brain/muscle | Running Distance/Time/Speed | <0.05 | 1 | 172.209894-172.219868 |  |  |
| 113 | Dawes 2014 | Transcript | Mstn | Mstn | Brain/muscle | Running Distance/Time/Speed | <0.05 | 1 | 53.06164-53.068079 |  |  |
| 114 | Eisener-Dorman 2010 | Transcript | Rora | Rora | Brain | Open-field distance, ambulations, rearing, velocity | <0.05 | 9 | 68.653786-69.388246 |  |  |
| 115 | Kas 2009 | Transcript | Fam124b (aka A830043J08Rik) | Fam124b (aka A830043J08Rik) | Brain | Sheltering bheavior, motor activity and consumption attempts | <0.05 | 1 | 80.198706-80.218473 |  |  |
| 116 | Kas 2009 | Transcript | Epha4 | Epha4 | Brain | Sheltering bheavior, motor activity and consumption attempts | <0.05 | 1 | 77.367185-77.515088 |  |  |
| 117 | Kumar 2010 | Transcript | Npw | Npw | Brain | Ambulations, wheel running distance/time/peak speed | <0.05 | 17 | 24.65733-24.658457 |  |  |
| 118 | Kumar 2010 | Transcript | Glo1 | Glo1 | Brain | Ambulations, wheel running distance/time/peak speed | <0.05 | 17 | 30.584936-30.626871 |  |  |
| 119 | Kumar 2010 | Transcript | Cyp4f15 | Cyp4f15 | Brain | Ambulations, wheel running distance/time/peak speed | <0.05 | 17 | 32.685627-32.703352 |  |  |
| 120 | Kumar 2010 | Transcript | Pla2g7 | Pla2g7 | Brain | Ambulations, wheel running distance/time/peak speed | <0.05 | 17 | 43.568098-43.612201 |  |  |
| 121 | Kumar 2010 | Transcript | Pde9a | Pde9a | Brain | Ambulations, wheel running distance/time/peak speed | <0.05 | 17 | 31.38621-31.47631 |  |  |
| 122 | Umemori 2009 | Transcript | Hylaq1 | Hylaq1 | Brain/muscle | Average ambulation activity | <0.05 | 2 | 84.013198-14.866747 |  |  |
| 123 | Umemori 2009 | Transcript | Hylaq2 | Hylaq2 | Brain/muscle | Active time | <0.05 | 2 | 151.640941-164.119146 |  |  |
| 124 | Umemori 2009 | Transcript | Hylaq3 | Hylaq3 | Brain/muscle | Total home-cage ambulations (average activity + active time) | <0.05 | 10 | 111.568226-117.986649 |  |  |
| 125 | Umemori 2009 | Transcript | Actre2 | Actre2 | Brain/muscle | Average ambulation activity | <0.05 | 2 | 80.005954-80.006113 |  |  |
| 126 | Umemori 2009 | Transcript | Etohr | Etohr | Brain/muscle | Average ambulation activity | <0.05 | 2 | 80.005954-80.006113 |  |  |
| 127 | Umemori 2009 | Transcript | Cplaq7 | Cplaq7 | Brain/muscle | Average ambulation activity | <0.05 | 2 | 93.421483--9.342165 |  |  |
| 128 | Umemori 2009 | Transcript | Actre3 | Actre3 | Brain/muscle | Average ambulation activity | <0.05 | 2 | 112.024987-128.173821 |  |  |
| 129 | Umemori 2009 | Transcript | Actre4 | Actre4 | Brain/muscle | Average ambulation activity | <0.05 | 2 | 112.024987-128.173821 |  |  |
| 130 | Umemori 2009 | Transcript | Rrodp2 | Rrodp2 | Brain/muscle | Active time | <0.05 | 2 | - |  |  |
| 131 | Umemori 2009 | Transcript | Slms2 | Slms2 | Brain/muscle | Active time | <0.05 | 2 | - |  |  |
| 132 | Umemori 2009 | Transcript | Nilac3 | Nilac3 | Brain/muscle | Active time | <0.05 | 2 | 165.664893-165.665019 |  |  |
| 133 | Umemori 2009 | Transcript | Dloc1 | Dloc1 | Brain/muscle | Active time | <0.05 | 2 | - |  |  |
| 134 | Yang 2009 | Transcript | D13Mit254 | D13Mit254 | Brain/muscle | Daily wheel running | <0.0001 | 13 | 76.1326-76.132738 |  |  |
| 135 | Furuse 2002 | Transcript | Loco1 | Loco1 | Brain/muscle | Spontaneous activity | <0.05 | 3 | 150.461893-150.462042 |  |  |
| 136 | Furuse 2002 | Transcript | Loco2 | Loco2 | Brain/muscle | Spontaneous activity | <0.05 | 17 | 43.884972-43.885114 |  |  |
| 137 | Caetano-Anolles 2016 | Transcript | Adora2a | Adora2a | Brain | High Physical Activity | <0.005 | 10 | 75.152711-75.170618 |  |  |
| 138 | Caetano-Anolles 2016 | Transcript | Arhgap8 | Arhgap8 | Brain | High Physical Activity | <0.005 | 15 | 84.604253-84.656408 |  |  |
| 139 | Caetano-Anolles 2016 | Transcript | Cpne4 | Cpne4 | Brain | High Physical Activity | <0.005 | 9 | 104.4439-104.911747 |  |  |
| 140 | Caetano-Anolles 2016 | Transcript | Cpne5 | Cpne5 | Brain | High Physical Activity | <0.005 | 17 | 29.375495-29.456764 |  |  |
| 141 | Caetano-Anolles 2016 | Transcript | Ctxn1 | Ctxn1 | Brain | High Physical Activity | <0.005 | 8 | 4.30766-4.309274 |  |  |
| 142 | Caetano-Anolles 2016 | Transcript | Ddn | Ddn | Brain | High Physical Activity | <0.005 | 15 | 98.701663-98.705806 |  |  |
| 143 | Caetano-Anolles 2016 | Transcript | Drd2 | Drd2 | Brain | High Physical Activity | <0.005 | 9 | 49.251927-49.319477 |  |  |
| 144 | Caetano-Anolles 2016 | Transcript | Egr3 | Egr3 | Brain | High Physical Activity | <0.005 | 14 | 70.314766-70.320062 |  |  |
| 145 | Caetano-Anolles 2016 | Transcript | Gda | Gda | Brain | High Physical Activity | <0.005 | 19 | 21.368671-21.450025 |  |  |
| 146 | Caetano-Anolles 2016 | Transcript | Gpr88 | Gpr88 | Brain | High Physical Activity | <0.005 | 3 | 116.043303-116.047152 |  |  |
| 147 | Caetano-Anolles 2016 | Transcript | Icam5 | Icam5 | Brain | High Physical Activity | <0.005 | 9 | 20.943369-20.950332 |  |  |
| 148 | Caetano-Anolles 2016 | Transcript | Kcnj4 | Kcnj4 | Brain | High Physical Activity | <0.005 | 15 | 79.367915-79.389442 |  |  |
| 149 | Caetano-Anolles 2016 | Transcript | Lamp5 | Lamp5 | Brain | High Physical Activity | <0.005 | 2 | 135.894159-135.911837 |  |  |
| 150 | Caetano-Anolles 2016 | Transcript | Lrrc10b | Lrrc10b | Brain | High Physical Activity | <0.005 | 19 | 10.432735-10.434811 |  |  |
| 151 | Caetano-Anolles 2016 | Transcript | Nrgn | Nrgn | Brain | High Physical Activity | <0.005 | 9 | 37.455789-37.464041 |  |  |
| 152 | Caetano-Anolles 2016 | Transcript | Nts | Nts | Brain | High Physical Activity | <0.005 | 10 | 102.317617-102.326294 |  |  |
| 153 | Caetano-Anolles 2016 | Transcript | Rxrg | Rxrg | Brain | High Physical Activity | <0.005 | 1 | 167.425953-167.467192 |  |  |
| 154 | Caetano-Anolles 2016 | Transcript | Scd4 | Scd4 | Brain | High Physical Activity | <0.005 | 19 | 44.321765-44.335182 |  |  |
| 155 | Caetano-Anolles 2016 | Transcript | Sst | Sst | Brain | High Physical Activity | <0.005 | 16 | 23.708323-23.709708 |  |  |
| 156 | Caetano-Anolles 2016 | Transcript | Tbr1 | Tbr1 | Brain | High Physical Activity | <0.005 | 2 | 61.633274-61.644458 |  |  |
| 157 | Caetano-Anolles 2016 | Transcript | Foxg1 | Foxg1 | Brain | High Physical Activity | <0.005 | 12 | 49.429666-49.43365 |  |  |
| 158 | Caetano-Anolles 2016 | Transcript | Kcnv1 | Kcnv1 | Brain | High Physical Activity | <0.005 | 15 | 44.96968-44.978316 |  |  |
| 159 | Caetano-Anolles 2016 | Transcript | Sp9 | Sp9 | Brain | High Physical Activity | <0.005 | 2 | 73.094805-73.106115 |  |  |
| 160 | Caetano-Anolles 2016 | Transcript | Dlx1 | Dlx1 | Brain | High Physical Activity | <0.005 | 2 | 71.358457-71.364325 |  |  |
| 161 | Caetano-Anolles 2016 | Transcript | Gpr6 | Gpr6 | Brain | High Physical Activity | <0.005 | 10 | 40.945973-40.948281 |  |  |
| 162 | Caetano-Anolles 2016 | Transcript | Drd1 | Drd1 | Brain | High Physical Activity | <0.005 | 13 | 54.205202-54.209677 |  |  |
| 163 | Caetano-Anolles 2016 | Transcript | Zfp831 | Zfp831 | Brain | High Physical Activity | <0.005 | 2 | 174.485327-174.552625 |  |  |
| 164 | Caetano-Anolles 2016 | Transcript | Kcnj4 | Kcnj4 | Brain | High Physical Activity | <0.005 | 15 | 79.367915-79.389442 |  |  |
| 165 | Caetano-Anolles 2016 | Transcript | Ankrd63 | Ankrd63 | Brain | High Physical Activity | <0.005 | 2 | 118.529584-118.534444 |  |  |
| 166 | Caetano-Anolles 2016 | Transcript | Kcnf1 | Kcnf1 | Brain | High Physical Activity | <0.005 | 12 | 17.222101-17.226889 |  |  |
| 167 | Caetano-Anolles 2016 | Transcript | Chrm1 | Chrm1 | Brain | High Physical Activity | <0.005 | 19 | 8.641369-8.66097 |  |  |
| 168 | Caetano-Anolles 2016 | Transcript | Kcng1 | Kcng1 | Brain | High Physical Activity | <0.005 | 2 | 168.102037-168.123453 |  |  |
| 169 | Caetano-Anolles 2016 | Transcript | Dlx2 | Dlx2 | Brain | High Physical Activity | <0.005 | 2 | 71.373752-71.377098 |  |  |
| 170 | Caetano-Anolles 2016 | Transcript | Rprml | Rprml | Brain | High Physical Activity | <0.005 | 11 | 103.540396-103.541405 |  |  |
| 171 | Caetano-Anolles 2016 | Transcript | Crhbp | Crhbp | Brain | High Physical Activity | <0.005 | 13 | 95.567884-95.581339 |  |  |
| 172 | Caetano-Anolles 2016 | Transcript | Rtn4rl2 | Rtn4rl2 | Brain | High Physical Activity | <0.005 | 2 | 84.702268-84.717054 |  |  |
| 173 | Caetano-Anolles 2016 | Transcript | Vdr | Vdr | Brain | High Physical Activity | <0.005 | 15 | 97.752308-97.806177 |  |  |
| 174 | Caetano-Anolles 2016 | Transcript | Actn2 | Actn2 | Brain | High Physical Activity | <0.005 | 13 | 12.284312-12.355613 |  |  |
| 175 | Caetano-Anolles 2016 | Transcript | Ptprv | Ptprv | Brain | High Physical Activity | <0.005 | 1 | 135.036236-135.060313 |  |  |
| 176 | Caetano-Anolles 2016 | Transcript | Kcnh4 | Kcnh4 | Brain | High Physical Activity | <0.005 | 11 | 100.631202-100.650768 |  |  |
| 177 | Caetano-Anolles 2016 | Transcript | Neurl1b | Neurl1b | Brain | High Physical Activity | <0.005 | 17 | 26.633833-26.665295 |  |  |
| 178 | Caetano-Anolles 2016 | Transcript | Sst | Sst | Brain | High Physical Activity | <0.005 | 16 | 23.708323-23.709708 |  |  |
| 179 | Caetano-Anolles 2016 | Transcript | D430019H16Rik | D430019H16Rik | Brain | High Physical Activity | <0.005 | 12 | 105.420115-105.459354 |  |  |
| 180 | Caetano-Anolles 2016 | Transcript | Prss12 | Prss12 | Brain | High Physical Activity | <0.005 | 3 | 123.240562-123.300246 |  |  |
| 181 | Caetano-Anolles 2016 | Transcript | Rab40b | Rab40b | Brain | High Physical Activity | <0.005 | 11 | 121.246951-121.279077 |  |  |
| 182 | Saul 2017 | Transcript | 5031434O11Rik | 5031434O11Rik | Brain | high | <0.05 | 3 | 51.467456-51.474538 |  |  |
| 183 | Saul 2017 | Transcript | Gak | Gak | Brain | high physical activity | <0.05 | 5 | 108.717277-108.777621 |  |  |
| 184 | Saul 2017 | Transcript | Mfsd7a | Mfsd7a | Brain | low physical activity | <0.05 | 5 | 108.58892-108.596966 |  |  |
| 185 | Saul 2017 | Transcript | Gpr3 | Gpr3 | Brain | high physical activity | <0.05 | 4 | 132.936651-132.939847 |  |  |
| 186 | Saul 2017 | Transcript | Slc38a2 | Slc38a2 | Brain | high physical activity | <0.05 | 15 | 96.585273-96.597611 |  |  |
| 187 | Saul 2017 | Transcript | Htr1b | Htr1b | Brain | high physical activity | <0.05 | 9 | 81.510344-81.515881 |  |  |
| 188 | Saul 2017 | Transcript | Tmed5 | Tmed5 | Brain | high | <0.05 | 5 | 108.269422-108.280486 |  |  |
| 189 | Saul 2017 | Transcript | Baz1a | Baz1a | Brain | high physical activity | <0.05 | 12 | 54.939774-55.061133 |  |  |
| 190 | Saul 2017 | Transcript | H1f0 | H1f0 | Brain | high physical activity | <0.05 | 15 | 78.91265-78.914704 |  |  |
| 191 | Pistilli 2011 | Transcript | IL15RA | IL15RA | Muscle | high physical activity | <0.05 | 2 | 11.709992-11.738796 |  |  |
| 192 | Hartmann 2008 | Transcript | sox15 | sox15 | Muscle | high physical activity | <0.05 | 11 | 69.54614-69.547553 |  |  |
| 193 | Hartmann 2008 | Transcript | chrnb1 | chrnb1 | Muscle | high physical activity | <0.05 | 11 | 69.674862-69.686769 |  |  |
| 194 | Hartmann 2008 | Transcript | rpl26 | rpl26 | Muscle | high physical activity | <0.05 | 11 | 68.792392-68.79536 |  |  |
| 195 | Hartmann 2008 | Transcript | Acadvl | Acadvl | Muscle | high physical activity | <0.05 | 11 | 69.901009-69.906237 |  |  |
| 196 | Hartmann 2008 | Transcript | Myh10 | Myh10 | Muscle | high physical activity | <0.05 | 11 | 68.582385-68.707458 |  |  |
| 197 | Hartmann 2008 | Transcript | Slc2a4 | Glut4 | Muscle | high physical activity | <0.05 | 11 | 69.833365-69.839014 |  |  |
| 198 | Hartmann 2008 | Transcript | Trp53 | Trp53 | Muscle | high physical activity | <0.05 | 11 | 69.471185-69.482699 |  |  |
| 199 | Kelly 2010 | Genome variant | rs31675929 |  | Brain/muscle | Distance | <0.05 | 1 | 3.466797 |  |  |
| 200 | Kelly 2010 | Genome variant | rs31317013 |  | Brain/muscle | Distance | <0.05 | 1 | 110.837757 |  |  |
| 201 | Kelly 2010 | Genome variant | rs30737566 |  | Brain/muscle | Distance | <0.05 | 1 | 120.69282 |  |  |
| 202 | Kelly 2010 | Genome variant | rs30816406 |  | Brain/muscle | Time | <0.05 | 1 | 134.462768 |  |  |
| 203 | Kelly 2010 | Genome variant | rs30241278 |  | Brain/muscle | Time | <0.05 | 1 | 66.283718 |  |  |
| 204 | Kelly 2010 | Genome variant | rs30737566 |  | Brain/muscle | Time | <0.05 | 1 | 120.69282 |  |  |
| 205 | Kelly 2010 | Genome variant | rs3677375 |  | Brain/muscle | Time | <0.05 | 1 | 132.481712 |  |  |
| 206 | Nehrenberg 2010 | Genome variant | rs13476818 |  | Brain/muscle | Running wheel duration | <0.05 | 2 | 144.75696 |  |  |
| 207 | Kelly 2010 | Genome variant | rs27380445 |  | Brain/muscle | Average Running Speed | <0.05 | 2 | 99.239681 |  |  |
| 208 | Kelly 2010 | Genome variant | rs28032050 |  | Brain/muscle | Maximum Running Speed | <0.05 | 2 | 82.974913 |  |  |
| 209 | Kelly 2010 | Genome variant | rs27368335 |  | Brain/muscle | Maximum Running Speed | <0.05 | 2 | 92.032833 |  |  |
| 210 | Kelly 2010 | Genome variant | rs27368335 |  | Brain/muscle | Maximum Running Speed | <0.05 | 2 | 92.032833 |  |  |
| 211 | Nehrenberg 2010 | Genome variant | rs13478156 |  | Brain/muscle | Running wheel maximum speed and avg. speed | <0.05 | 5 | 28.007545 |  |  |
| 212 | Kelly 2010 | Genome variant | rs48305016 |  | Brain/muscle | Distance | <0.05 | 5 | 52.616733 |  |  |
| 213 | Kelly 2010 | Genome variant | rs48305016 |  | Brain/muscle | Time | <0.05 | 5 | 52.616733 |  |  |
| 214 | Lightfoot 2010 | Genome variant | rs47227633 |  | Brain/muscle | Distance | <0.001 | 5 | 118 |  |  |
| 215 | Nehrenberg 2010 | Genome variant |  |  | Brain/muscle | Running wheel maximum speed | <0.05 | 6 | NA |  |  |
| 216 | Kelly 2010 | Genome variant | rs30981553 |  | Brain/muscle | Distance | <0.05 | 6 | 46.884541 |  |  |
| 217 | Kelly 2010 | Genome variant | rs30462460 |  | Brain/muscle | Distance | <0.05 | 6 | 36.37036 |  |  |
| 218 | Kelly 2010 | Genome variant | rs30462460 |  | Brain/muscle | Distance | <0.05 | 6 | 36.37036 |  |  |
| 219 | Kelly 2010 | Genome variant | rs33841297 |  | Brain/muscle | Time | <0.05 | 6 | 28.866986 |  |  |
| 220 | Kelly 2010 | Genome variant | rs30462460 |  | Brain/muscle | Time | <0.05 | 6 | 36.37036 |  |  |
| 221 | Lightfoot 2010 | Genome variant | rs30945756 |  | Brain/muscle | Distance | <0.001 | 6 | 145.505475 |  |  |
| 222 | Lightfoot 2010 | Genome variant | rs31691517 |  | Brain/muscle | Speed | <0.001 | 6 | 119.815155 |  |  |
| 223 | Nehrenberg 2010 | Genome variant |  |  | Brain/muscle | Running wheel maximum speed and avg. speed | <0.05 | 7 | NA |  |  |
| 224 | Kelly 2010 | Genome variant | rs31104131 |  | Brain/muscle | Distance | <0.05 | 7 | 109.127278 |  |  |
| 225 | Kelly 2010 | Genome variant | rs3675993 |  | Brain/muscle | Distance | <0.05 | 7 | 115.14664 |  |  |
| 226 | Kelly 2010 | Genome variant | rs33245012 |  | Brain/muscle | Time | <0.05 | 7 | 82.760104 |  |  |
| 227 | Kelly 2010 | Genome variant | rs31104131 |  | Brain/muscle | Time | <0.05 | 7 | 109.127278 |  |  |
| 228 | Kelly 2010 | Genome variant | rs32458319 |  | Brain/muscle | Time | <0.05 | 7 | 90.075056 |  |  |
| 229 | Kelly 2010 | Genome variant | rs31923186 |  | Brain/muscle | Time | <0.05 | 7 | 122.591825 |  |  |
| 230 | Kelly 2010 | Genome variant | rs31104131 |  | Brain/muscle | Time | <0.05 | 7 | 109.127278 |  |  |
| 231 | Kelly 2010 | Genome variant | rs31104131 |  | Brain/muscle | Time | <0.05 | 7 | 109.127278 |  |  |
| 232 | Lightfoot 2010 | Genome variant | rs49863494 |  | Brain/muscle | Distance | <0.001 | 8 | 58.894155 |  |  |
| 233 | Lightfoot 2010 | Genome variant | rs32100214 |  | Brain/muscle | Distance | <0.001 | 8 | 94.559416 |  |  |
| 234 | Lightfoot 2008 | Genome variant | rs13480073 |  | Brain/muscle | Running wheel speed of exercise | <0.01 | 9 | 13.430544 |  |  |
| 235 | Kelly 2010 | Genome variant | rs26880493 |  | Brain/muscle | Maximum Running Speed | <0.05 | 11 | 10.038702 |  |  |
| 236 | Kelly 2010 | Genome variant | rs26945355 |  | Brain/muscle | Maximum Running Speed | <0.05 | 11 | 53.393768 |  |  |
| 237 | Kelly 2010 | Genome variant | rs26945355 |  | Brain/muscle | Maximum Running Speed | <0.05 | 11 | 53.393768 |  |  |
| 238 | Kelly 2010 | Genome variant | rs26880493 |  | Brain/muscle | Maximum Running Speed | <0.05 | 11 | 10.038702 |  |  |
| 239 | Kelly 2010 | Genome variant | rs26945355 |  | Brain/muscle | Maximum Running Speed | <0.05 | 11 | 53.393768 |  |  |
| 240 | Lightfoot 2010 | Genome variant | rs26981291 |  | Brain/muscle | Distance | <0.001 | 11 | 85.230648 |  |  |
| 241 | Lightfoot 2010 | Genome variant | rs28225821 |  | Brain/muscle | Speed | <0.001 | 11 | 84.026786 |  |  |
| 242 | Kelly 2010 | Genome variant | rs32432654 |  | Brain/muscle | Average Running Speed | <0.05 | 12 | 75.702379 |  |  |
| 243 | Lightfoot 2010 | Genome variant | rs46860253 |  | Brain/muscle | Distance | <0.001 | 12 | 88.343051 |  |  |
| 244 | Lightfoot 2008 | Genome variant | rs6329684 |  | Brain/muscle | Running wheel distance/day | <0.01 | 13 | 10.065818 |  |  |
| 245 | Lightfoot 2008 | Genome variant | rs6329684 |  | Brain/muscle | Running wheel duration of exercise | <0.01 | 13 | 10.065818 |  |  |
| 246 | Lightfoot 2008 | Genome variant | rs6329684 |  | Brain/muscle | Running wheel speed of exercise | <0.01 | 13 | 10.065818 |  |  |
| 247 | Kelly 2010 | Genome variant | rs29736244 |  | Brain/muscle | Time | <0.05 | 13 | 10.545336 |  |  |
| 248 | Yang 2012 | Genome variant | rs3720620 |  | Brain/muscle | Running wheel activity + wheel runnning ambulations | <0.05 | 13 | 38.753652 |  |  |
| 249 | Lightfoot 2010 | Genome variant | rs46617906 |  | Brain/muscle | Distance | <0.001 | 13 | 95.707495 |  |  |
| 250 | Kelly 2010 | Genome variant | ? |  | Brain/muscle | Average Running Speed | <0.05 | 14 | 81.164271 |  |  |
| 251 | Kelly 2010 | Genome variant | rs50154491 |  | Brain/muscle | Average Running Speed | <0.05 | 17 | 33.542759 |  |  |
| 252 | Lightfoot 2010 | Genome variant | rs3692956 |  | Brain/muscle | Distance | <0.001 | 18 | 11.417832 |  |  |
| 253 | Kelly 2010 | Genome variant | rs30405098 |  | Brain/muscle | Time | <0.05 | 19 | 46.380357 |  |  |
| 254 | Lightfoot 2010 | Genome variant | rs36666297 |  | Brain/muscle | Distance | <0.001 | 19 | 16.036727 |  |  |
| 255 | Lightfoot 2010 | Genome variant | rs33880738 |  | Brain/muscle | Duration | <0.001 | X | 111.216615 |  |  |
| 256 | Kostrzewa 2014 | Genome variant | rs27289254 |  | Brain/muscle | wheel running activity | <0.05 | 2 | 171.933137 |  |  |
| 257 | Kas 2009 | Genome variant | rs32957531 |  | Brain/muscle | Motor activity | <0.05 | 1 | 80.207361 |  |  |
| 258 | Kas 2009 | Genome variant | rs32957529 |  | Brain/muscle | Motor activity | <0.05 | 1 | 80.207911 |  |  |
| 259 | Leamy 2010 | Genome variant | rs13478553 |  | Brain/muscle | Distance | <0.01 | 5 | 138.345042 |  |  |
| 260 | Leamy 2010 | Genome variant | rs13479600 |  | Brain/muscle | Distance | <0.01 | 8 | NA |  |  |
| 261 | Leamy 2010 | Genome variant | rs13480073 |  | Brain/muscle | Distance | <0.01 | 9 | 13.430544 |  |  |
| 262 | Leamy 2010 | Genome variant | rs6329684 |  | Brain/muscle | Distance | <0.01 | 13 | 10.065818 |  |  |
| 263 | Leamy 2010 | Genome variant | rs13476352 |  | Brain/muscle | Distance | <0.05 | 2 | 13.436218 |  |  |
| 264 | Leamy 2010 | Genome variant | gnf06.086.089 |  | Brain/muscle | Distance | <0.01 | NA | NA |  |  |
| 265 | Leamy 2010 | Genome variant | rs3714636 |  | Brain/muscle | Distance | <0.01 | 7 | 144.946744 |  |  |
| 266 | Leamy 2010 | Genome variant | rs3703161 |  | Brain/muscle | Distance | <0.05 | 8 | 128.357744 |  |  |
| 267 | Leamy 2010 | Genome variant | rs13480273 |  | Brain/muscle | Distance | <0.01 | 9 | 73.272754 |  |  |
| 268 | Leamy 2010 | Genome variant | rs3712998 |  | Brain/muscle | Distance | <0.01 | 10 | 10.193642 |  |  |
| 269 | Leamy 2010 | Genome variant | rs13480273 |  | Brain/muscle | Distance | <0.01 | 9 | 73.272754 |  |  |
| 270 | Leamy 2010 | Genome variant | rs6361467 |  | Brain/muscle | Distance | <0.01 | 12 | 110.451747 |  |  |
| 271 | Leamy 2010 | Genome variant | rs13481855 |  | Brain/muscle | Distance | <0.01 | 13 | 65.296578 |  |  |
| 272 | Leamy 2010 | Genome variant | rs6154379 |  | Brain/muscle | Duration | <0.01 | 1 | 185.177962 |  |  |
| 273 | Leamy 2010 | Genome variant | rs3715009 |  | Brain/muscle | Duration | <0.01 | 4 | 59.388602 |  |  |
| 274 | Leamy 2010 | Genome variant | CEL-5_11773662 |  | Brain/muscle | Duration | <0.01 | NA | NA |  |  |
| 275 | Leamy 2010 | Genome variant | gnf06112.868 |  | Brain/muscle | Duration | <0.05 | NA | NA |  |  |
| 276 | Leamy 2010 | Genome variant | rs13479600 |  | Brain/muscle | Duration | <0.05 | 8 | 8.884954 |  |  |
| 277 | Leamy 2010 | Genome variant | rs3659852 |  | Brain/muscle | Duration | <0.01 | 8 | NA |  |  |
| 278 | Leamy 2010 | Genome variant | rs13480073 |  | Brain/muscle | Duration | <0.01 | 9 | 13.430544 |  |  |
| 279 | Leamy 2010 | Genome variant | rs13480409 |  | Brain/muscle | Duration | <0.01 | 9 | NA |  |  |
| 280 | Leamy 2010 | Genome variant | rs6329684 |  | Brain/muscle | Duration | <0.01 | 13 | NA |  |  |
| 281 | Leamy 2010 | Genome variant | rs13481855 |  | Brain/muscle | Duration | <0.01 | 13 | NA |  |  |
| 282 | Leamy 2010 | Genome variant | rs3708665 |  | Brain/muscle | Duration | <0.01 | 14 | 100.861042 |  |  |
| 283 | Leamy 2010 | Genome variant | rs6329684 |  | Brain/muscle | Duration | <0.01 | 13 | 10.065818 |  |  |
| 284 | Leamy 2010 | Genome variant | rs8259436 |  | Brain/muscle | Duration | <0.05 | 15 | 74.849949 |  |  |
| 285 | Leamy 2010 | Genome variant | rs13476352 |  | Brain/muscle | Speed | <0.01 | 2 | 13.436218 |  |  |
| 286 | Leamy 2010 | Genome variant | rs3664044 |  | Brain/muscle | Speed | <0.01 | 2 | 173.607823 |  |  |
| 287 | Leamy 2010 | Genome variant | rs13478157 |  | Brain/muscle | Speed | <0.01 | 5 | NA |  |  |
| 288 | Leamy 2010 | Genome variant | mCV22996021 |  | Brain/muscle | Speed | <0.01 | NA | NA |  |  |
| 289 | Leamy 2010 | Genome variant | rs3023193 |  | Brain/muscle | Speed | <0.01 | 8 | 84.721303 |  |  |
| 290 | Leamy 2010 | Genome variant | rs3703161 |  | Brain/muscle | Speed | <0.01 | 8 | NA |  |  |
| 291 | Leamy 2010 | Genome variant | rs13480073 |  | Brain/muscle | Speed | <0.01 | 9 | NA |  |  |
| 292 | Leamy 2010 | Genome variant | rs3704618 |  | Brain/muscle | Speed | <0.01 | 10 | 29.037602 |  |  |
| 293 | Leamy 2010 | Genome variant | mCV24434350 |  | Brain/muscle | Speed | <0.01 | NA | NA |  |  |
| 294 | Leamy 2010 | Genome variant | rs6329684 |  | Brain/muscle | Speed | <0.01 | 13 | 10.065818 |  |  |
| 295 | Leamy 2010 | Genome variant | rs13482429 |  | Brain/muscle | Speed | <0.01 | 15 | 10.575512 |  |  |
| 296 | Leamy 2010 | Genome variant | rs3686133 |  | Brain/muscle | Speed | <0.01 | 15 | 102.360423 |  |  |
| 297 | Leamy 2010 | Genome variant | rs3671328 |  | Brain/muscle | Speed | <0.01 | 19 | 12.624583 |  |  |
| 298 | Leamy 2010 | Genome variant | rs3713675 |  | Brain/muscle | Slope of Activity Traits | <0.05 | 9 | 33.309504 |  |  |
| 299 | Leamy 2010 | Genome variant | rs13480409 |  | Brain/muscle | Slope of Activity Traits | <0.05 | 9 | 109.089545 |  |  |
| 300 | Leamy 2010 | Genome variant | rs6259521 |  | Brain/muscle | Slope of Activity Traits | <0.01 | 9 | 23.946802 |  |  |
| 301 | Leamy 2010 | Genome variant | rs4136370 |  | Brain/muscle | Slope of Activity Traits | <0.01 | 4 | 91.354676 |  |  |
| 302 | Leamy 2010 | Genome variant | rs3713675 |  | Brain/muscle | Slope of Activity Traits | <0.01 | 9 | 33.309504 |  |  |
| 303 | Leamy 2010 | Genome variant | rs13480409 |  | Brain/muscle | Slope of Activity Traits | <0.01 | 9 | 109.089545 |  |  |
| 304 | Leamy 2010 | Genome variant | rs13480409 |  | Brain/muscle | Slope of Activity Traits | <0.01 | 9 | 109.089545 |  |  |
| 305 | Dawes 2015 | microRNA | mmu‐miR‐375 |  | brain | high or low wheel activity | 0.03 | 1 | 74.947292 |  |  |
| 306 | Dawes 2015 | microRNA | mmu‐miR‐5117 |  | brain | high or low wheel activity | <0.001 | 1 | 162.967492 |  |  |
| 307 | Dawes 2015 | microRNA | mmu‐miR‐205 |  | muscle | high or low wheel activity | <0.001 | 1 | 195.333684 |  |  |
| 308 | Dawes 2015 | microRNA | mmu‐miR‐5117 |  | muscle | high or low wheel activity | <0.001 | 1 | 162.967492 |  |  |
| 309 | Dawes 2015 | microRNA | mmu‐miR‐1927 |  | muscle | high or low wheel activity | <0.001 | 1 | 162.226082 |  |  |
| 310 | Dawes 2015 | microRNA | mmu‐miR‐ 5103 |  | muscle | high or low wheel activity | <0.001 | 1 | 34.490035 |  |  |
| 311 | Dawes 2015 | microRNA | mmu‐miR‐ 5117 |  | muscle | high or low wheel activity | <0.001 | 1 | 162.967492 |  |  |
| 312 | Dawes 2015 | microRNA | mmu‐miR‐466d‐3p |  | brain | high or low wheel activity | <0.001 | 2 | 1.043365 |  |  |
| 313 | Dawes 2015 | microRNA | mmu‐miR‐467ac |  | brain | high or low wheel activity | <0.01 | 2 | 10.398019 |  |  |
| 314 | Dawes 2015 | microRNA | mmu‐miR‐467b |  | brain | high or low wheel activity | <0.01 | 2 | 10.402887 |  |  |
| 315 | Dawes 2015 | microRNA | mmu‐miR‐467c |  | brain | high or low wheel activity | <0.001 | 2 | 10.395572 |  |  |
| 316 | Dawes 2015 | microRNA | mmu‐miR‐669f‐3p |  | brain | high or low wheel activity | <0.01 | 2 | 10.388917 |  |  |
| 317 | Dawes 2015 | microRNA | mmu‐miR‐6691 |  | brain | high or low wheel activity | <0.001 | 2 | 10.390015 |  |  |
| 318 | Dawes 2015 | microRNA | mmu‐miR‐669n |  | brain | high or low wheel activity | <0.01 | 2 | 10.43095 |  |  |
| 319 | Dawes 2015 | microRNA | mmu‐miR‐3091‐5p |  | muscle | high or low wheel activity | <0.001 | 2 | 179.992262 |  |  |
| 320 | Dawes 2015 | microRNA | mmu‐miR‐466b‐3p |  | muscle | high or low wheel activity | <0.001 | 2 | 10.395901 |  |  |
| 321 | Dawes 2015 | microRNA | mmu‐miR‐467ac |  | muscle | high or low wheel activity | <0.001 | 2 | 10.398019 |  |  |
| 322 | Dawes 2015 | microRNA | mmu‐miR‐467b |  | muscle | high or low wheel activity | <0.001 | 2 | 10.402887 |  |  |
| 323 | Dawes 2015 | microRNA | mmu‐miR‐467e |  | muscle | high or low wheel activity | <0.001 | 2 | 10.427362 |  |  |
| 324 | Dawes 2015 | microRNA | mmu‐miR‐3095‐3p |  | brain | high or low wheel activity | <0.001 | 4 | 58.453959 |  |  |
| 325 | Dawes 2015 | microRNA | mmu‐miR‐1957 |  | muscle | high or low wheel activity | <0.001 | 4 | 118.802399 |  |  |
| 326 | Dawes 2015 | microRNA | mmu‐miR‐1960 |  | muscle | high or low wheel activity | <0.001 | 5 | 30.497297 |  |  |
| 327 | Dawes 2015 | microRNA | mmu‐miR‐148a |  | muscle | high or low wheel activity | 0.02 | 6 | 51.219892 |  |  |
| 328 | Dawes 2015 | microRNA | mmu‐miR‐326 |  | muscle | high or low wheel activity | <0.001 | 7 | 106.700843 |  |  |
| 329 | Dawes 2015 | microRNA | mmu‐miR‐1966 |  | muscle | high or low wheel activity | <0.001 | 8 | 108.139391 |  |  |
| 330 | Dawes 2015 | microRNA | mmu‐miR‐1967 |  | muscle | high or low wheel activity | <0.001 | 8 | 126.546597 |  |  |
| 331 | Dawes 2015 | microRNA | mmu‐miR‐711 |  | muscle | high or low wheel activity | <0.001 | 9 | 108.872022 |  |  |
| 332 | Dawes 2015 | microRNA | mmu‐miR‐193c |  | muscle | high or low wheel activity | <0.001 | 11 | 79.525486 |  |  |
| 333 | Dawes 2015 | microRNA | mmu‐miR‐342‐3p |  | brain | high or low wheel activity | <0.001 | 12 | 109.896897 |  |  |
| 334 | Dawes 2015 | microRNA | mmu‐miR‐1843‐5p |  | muscle | high or low wheel activity | <0.001 | 12 | 81.492623 |  |  |
| 335 | Dawes 2015 | microRNA | mmu‐miR‐342‐5p |  | brain | high or low wheel activity | <0.001 | l2 | 109.896852 |  |  |
| 336 | Dawes 2015 | microRNA | mmu‐miR‐376c |  | brain | high or low wheel activity | <0.01 | l2 | 110.960981 |  |  |
| 337 | Dawes 2015 | microRNA | mmu‐miR‐342‐3p |  | muscle | high or low wheel activity | 0.02 | l2 | 109.896897 |  |  |
| 338 | Dawes 2015 | microRNA | mmu‐miR‐299 |  | muscle | high or low wheel activity | <0.001 | l2 | 110.948892 |  |  |
| 339 | Dawes 2015 | microRNA | mmu‐miR‐342‐3p |  | muscle | high or low wheel activity | <0.001 | l2 | 109.896897 |  |  |
| 340 | Dawes 2015 | microRNA | mmu‐miR‐376a |  | muscle | high or low wheel activity | <0.001 | l2 | 110.962039 |  |  |
| 341 | Dawes 2015 | microRNA | mmu‐miR‐431 |  | muscle | high or low wheel activity | <0.001 | l2 | 110.828676 |  |  |
| 342 | Dawes 2015 | microRNA | mmu‐miR‐582‐5p |  | muscle | high or low wheel activity | <0.001 | 13 | 110.114949 |  |  |
| 343 | Dawes 2015 | microRNA | mmu‐miR‐5118 |  | muscle | high or low wheel activity | <0.001 | l6 | 55.494889 |  |  |
| 344 | Dawes 2015 | microRNA | mmu‐miR‐99ac |  | muscle | high or low wheel activity | <0.001 | l6 | 77.599226 |  |  |
| 345 | Dawes 2015 | microRNA | mmu‐miR‐3 62‐5p |  | muscle | high or low wheel activity | <0.001 | X | 6.819135 |  |  |
| 346 | Hartmann 2008 | microRNA | mmu-miR-324-3P |  | muscle | mini-muscle | <0.01 | 11 | 70.012043 |  |  |
| 347 | Hartmann 2008 | microRNA | mmu-miR-497 |  | muscle | mini-muscle | <0.01 | 11 | 70.234717 |  |  |
| 348 | Hartmann 2008 | microRNA | mmu-miR-195 |  | muscle | mini-muscle | <0.01 | 11 | 70.235042 |  |  |
| 349 | Bruneau 2018 | Genome variant | rs2228059 | IL15RA | Brain/muscle | Light Activity (hr/week) | 0.009 | 10 | 5.960355-5.960455 | 2 | 11.723562-11.7235820 |
| 350 | Cai 2006 | Genome variant | Accel-Cai | MCR4 | Brain/muscle | Accelerometry | 0.001 | 18 | 60.371062-60.372775 |  |  |
| 351 | Comuzzie 2012 | Genome variant | rs16933006 | Closest RPL7P3 | Brain/muscle | Light Activity (min/d) | 7.49E-08 | 9 | 15.335866-15.335966 |  |  |
| 352 | Comuzzie 2012 | Genome variant | rs6025590 | CTCFL | Brain/muscle | Sedentary and Light Activity (min/d) | 3.61E-08 | 20 | 57.495399-57.495499 | 2 | 173.089694-173.089720 |
| 353 | De Moor 2007 | Genome variant | Liabl-DeMoor | D19S247 | Brain/muscle | liability for exercise participation | <0.05 | 19 | 19.2-19.3 |  |  |
| 354 | De Moor 2009 | Genome variant | rs12612420 | Closest DNAPTP6 | Brain/muscle | Exercise Participation | 7.65E-05 | 2 | 200.293349-200.293449 |  |  |
| 355 | De Moor 2009 | Genome variant | rs17592517 | Closest DNAPTP6 | Brain/muscle | Exercise Participation | <1.0E-5 | 2 | 200.292162-200.292262 |  |  |
| 356 | De Moor 2009 | Genome variant | rs10887741 | Closest PAPSS2 | Brain/muscle | Exercise Participation | 6.26E-06 | 10 | 87.683503-87.683603 |  |  |
| 357 | De Moor 2009 | Genome variant | rs1980647 | PAPSS2 | Brain/muscle | Exercise Participation | <1.0E-5 | 10 | 87.66248-87.66258 | 19 | 32.598663-32.598680 |
| 358 | De Moor 2009 | Genome variant | rs4934355 | PAPSS2 | Brain/muscle | Exercise Participation | <1.0E-5 | 10 | 87.665604-87.665704 | 19 | 32.600638-32.600651 |
| 359 | De Moor 2009 | Genome variant | rs1358864 | PAPSS2 | Brain/muscle | Exercise Participation | <1.0E-5 | 10 | 87.666823-87.666923 | 19 | 32.601621-32.601644 |
| 360 | De Moor 2009 | Genome variant | rs7903568 | PAPSS2 | Brain/muscle | Exercise Participation | <1.0E-5 | 10 | 87.674124-87.674224 |  |  |
| 361 | De Moor 2009 | Genome variant | rs2077695 | PAPSS2 | Brain/muscle | Exercise Participation | <1.0E-5 | 10 | 87.675676-87.675776 | 19 | 32.610155-32.610169 |
| 362 | De Moor 2009 | Genome variant | rs10218939 | PAPSS2 | Brain/muscle | Exercise Participation | <1.0E-5 | 10 | 87.677046-87.677146 |  |  |
| 363 | De Moor 2009 | Genome variant | rs11202501 | PAPSS2 | Brain/muscle | Exercise Participation | <1.0E-5 | 10 | 87.677401-87.677501 | 19 | 32.611441-32.611461 |
| 364 | De Moor 2009 | Genome variant | rs12412482 | PAPSS2 | Brain/muscle | Exercise Participation | <1.0E-5 | 10 | 87.683992-87.684092 | 19 | 32.617716-32.617737 |
| 365 | De Moor 2009 | Genome variant | rs7908056 | PAPSS2 | Brain/muscle | Exercise Participation | <1.0E-5 | 10 | 87.684843-87.684943 |  |  |
| 366 | De Moor 2009 | Genome variant | rs8097348 | Closest C18orf2 | Brain/muscle | Exercise Participation | 6.99E-05 | 18 | 1.59497-1.59507 |  |  |
| 367 | De Moor 2009 | Genome variant | rs4502301 | C18orf2 | Brain/muscle | Exercise Participation | <1.0E-5 | 18 | 1.594502-1.594602 |  |  |
| 368 | De Moor 2009 | Genome variant | rs2345036 | C18orf2 | Brain/muscle | Exercise Participation | <1.0E-5 | 18 | 1.596485-1.596585 |  |  |
| 369 | De Moor 2009 | Genome variant | rs2111926 | C18orf2 | Brain/muscle | Exercise Participation | <1.0E-5 | 18 | 1.597098-1.597198 |  |  |
| 370 | De Moor 2009 | Genome variant | rs11080871 | C18orf2 | Brain/muscle | Exercise Participation | <1.0E-5 | 18 | 1.596838-1.596938 |  |  |
| 371 | De Moor 2009 | Genome variant | rs2111925 | C18orf2 | Brain/muscle | Exercise Participation | <1.0E-5 | 18 | 1.597358-1.597458 |  |  |
| 372 | De Moor 2009 | Genome variant | rs2160961 | C18orf2 | Brain/muscle | Exercise Participation | <1.0E-5 | 18 | 1.597385-1.597485 |  |  |
| 373 | De Moor 2009 | Genome variant | rs2052420 | C18orf2 | Brain/muscle | Exercise Participation | <1.0E-5 | 18 | 1.597852-1.597952 |  |  |
| 374 | De Moor 2009 | Genome variant | rs2052419 | C18orf2 | Brain/muscle | Exercise Participation | <1.0E-5 | 18 | 1.597931-1.598031 |  |  |
| 375 | De Moor 2009 | Genome variant | rs12605872 | C18orf2 | Brain/muscle | Exercise Participation | <1.0E-5 | 18 | 1.598026-1.598126 |  |  |
| 376 | De Moor 2009 | Genome variant | rs9646479 | C18orf2 | Brain/muscle | Exercise Participation | <1.0E-5 | 18 | 1.598132-1.598232 |  |  |
| 377 | De Moor 2009 | Genome variant | rs9646456 | C18orf2 | Brain/muscle | Exercise Participation | <1.0E-5 | 18 | 1.598196-1.598296 |  |  |
| 378 | De Moor 2009 | Genome variant | rs8095975 | C18orf2 | Brain/muscle | Exercise Participation | <1.0E-5 | 18 | 1.598836-1.598936 |  |  |
| 379 | De Moor 2009 | Genome variant | rs8083526 | C18orf2 | Brain/muscle | Exercise Participation | <1.0E-5 | 18 | 1.599038-1.599138 |  |  |
| 380 | De Moor 2009 | Genome variant | rs4389203 | C18orf2 | Brain/muscle | Exercise Participation | <1.0E-5 | 18 | 1.599073-1.599173 |  |  |
| 381 | De Moor 2009 | Genome variant | rs8099756 | C18orf2 | Brain/muscle | Exercise Participation | <1.0E-5 | 18 | 1.599099-1.599199 |  |  |
| 382 | De Moor 2009 | Genome variant | rs4797926 | C18orf2 | Brain/muscle | Exercise Participation | <1.0E-5 | 18 | 1.599161-1.599261 |  |  |
| 383 | De Moor 2009 | Genome variant | rs4797928 | C18orf2 | Brain/muscle | Exercise Participation | <1.0E-5 | 18 | 1.599212-1.599312 |  |  |
| 384 | De Moor 2009 | Genome variant | rs4797929 | C18orf2 | Brain/muscle | Exercise Participation | <1.0E-5 | 18 | 1.59933-1.59943 |  |  |
| 385 | De Moor 2009 | Genome variant | rs4797930 | C18orf2 | Brain/muscle | Exercise Participation | <1.0E-5 | 18 | 1.599447-1.599547 |  |  |
| 386 | De Moor 2009 | Genome variant | rs1869532 | C18orf2 | Brain/muscle | Exercise Participation | <1.0E-5 | 18 | 1.60003-1.60013 |  |  |
| 387 | De Moor 2009 | Genome variant | rs1869533 | C18orf2 | Brain/muscle | Exercise Participation | <1.0E-5 | 18 | 1.600165-1.600265 |  |  |
| 388 | De Moor 2009 | Genome variant | rs16941818 | C18orf2 | Brain/muscle | Exercise Participation | <1.0E-5 | 18 | 1.600894-1.600994 |  |  |
| 389 | De Moor 2009 | Genome variant | rs7229607 | C18orf2 | Brain/muscle | Exercise Participation | <1.0E-5 | 18 | 1.601616-1.601716 |  |  |
| 390 | De Moor 2009 | Genome variant | rs1051393 | IFNAR2 | Brain/muscle | Exercise Participation | <1.0E-5 | 21 | 33.2419-33.242 | 16 | 91.383916-91.383936 |
| 391 | Doherty 2018 | Genome variant | rs1858242 | LOC105377146 | Brain/muscle | Sedentary | 3.10E-09 | 3 | 68.477934-68.478034 | 6 | 96.570013-96.570033 |
| 392 | Doherty 2018 | Genome variant | rs26579 | MEF2C-AS2 | Brain/muscle | Sedentary | 2.60E-09 | 5 | 88.689428-88.689528 | 13 | 83.716752-83.716772 |
| 393 | Doherty 2018 | Genome variant | rs25981 | EFNAS | Brain/muscle | Sedentary | 3.00E-09 | 5 | 107.487157-107.487257 | 17 | 62.708832-62.708838 |
| 394 | Doherty 2018 | Genome variant | rs34858520 | CALN1 | Brain/muscle | Sedentary | 4.20E-09 | 7 | 72.258848-72.258948 |  |  |
| 395 | Doherty 2018 | Genome variant | rs2942357 | SKIDA1 | Brain/muscle | Overall Activity | 4.20E-09 | 10 | 16.985894-16.985994 | 2 | 13.384810-13.384817 |
| 396 | Gielen 2014 | Genome variant | rs8192678 | PPARGC1A | Brain/muscle | High Intensity PA Participation | 0.001 | 4 | 23.813989-23.814089 | 5 | 51.473835-51.473855 |
| 397 | Gielen 2014 | Genome variant | rs2267668 | PPARD | Brain/muscle | Habitual PA Participation | 0.005 | 6 | 35.410095-35.410195 |  |  |
| 398 | Gielen 2014 | Genome variant | rs2076168 | PPARD | Brain/muscle | Habitual PA Participation | 0.006 | 6 | 35.422172-35.422272 | 17 | 28.297420-28.297441 |
| 399 | Klimentidis 2017 | Genome variant | rs12142550 | Closest HAXI | Brain/muscle | MVPA | 4.60E-08 | 1 | 154.286023-154.286123 |  |  |
| 400 | Klimentidis 2017 | Genome variant | rs6667222 | HAX1 & UBAP2L | Brain/muscle | Vigorous PA: >3 vs 0 days/week | 9.55E-09 | 1 | 154.281135-154.281235 |  |  |
| 401 | Klimentidis 2017 | Genome variant | rs705692 | CAMTA1 | Brain/muscle | Strenuous Sports or Exercise: >2-3 vs 0 days/week | 1.53E-08 | 1 | 7.420107-7.420207 |  |  |
| 402 | Klimentidis 2017 | Genome variant | rs34517439 | FUBP1 (DNAJB4) | Brain/muscle | Accelerometry - Avg Acceleration | 4.23E-08 | 1 | 77.984783-77.984883 |  |  |
| 403 | Klimentidis 2017 | Genome variant | rs181053839 | WDPCP | Brain/muscle | Strenuous Sports or Exercise: >2-3 vs 0 days/week | 3.02E-08 | 2 | 63.258956-63.259056 | 1 | 21.814126-21.814139 |
| 404 | Klimentidis 2017 | Genome variant | rs4361077 | Closest ACVR1 | Brain/muscle | MVPA, VPA, SSOE, AA, AF>425 | <0.005 | 2 | 157.700273-157.700373 |  |  |
| 405 | Klimentidis 2017 | Genome variant | rs10930438 | AK023515 | Brain/muscle | MVPA, VPA, SSOE, AA, AF>425 | <0.005 | 2 | 170.750311-170.750411 |  |  |
| 406 | Klimentidis 2017 | Genome variant | rs1376935 | CADM2 | Brain/muscle | Strenuous Sports or Exercise: >2-3 vs 0 days/week | 1.28E-12 | 3 | 85.187225-85.187325 |  |  |
| 407 | Klimentidis 2017 | Genome variant | rs148854222 | Closest SLC1A3 | Brain/muscle | MVPA | 2.87E-08 | 5 | 36.505792-36.505892 | 15 | 8.801521-8.801540 |
| 408 | Klimentidis 2017 | Genome variant | rs2113077 | Closest ISL1 | Brain/muscle | MVPA, VPA, SSOE, AA, AF>425 | <0.005 | 5 | 51.503558-51.503658 |  |  |
| 409 | Klimentidis 2017 | Genome variant | rs3129981 | Closest HCG20 | Brain/muscle | MVPA | 5.13E-09 | 6 | 30.79103-30.79113 | 17 | 35.768708-35.768720 |
| 410 | Klimentidis 2017 | Genome variant | rs6909774 | MMS22L (MIR548H3) | Brain/muscle | Vigorous PA: >3 vs 0 days/week | 3.34E-08 | 6 | 97.239545-97.239645 |  |  |
| 411 | Klimentidis 2017 | Genome variant | rs10946808 | HIST1H1D | Brain/muscle | Strenuous Sports or Exercise: >2-3 vs 0 days/week | 3.10E-09 | 6 | 26.233109-26.233209 | 13 | 22.042846-22.042866 |
| 412 | Klimentidis 2017 | Genome variant | rs35622985 | MMS22L | Brain/muscle | Strenuous Sports or Exercise: >2-3 vs 0 days/week | 8.81E-09 | 6 | 97.335873-97.335973 |  |  |
| 413 | Klimentidis 2017 | Genome variant | rs1043595 | CALU | Brain/muscle | MVPA | 1.48E-09 | 7 | 128.769908-128.770008 | 6 | 29.375300-29.375316 |
| 414 | Klimentidis 2017 | Genome variant | rs921917 | Closest C7orf72 | Brain/muscle | MVPA | 1.19E-08 | 7 | 50.189092-50.189192 |  |  |
| 415 | Klimentidis 2017 | Genome variant | rs6955240 | EXOC4 | Brain/muscle | Vigorous PA: >3 vs 0 days/week | 9.57E-10 | 7 | 133.89707-133.89717 |  |  |
| 416 | Klimentidis 2017 | Genome variant | rs1993246 | KCCAT333 | Brain/muscle | MVPA, VPA, SSOE, AA, AF>425 | <0.005 | 7 | 17.437503-17.437603 |  |  |
| 417 | Klimentidis 2017 | Genome variant | rs124672 | KCNK9 | Brain/muscle | MVPA | 2.22E-08 | 8 | 139.709885-139.709985 | 15 | 72.553049-72.553060 |
| 418 | Klimentidis 2017 | Genome variant | rs72737787 | ZCCHC7 | Brain/muscle | MVPA | 3.66E-10 | 9 | 37.336694-37.336794 | 4 | 44.917425-44.917445 |
| 419 | Klimentidis 2017 | Genome variant | rs13284832 | MAPKAP1 | Brain/muscle | MVPA, VPA, SSOE, AA, AF>425 | <0.005 | 9 | 125.672797-125.672897 |  |  |
| 420 | Klimentidis 2017 | Genome variant | rs3781411 | CTBP2 | Brain/muscle | Vigorous PA: >3 vs 0 days/week | 4.88E-09 | 10 | 125.026817-125.026917 | 7 | 133.014320-133.014340 |
| 421 | Klimentidis 2017 | Genome variant | rs4747438 | DNAJC1 | Brain/muscle | Accelerometry - Avg Acceleration | 4.62E-08 | 10 | 21.835284-21.835384 |  |  |
| 422 | Klimentidis 2017 | Genome variant | rs7910002 | DNAJC1 | Brain/muscle | MVPA, VPA, SSOE, AA, AF>425 | <0.005 | 10 | 21.761591-21.761691 |  |  |
| 423 | Klimentidis 2017 | Genome variant | rs4411372 | STK24 | Brain/muscle | Strenuous Sports or Exercise: >2-3 vs 0 days/week | 2.51E-08 | 13 | 98.478119-98.478219 |  |  |
| 424 | Klimentidis 2017 | Genome variant | rs9579775 | ZMYM2 | Brain/muscle | MVPA, VPA, SSOE, AA, AF>425 | <0.005 | 13 | 20.042367-20.042467 |  |  |
| 425 | Klimentidis 2017 | Genome variant | rs12147808 | Closest C14orf64 | Brain/muscle | MVPA | 1.30E-08 | 14 | 98.152318-98.152418 | 12 | 107.100043-107.100062 |
| 426 | Klimentidis 2017 | Genome variant | rs1959759 | DCAF5 | Brain/muscle | Strenuous Sports or Exercise: >2-3 vs 0 days/week | 1.62E-08 | 14 | 69.16611-69.16621 |  |  |
| 427 | Klimentidis 2017 | Genome variant | rs10135643 | DCAF5 | Brain/muscle | MVPA, VPA, SSOE, AA, AF>425 | <0.005 | 14 | 69.050639-69.050739 | 12 | 80.335612-80.335632 |
| 428 | Klimentidis 2017 | Genome variant | rs10145335 | Closest C14ord177 | Brain/muscle | MVPA, VPA, SSOE, AA, AF>425 | <0.005 | 14 | 98.081361-98.081461 |  |  |
| 429 | Klimentidis 2017 | Genome variant | rs10851869 | PML | Brain/muscle | Accelerometry - Fraction Acceleration | 3.66E-08 | 15 | 74.038692-74.038792 | 9 | 58.225031-58.225051 |
| 430 | Klimentidis 2017 | Genome variant | rs5742915 | PML | Brain/muscle | MVPA, VPA, SSOE, AA, AF>425 | <0.005 | 15 | 74.044242-74.044342 | 9 | 58.220491-58.220508 |
| 431 | Klimentidis 2017 | Genome variant | rs1638525 | AKAP10 | Brain/muscle | Strenuous Sports or Exercise: >2-3 vs 0 days/week | 7.90E-09 | 17 | 19.945231-19.945331 |  |  |
| 432 | Klimentidis 2017 | Genome variant | rs62055545 | MAPT-AS1 | Brain/muscle | Accelerometry - Avg Acceleration | 1.41E-09 | 17 | 45.887145-45.887245 |  |  |
| 433 | Klimentidis 2017 | Genome variant | rs185231044 | RHBDL3 | Brain/muscle | MVPA, VPA, SSOE, AA, AF>425 | <0.005 | 17 | 32.310917-32.311017 |  |  |
| 434 | Klimentidis 2017 | Genome variant | rs9949626 | LOC100131655 | Brain/muscle | Strenuous Sports or Exercise: >2-3 vs 0 days/week | 3.10E-09 | 18 | 76.797952-76.798052 |  |  |
| 435 | Klimentidis 2017 | Genome variant | rs113351744 | Closest LINC01029 | Brain/muscle | MVPA, VPA, SSOE, AA, AF>425 | <0.005 | 18 | 77.873408-77.873508 |  |  |
| 436 | Klimentidis 2017 | Genome variant | rs429358 | APOE | Brain/muscle | MVPA | 4.98E-11 | 19 | 44.908634-44.908734 | 7 | 19.696942-19.696962 |
| 437 | Klimentidis 2017 | Genome variant | rs12460611 | Closest CCNE1 | Brain/muscle | MVPA, VPA, SSOE, AA, AF>425 | <0.005 | 19 | 29.835643-29.835743 |  |  |
| 438 | Klimentidis 2017 | Genome variant | rs1921981 | Closest BACE2 | Brain/muscle | MVPA | 4.03E-09 | 21 | 41.05057-41.05067 |  |  |
| 439 | Kostrzewa 2014 | Genome variant | rs459465 | DOK5 | Brain/muscle | "Excessive Exercise" (>5hrs/week) | 0.001 | 20 | 54.806433-54.806533 | 2 | 171.006365-171.006384 |
| 440 | Kostrzewa 2015 | Genome variant | rs6022999 | CYP24A1 | Brain/muscle | "Excessive Exercise" (>5hrs/week) | 0.011 | 20 | 54.171424-54.171524 | 2 | 170.495207-170.495000 |
| 441 | Kostrzewa 2016 | Genome variant | rs6092090 | DOK5 | Brain/muscle | "Excessive Exercise" (>5hrs/week) | 0.012 | 20 | 55.215067-55.215167 | 2 | 171.334832-171.334852 |
| 442 | Lin 2018 | Genome variant | rs116550874 | ENO1 | Brain/muscle | Leisure-Time Physical Activity | <1.0E-05 | 1 | 8.880557-8.880657 | 4 | 150.234745-150.234765 |
| 443 | Lin 2018 | Genome variant | rs7650960 | Closest CASR | Brain/muscle | Leisure-Time Physical Activity | <0.005 | 3 | 122.157407-122.157507 |  |  |
| 444 | Lin 2018 | Genome variant | rs112909877 | Closest CASR | Brain/muscle | Leisure-Time Physical Activity | <0.005 | 3 | 122.16509-122.16519 |  |  |
| 445 | Lin 2018 | Genome variant | rs146555373 | CASR | Brain/muscle | Leisure-Time Physical Activity | <0.005 | 3 | 122.266769-122.266869 |  |  |
| 446 | Lin 2018 | Genome variant | rs55716378 | Closest CASR | Brain/muscle | Leisure-Time Physical Activity | <0.005 | 3 | 122.304061-122.304161 |  |  |
| 447 | Lin 2018 | Genome variant | rs3792874 | SLC22A4 | Brain/muscle | Leisure-Time Physical Activity | <1.0E-05 | 5 | 132.301232-132.301332 | 11 | 54.023180-54.023187 |
| 448 | Lin 2018 | Genome variant | rs3792877 | SLC22A4 | Brain/muscle | Leisure-Time Physical Activity | <1.0E-05 | 5 | 132.301659-132.301759 | 11 | 54.022797-54.022817 |
| 449 | Lin 2018 | Genome variant | rs3792878 | SLC22A4 | Brain/muscle | Leisure-Time Physical Activity | <1.0E-05 | 5 | 132.302361-132.302461 | 11 | 54.021421-54.021430 |
| 450 | Lin 2018 | Genome variant | rs79173796 | SLC22A4 | Brain/muscle | Leisure-Time Physical Activity | <1.0E-05 | 5 | 132.298945-132.299045 | 11 | 54.024916-54.024939 |
| 451 | Lin 2018 | Genome variant | rs1819162 | ATAD1 (PAPSS2) | Brain/muscle | Leisure-Time Physical Activity | <0.005 | 10 | 87.767403-87.767503 |  |  |
| 452 | Lin 2018 | Genome variant | rs28524846 | MPP5 | Brain/muscle | Leisure-Time Physical Activity | <1.0E-05 | 14 | 67.329417-67.329517 |  |  |
| 453 | Lin 2018 | Genome variant | rs72707657 | GABRG3 | Brain/muscle | Leisure-Time Physical Activity | <0.005 | 15 | 27.480301-27.480401 | 7 | 56.773629-56.773650 |
| 454 | Lin 2018 | Genome variant | rs12438610 | GABRA5 | Brain/muscle | Leisure-Time Physical Activity | <0.005 | 15 | 26.944571-26.944671 |  |  |
| 455 | Lin 2018 | Genome variant | rs12902711 | GABRG3 | Brain/muscle | Leisure-Time Physical Activity | <0.005 | 15 | 26.977627-26.977727 |  |  |
| 456 | Lin 2018 | Genome variant | rs62020072 | Closest CYP19A1 | Brain/muscle | Leisure-Time Physical Activity | <0.005 | 15 | 51.183405-51.183505 |  |  |
| 457 | Lin 2018 | Genome variant | rs12595253 | GABRG3 | Brain/muscle | Leisure-Time Physical Activity | <0.005 | 15 | 27.067093-27.067193 |  |  |
| 458 | Maestu 2013 | Genome variant | rs1799752 | ACE | Brain/muscle | Light Physical Activity (min/d) | <0.05 | 17 | 63.48848-63.488593 | 11 | 105.979037-105.979071 |
| 459 | Pistilli 2011 | Genome variant | rs2228059 | IL15RA | Brain/muscle | Endurance vs Sprint Athelete Status | <0.05 | 10 | 5.960355-5.960455 |  |  |
| 460 | Stefan 2002 | Genome variant | PAL-Stefan | LEPR | Brain/muscle | "Physical Activity Level" | <0.05 |  | 0-0 |  |  |
| 461 | Wilkinson 2013 | Genome variant | rs6454672 | CNR1 | Brain/muscle | Decreased Liklihood to Meet Guidelines | 0.004 | 6 | 88.151801-88.151901 | 4 | 33.938182-33.938200 |
| 462 | Wilkinson 2013 | Genome variant | rs11615016 | TPH2 | Brain/muscle | Increased Liklihood to Meet Guidelines | 0.007 | 12 | 72.022164-72.022264 | 10 | 115.091095-115.091116 |
| 463 | Wilkinson 2013 | Genome variant | rs8066276 | ACE | Brain/muscle | Increased Liklihood to Meet Guidelines | 0.003 | 17 | 63.511854-63.511954 |  |  |
| 464 | Wilkinson 2013 | Genome variant | rs363035 | SNAP25 | Brain/muscle | Decreased Liklihood to Meet Guidelines | 0.004 | 20 | 10.23826-10.23836 |  |  |
| 465 | Kenney 2012 | Protein | Cannabinoid receptor-1 | CNR1 | Brain | Decreased wheel-running | 0.0007 females; 0.0025 males | 4 | 33.924593-33.948831 |  |  |
| 466 | Kenney 2008 | Protein | Cannabinoid receptor-1 | CNR1 | Brain | Decreased wheel-running | 0.0001 females; 0.05 males | 4 | 33.924593-33.948831 |  |  |
| 467 | Hillis 2020 | Genomic variant | Sortillin-related receptor, L | Sorl1 | Brain | High-runner mice (wheel-running) |  | 9 | 41.240184 – 42.275833 |  |  |
| 468 | Hillis 2020 | Genomic variant | Dachshund homolog 1 | Dach1 | Brain/muscle | High-runner mice (wheel-running) |  | 14 | 97.645171-98.679965 |  |  |
| 469 | Hillis 2020 | Genomic variant | Cadherin 10 | Cdh10 | Brain | High-runner mice (wheel-running) |  | 15 | 18.960135-20.609074 |  |  |

Chrm = Chromosome; NA = Not available
