## Supplemental Table 2: Mapped molecules in database. for "“It All Rolls Downstream: Upstream Control of Physical Activity Regulation”"

**Supplemental Table 2: Mapped Molecules Associated with Target Molecules in Dataset**

| **BRAIN MAPPED MOLECULES** | | | | |
| --- | --- | --- | --- | --- |
| **ID of molecule from literature (see Suppl Table 1)** | **Symbol of molecule from IPA database** | **Entrez Gene Name** | **Cellular Location** | **Type of molecule** |
| rs30737566 | 2610027F03Rik |  | Other | other |
| rs30737566 | 2610027F03Rik |  | Other | other |
| 5031434O11Rik | 5031434O11Rik | RIKEN cDNA 5031434O11 gene | Other | other |
| rs48305016 | 8030423F21Rik | RIKEN cDNA 8030423F21 gene | Other | other |
| rs48305016 | 8030423F21Rik | RIKEN cDNA 8030423F21 gene | Other | other |
| rs30462460 | 9330158H04Rik | RIKEN cDNA 9330158H04 gene | Other | other |
| rs30462460 | 9330158H04Rik | RIKEN cDNA 9330158H04 gene | Other | other |
| rs30462460 | 9330158H04Rik | RIKEN cDNA 9330158H04 gene | Other | other |
| Abhd12 | ABHD12 | abhydrolase domain containing 12, lysophospholipase | Plasma Membrane | enzyme |
| rs1799752 | ACE | angiotensin I converting enzyme | Plasma Membrane | peptidase |
| Acp1 | ACP1 | acid phosphatase 1 | Cytoplasm | phosphatase |
| Actn2 | ACTN2 | actinin alpha 2 | Nucleus | transcription regulator |
| rs30405098 | ACTR1A | actin related protein 1A | Cytoplasm | other |
| rs50154491 | ADAMTS10 | ADAM metallopeptidase with thrombospondin type 1 motif 10 | Extracellular Space | peptidase |
| Adora2a | ADORA2A | adenosine A2a receptor | Plasma Membrane | G-protein coupled receptor |
| rs26945355 | AFF4 | AF4/FMR2 family member 4 | Nucleus | transcription regulator |
| rs26945355 | AFF4 | AF4/FMR2 family member 4 | Nucleus | transcription regulator |
| rs26945355 | AFF4 | AF4/FMR2 family member 4 | Nucleus | transcription regulator |
| Ahr | AHR | aryl hydrocarbon receptor | Nucleus | ligand-dependent nuclear receptor |
| rs1638525 | AKAP10 | A-kinase anchoring protein 10 | Cytoplasm | other |
| Amn | AMN | amnion associated transmembrane protein | Plasma Membrane | other |
| Ankrd63 | ANKRD63 | ankyrin repeat domain 63 | Other | other |
| Anxa2 | ANXA2 | annexin A2 | Plasma Membrane | other |
| rs6259521 | APBA1 | amyloid beta precursor protein binding family A member 1 | Cytoplasm | transporter |
| rs429358 | APOE | apolipoprotein E | Extracellular Space | transporter |
| Arhgap8 | ARHGAP8/PRR5-ARHGAP8 | Rho GTPase activating protein 8 | Cytoplasm | other |
| Arrdc4 | ARRDC4 | arrestin domain containing 4 | Plasma Membrane | other |
| rs1819162 | ATAD1 | ATPase family AAA domain containing 1 | Plasma Membrane | enzyme |
| P50516 | ATP6V1A | ATPase H+ transporting V1 subunit A | Plasma Membrane | transporter |
| Baz1a | BAZ1A | bromodomain adjacent to zinc finger domain 1A | Nucleus | other |
| rs1376935 | CADM2 | cell adhesion molecule 2 | Plasma Membrane | other |
| rs34858520 | CALN1 | calneuron 1 | Cytoplasm | other |
| rs1043595 | CALU | calumenin | Cytoplasm | other |
| rs705692 | CAMTA1 | calmodulin binding transcription activator 1 | Other | other |
| Casq1 | CASQ1 | calsequestrin 1 | Cytoplasm | other |
| rs146555373 | CASR | calcium sensing receptor | Plasma Membrane | G-protein coupled receptor |
| rs13480409 | CCDC51 | coiled-coil domain containing 51 | Cytoplasm | transporter |
| rs13480409 | CCDC51 | coiled-coil domain containing 51 | Cytoplasm | transporter |
| rs13480409 | CCDC51 | coiled-coil domain containing 51 | Cytoplasm | transporter |
| rs13480409 | CCDC51 | coiled-coil domain containing 51 | Cytoplasm | transporter |
| CDH10 | CDH10 | cadherin 10 | Plasma Membrane | other |
| Chrm1 | CHRM1 | cholinergic receptor muscarinic 1 | Plasma Membrane | G-protein coupled receptor |
| rs6329684 | CHRM3 | cholinergic receptor muscarinic 3 | Plasma Membrane | G-protein coupled receptor |
| rs6329684 | CHRM3 | cholinergic receptor muscarinic 3 | Plasma Membrane | G-protein coupled receptor |
| rs6329684 | CHRM3 | cholinergic receptor muscarinic 3 | Plasma Membrane | G-protein coupled receptor |
| rs6329684 | CHRM3 | cholinergic receptor muscarinic 3 | Plasma Membrane | G-protein coupled receptor |
| rs6329684 | CHRM3 | cholinergic receptor muscarinic 3 | Plasma Membrane | G-protein coupled receptor |
| rs6329684 | CHRM3 | cholinergic receptor muscarinic 3 | Plasma Membrane | G-protein coupled receptor |
| rs6329684 | CHRM3 | cholinergic receptor muscarinic 3 | Plasma Membrane | G-protein coupled receptor |
| Q04447 | CKB | creatine kinase B | Cytoplasm | kinase |
| Clasp1 | CLASP1 | cytoplasmic linker associated protein 1 | Cytoplasm | other |
| P47746 | CNR1 | cannabinoid receptor 1 | Plasma Membrane | G-protein coupled receptor |
| rs6454672 | CNR1 | cannabinoid receptor 1 | Plasma Membrane | G-protein coupled receptor |
| CNR1 | CNR1 | cannabinoid receptor 1 | Plasma Membrane | G-protein coupled receptor |
| CNR1 | CNR1 | cannabinoid receptor 1 | Plasma Membrane | G-protein coupled receptor |
| rs30981553 | CNTNAP2 | contactin associated protein 2 | Plasma Membrane | other |
| Cpne4 | CPNE4 | copine 4 | Cytoplasm | other |
| Cpne5 | CPNE5 | copine 5 | Plasma Membrane | other |
| Crhbp | CRHBP | corticotropin releasing hormone binding protein | Extracellular Space | other |
| rs3781411 | CTBP2 | C-terminal binding protein 2 | Nucleus | transcription regulator |
| Ctxn1 | CTXN1 | cortexin 1 | Other | other |
| rs13476352 | CUBN | cubilin | Plasma Membrane | transmembrane receptor |
| rs13476352 | CUBN | cubilin | Plasma Membrane | transmembrane receptor |
| rs2942357 | CUBN | cubilin | Plasma Membrane | transmembrane receptor |
| rs6022999 | CYP24A1 | cytochrome P450 family 24 subfamily A member 1 | Cytoplasm | enzyme |
| Cyp2d22 | Cyp2d22 | cytochrome P450, family 2, subfamily d, polypeptide 22 | Cytoplasm | enzyme |
| Cyp4f15 | CYP4F8 | cytochrome P450 family 4 subfamily F member 8 | Cytoplasm | enzyme |
| D430019H16Rik | D430019H16Rik | RIKEN cDNA D430019H16 gene | Other | other |
| DACH1 | DACH1 | dachshund family transcription factor 1 | Nucleus | transcription regulator |
| rs10135643 | DCAF5 | DDB1 and CUL4 associated factor 5 | Cytoplasm | other |
| Ddn | DDN | dendrin | Cytoplasm | transcription regulator |
| Dlx1 | DLX1 | distal-less homeobox 1 | Nucleus | transcription regulator |
| Dlx2 | DLX2 | distal-less homeobox 2 | Nucleus | transcription regulator |
| rs34517439 | DNAJB4 | DnaJ heat shock protein family (Hsp40) member B4 | Nucleus | other |
| rs4747438 | DNAJC1 | DnaJ heat shock protein family (Hsp40) member C1 | Cytoplasm | other |
| rs7910002 | DNAJC1 | DnaJ heat shock protein family (Hsp40) member C1 | Cytoplasm | other |
| Q9EQF5 | DPYS | dihydropyrimidinase | Cytoplasm | enzyme |
| Q61616 | DRD1 | dopamine receptor D1 | Plasma Membrane | G-protein coupled receptor |
| Drd1 | DRD1 | dopamine receptor D1 | Plasma Membrane | G-protein coupled receptor |
| Drd2 | DRD2 | dopamine receptor D2 | Plasma Membrane | G-protein coupled receptor |
| rs13476818 | DTD1 | D-aminoacyl-tRNA deacylase 1 | Cytoplasm | enzyme |
| rs33245012 | EFL1 | elongation factor like GTPase 1 | Cytoplasm | translation regulator |
| rs25981 | EFNA5 | ephrin A5 | Plasma Membrane | kinase |
| Egr3 | EGR3 | early growth response 3 | Nucleus | transcription regulator |
| rs4136370 | ELAVL2 | ELAV like RNA binding protein 2 | Cytoplasm | other |
| rs116550874 | ENO1 | enolase 1 | Cytoplasm | enzyme |
| Ensa | ENSA | endosulfine alpha | Cytoplasm | transporter |
| Epha4 | EPHA4 | EPH receptor A4 | Plasma Membrane | kinase |
| rs31691517 | ERC1 | ELKS/RAB6-interacting/CAST family member 1 | Cytoplasm | other |
| rs6955240 | EXOC4 | exocyst complex component 4 | Cytoplasm | transporter |
| Fam107a | FAM107A | family with sequence similarity 107 member A | Nucleus | other |
| Foxg1 | FOXG1 | forkhead box G1 | Nucleus | transcription regulator |
| rs28032050 | FSIP2 | fibrous sheath interacting protein 2 | Cytoplasm | other |
| rs12438610 | GABRA5 | gamma-aminobutyric acid type A receptor subunit alpha5 | Plasma Membrane | ion channel |
| rs72707657 | GABRG3 | gamma-aminobutyric acid type A receptor subunit gamma3 | Plasma Membrane | ion channel |
| rs12902711 | GABRG3 | gamma-aminobutyric acid type A receptor subunit gamma3 | Plasma Membrane | ion channel |
| rs12595253 | GABRG3 | gamma-aminobutyric acid type A receptor subunit gamma3 | Plasma Membrane | ion channel |
| Gak | GAK | cyclin G associated kinase | Nucleus | kinase |
| rs49863494 | GALNTL6 | polypeptide N-acetylgalactosaminyltransferase like 6 | Other | enzyme |
| Gda | GDA | guanine deaminase | Cytoplasm | enzyme |
| Glo1 | GLO1 | glyoxalase I | Cytoplasm | enzyme |
| rs27368335 | Gm31473 | predicted gene, 31473 | Other | other |
| rs27368335 | Gm31473 | predicted gene, 31473 | Other | other |
| rs3675993 | Gm34282 | predicted gene, 34282 | Other | other |
| rs13478156 | Gm35223 | predicted gene, 35223 | Other | other |
| rs30945756 | Gm38475 | predicted gene, 38475 | Other | other |
| rs32432654 | Gm40890 | predicted gene, 40890 | Other | other |
| rs3671328 | Gm4952 | predicted gene 4952 | Cytoplasm | other |
| Gpr3 | GPR3 | G protein-coupled receptor 3 | Plasma Membrane | G-protein coupled receptor |
| Gpr34 | GPR34 | G protein-coupled receptor 34 | Plasma Membrane | G-protein coupled receptor |
| Gpr6 | GPR6 | G protein-coupled receptor 6 | Plasma Membrane | G-protein coupled receptor |
| Gpr88 | GPR88 | G protein-coupled receptor 88 | Plasma Membrane | G-protein coupled receptor |
| Gstm2 | GSTM1 | glutathione S-transferase mu 1 | Cytoplasm | enzyme |
| H1f0 | H1-0 | H1.0 linker histone | Nucleus | other |
| rs3129981 | HCG20 | HLA complex group 20 | Other | other |
| P38647 | HSPA9 | heat shock protein family A (Hsp70) member 9 | Cytoplasm | other |
| Htr1b | HTR1B | 5-hydroxytryptamine receptor 1B | Plasma Membrane | G-protein coupled receptor |
| P97288 | HTR4 | 5-hydroxytryptamine receptor 4 | Plasma Membrane | G-protein coupled receptor |
| Htra2 | HTRA2 | HtrA serine peptidase 2 | Cytoplasm | peptidase |
| Icam5 | ICAM5 | intercellular adhesion molecule 5 | Plasma Membrane | other |
| rs1051393 | IFNAR2 | interferon alpha and beta receptor subunit 2 | Plasma Membrane | transmembrane receptor |
| IL15 | IL15 | interleukin 15 | Extracellular Space | cytokine |
| rs46617906 | IQGAP2 | IQ motif containing GTPase activating protein 2 | Cytoplasm | other |
| Kcnf1 | KCNF1 | potassium voltage-gated channel modifier subfamily F member 1 | Plasma Membrane | ion channel |
| Kcng1 | KCNG1 | potassium voltage-gated channel modifier subfamily G member 1 | Plasma Membrane | ion channel |
| Kcnh4 | KCNH4 | potassium voltage-gated channel subfamily H member 4 | Plasma Membrane | ion channel |
| Kcnj4 | KCNJ4 | potassium inwardly rectifying channel subfamily J member 4 | Plasma Membrane | ion channel |
| Kcnj4 | KCNJ4 | potassium inwardly rectifying channel subfamily J member 4 | Plasma Membrane | ion channel |
| Kcnv1 | KCNV1 | potassium voltage-gated channel modifier subfamily V member 1 | Plasma Membrane | ion channel |
| Kif3a | KIF3A | kinesin family member 3A | Cytoplasm | enzyme |
| rs30816406 | KLHL12 | kelch like family member 12 | Cytoplasm | other |
| rs47227633 | KSR2 | kinase suppressor of ras 2 | Cytoplasm | kinase |
| Lamp5 | LAMP5 | lysosomal associated membrane protein family member 5 | Cytoplasm | other |
| Lgals1 | LGALS1 | galectin 1 | Extracellular Space | other |
| rs10930438 | LOC101926913 | uncharacterized LOC101926913 | Other | other |
| rs35622985 | LOC101927314 | uncharacterized LOC101927314 | Other | other |
| rs10145335 | LOC105370655 | uncharacterized LOC105370655 | Other | other |
| Lrrc10b | LRRC10B | leucine rich repeat containing 10B | Other | other |
| rs30241278 | MAP2 | microtubule associated protein 2 | Plasma Membrane | other |
| rs13284832 | MAPKAP1 | MAPK associated protein 1 | Cytoplasm | other |
| rs62055545 | MAPT-AS1 | MAPT antisense RNA 1 | Other | other |
| rs62020072 | MIR4713HG | MIR4713 host gene | Other | other |
| rs6909774 | MMS22L | MMS22 like, DNA repair protein | Nucleus | other |
| Mstn | MSTN | myostatin | Extracellular Space | growth factor |
| Neurl1b | NEURL1B | neuralized E3 ubiquitin protein ligase 1B | Cytoplasm | enzyme |
| Npw | NPW | neuropeptide W | Extracellular Space | other |
| Nrgn | Nrgn | neurogranin | Plasma Membrane | other |
| Nts | NTS | neurotensin | Extracellular Space | other |
| Ociad2 | OCIAD2 | OCIA domain containing 2 | Cytoplasm | other |
| rs10887741 | PAPSS2 | 3'-phosphoadenosine 5'-phosphosulfate synthase 2 | Cytoplasm | enzyme |
| rs1980647 | PAPSS2 | 3'-phosphoadenosine 5'-phosphosulfate synthase 2 | Cytoplasm | enzyme |
| rs4934355 | PAPSS2 | 3'-phosphoadenosine 5'-phosphosulfate synthase 2 | Cytoplasm | enzyme |
| rs1358864 | PAPSS2 | 3'-phosphoadenosine 5'-phosphosulfate synthase 2 | Cytoplasm | enzyme |
| rs7903568 | PAPSS2 | 3'-phosphoadenosine 5'-phosphosulfate synthase 2 | Cytoplasm | enzyme |
| rs2077695 | PAPSS2 | 3'-phosphoadenosine 5'-phosphosulfate synthase 2 | Cytoplasm | enzyme |
| rs10218939 | PAPSS2 | 3'-phosphoadenosine 5'-phosphosulfate synthase 2 | Cytoplasm | enzyme |
| rs11202501 | PAPSS2 | 3'-phosphoadenosine 5'-phosphosulfate synthase 2 | Cytoplasm | enzyme |
| rs12412482 | PAPSS2 | 3'-phosphoadenosine 5'-phosphosulfate synthase 2 | Cytoplasm | enzyme |
| rs7908056 | PAPSS2 | 3'-phosphoadenosine 5'-phosphosulfate synthase 2 | Cytoplasm | enzyme |
| Pde9a | PDE9A | phosphodiesterase 9A | Cytoplasm | enzyme |
| Phlda3 | PHLDA3 | pleckstrin homology like domain family A member 3 | Plasma Membrane | other |
| Pla2g7 | PLA2G7 | phospholipase A2 group VII | Extracellular Space | enzyme |
| rs10851869 | PML | PML nuclear body scaffold | Nucleus | transcription regulator |
| rs5742915 | PML | PML nuclear body scaffold | Nucleus | transcription regulator |
| rs2267668 | PPARD | peroxisome proliferator activated receptor delta | Nucleus | ligand-dependent nuclear receptor |
| rs2076168 | PPARD | peroxisome proliferator activated receptor delta | Nucleus | ligand-dependent nuclear receptor |
| rs8192678 | PPARGC1A | PPARG coactivator 1 alpha | Nucleus | transcription regulator |
| rs6361467 | Ppp2r5c | protein phosphatase 2, regulatory subunit B', gamma | Nucleus | phosphatase |
| rs3664044 | Ppp4r1l-ps | protein phosphatase 4, regulatory subunit 1-like, pseudogene | Other | other |
| Prcp | PRCP | prolylcarboxypeptidase | Cytoplasm | peptidase |
| rs31923186 | PRKCB | protein kinase C beta | Cytoplasm | kinase |
| Prss12 | PRSS12 | serine protease 12 | Extracellular Space | peptidase |
| Ptprv | Ptprv | protein tyrosine phosphatase, receptor type, V | Other | other |
| rs13478553 | PVRIG | PVR related immunoglobulin domain containing | Plasma Membrane | other |
| Rab40b | RAB40B | RAB40B, member RAS oncogene family | Plasma Membrane | enzyme |
| rs13482429 | RAI14 | retinoic acid induced 14 | Nucleus | transcription regulator |
| rs3677375 | RBBP5 | RB binding protein 5, histone lysine methyltransferase complex subunit | Nucleus | transcription regulator |
| rs13478157 | Rbm33 | RNA binding motif protein 33 | Other | other |
| rs185231044 | RHBDL3 | rhomboid like 3 | Plasma Membrane | peptidase |
| Rora | RORA | RAR related orphan receptor A | Nucleus | ligand-dependent nuclear receptor |
| Rprml | RPRML | reprimo like | Other | other |
| Rtn4rl2 | RTN4RL2 | reticulon 4 receptor like 2 | Plasma Membrane | other |
| Rxrg | RXRG | retinoid X receptor gamma | Nucleus | ligand-dependent nuclear receptor |
| Scd4 | Scd4 | stearoyl-coenzyme A desaturase 4 | Cytoplasm | enzyme |
| Q62420 | SH3GL2 | SH3 domain containing GRB2 like 2, endophilin A1 | Plasma Membrane | enzyme |
| rs3792874 | SLC22A4 | solute carrier family 22 member 4 | Plasma Membrane | transporter |
| rs3792877 | SLC22A4 | solute carrier family 22 member 4 | Plasma Membrane | transporter |
| rs3792878 | SLC22A4 | solute carrier family 22 member 4 | Plasma Membrane | transporter |
| rs79173796 | SLC22A4 | solute carrier family 22 member 4 | Plasma Membrane | transporter |
| Slc38a2 | SLC38A2 | solute carrier family 38 member 2 | Plasma Membrane | transporter |
| Mfsd7a | SLC49A3 | solute carrier family 49 member 3 | Other | other |
| Slco2a1 | SLCO2A1 | solute carrier organic anion transporter family member 2A1 | Plasma Membrane | transporter |
| rs363035 | SNAP25 | synaptosome associated protein 25 | Plasma Membrane | transporter |
| rs33841297 | SND1 | staphylococcal nuclease and tudor domain containing 1 | Nucleus | enzyme |
| Snrnp25 | SNRNP25 | small nuclear ribonucleoprotein U11/U12 subunit 25 | Nucleus | other |
| SORL1 | SORL1 | sortilin related receptor 1 | Cytoplasm | transporter |
| rs3686133 | SP7 | Sp7 transcription factor | Nucleus | transcription regulator |
| Sp9 | SP9 | Sp9 transcription factor | Nucleus | transcription regulator |
| rs3659852 | SPOCK3 | SPARC (osteonectin), cwcv and kazal like domains proteoglycan 3 | Extracellular Space | other |
| Sst | SST | somatostatin | Extracellular Space | other |
| Sst | SST | somatostatin | Extracellular Space | other |
| rs4411372 | STK24 | serine/threonine kinase 24 | Cytoplasm | kinase |
| Q9WUM5 | SUCLG1 | succinate-CoA ligase GDP/ADP-forming subunit alpha | Cytoplasm | enzyme |
| rs3715009 | SUSD1 | sushi domain containing 1 | Other | other |
| rs28225821 | SYNRG | synergin gamma | Cytoplasm | other |
| rs1858242 | TAFA1 | TAFA chemokine like family member 1 | Extracellular Space | other |
| Tbr1 | TBR1 | T-box brain transcription factor 1 | Nucleus | transcription regulator |
| P10711 | TCEA1 | transcription elongation factor A1 | Nucleus | transcription regulator |
| P24529 | TH | tyrosine hydroxylase | Cytoplasm | enzyme |
| Thra | THRA | thyroid hormone receptor alpha | Nucleus | ligand-dependent nuclear receptor |
| Tln2 | TLN2 | talin 2 | Nucleus | other |
| Tmed5 | TMED5 | transmembrane p24 trafficking protein 5 | Cytoplasm | other |
| Tmub2 | TMUB2 | transmembrane and ubiquitin like domain containing 2 | Other | other |
| Tmub2 | TMUB2 | transmembrane and ubiquitin like domain containing 2 | Other | other |
| rs11615016 | TPH2 | tryptophan hydroxylase 2 | Plasma Membrane | enzyme |
| Ugt1a6a | UGT1A6 | UDP glucuronosyltransferase family 1 member A6 | Cytoplasm | enzyme |
| Vdr | VDR | vitamin D receptor | Nucleus | transcription regulator |
| rs181053839 | WDPCP | WD repeat containing planar cell polarity effector | Plasma Membrane | other |
| Wipf3 | WIPF3 | WAS/WASL interacting protein family member 3 | Plasma Membrane | other |
| rs31675929 | XKR4 | XK related 4 | Plasma Membrane | other |
| rs72737787 | ZCCHC7 | zinc finger CCHC-type containing 7 | Nucleus | other |
| rs9579775 | ZMYM2 | zinc finger MYM-type containing 2 | Nucleus | transcription regulator |
| rs9949626 | ZNF236-DT | ZNF236 divergent transcript | Other | other |
| rs13481855 | ZNF274 | zinc finger protein 274 | Nucleus | transcription regulator |
| rs13481855 | ZNF274 | zinc finger protein 274 | Nucleus | transcription regulator |
| Zfp831 | ZNF831 | zinc finger protein 831 | Other | other |
| **MUSCLE MAPPED MOLECULES** | | | | |
| **ID of molecule from literature (see Suppl Table 1)** | **Symbol of molecule from IPA database** | **Entrez Gene Name** | **Cellular Location** | **Type of molecule** |
| rs30737566 | 2610027F03Rik |  | Other | other |
| rs30737566 | 2610027F03Rik |  | Other | other |
| rs48305016 | 8030423F21Rik | RIKEN cDNA 8030423F21 gene | Other | other |
| rs48305016 | 8030423F21Rik | RIKEN cDNA 8030423F21 gene | Other | other |
| rs30462460 | 9330158H04Rik | RIKEN cDNA 9330158H04 gene | Other | other |
| rs30462460 | 9330158H04Rik | RIKEN cDNA 9330158H04 gene | Other | other |
| rs30462460 | 9330158H04Rik | RIKEN cDNA 9330158H04 gene | Other | other |
| Q8BGQ7 | AARS1 | alanyl-tRNA synthetase 1 | Cytoplasm | enzyme |
| Acadvl | ACADVL | acyl-CoA dehydrogenase very long chain | Cytoplasm | enzyme |
| rs1799752 | ACE | angiotensin I converting enzyme | Plasma Membrane | peptidase |
| Acot8 | ACOT8 | acyl-CoA thioesterase 8 | Cytoplasm | enzyme |
| Acp1 | ACP1 | acid phosphatase 1 | Cytoplasm | phosphatase |
| Q9JI91 | ACTN2 | actinin alpha 2 | Nucleus | transcription regulator |
| rs30405098 | ACTR1A | actin related protein 1A | Cytoplasm | other |
| rs50154491 | ADAMTS10 | ADAM metallopeptidase with thrombospondin type 1 motif 10 | Extracellular Space | peptidase |
| rs26945355 | AFF4 | AF4/FMR2 family member 4 | Nucleus | transcription regulator |
| rs26945355 | AFF4 | AF4/FMR2 family member 4 | Nucleus | transcription regulator |
| rs26945355 | AFF4 | AF4/FMR2 family member 4 | Nucleus | transcription regulator |
| Ahr | AHR | aryl hydrocarbon receptor | Nucleus | ligand-dependent nuclear receptor |
| P29699 | AHSG | alpha 2-HS glycoprotein | Extracellular Space | other |
| rs1638525 | AKAP10 | A-kinase anchoring protein 10 | Cytoplasm | other |
| P24549 | ALDH1A1 | aldehyde dehydrogenase 1 family member A1 | Cytoplasm | enzyme |
| P97429 | ANXA4 | annexin A4 | Plasma Membrane | other |
| P48036 | ANXA5 | annexin A5 | Plasma Membrane | transporter |
| P14824 | ANXA6 | annexin A6 | Plasma Membrane | ion channel |
| P14824 | ANXA6 | annexin A6 | Plasma Membrane | ion channel |
| rs6259521 | APBA1 | amyloid beta precursor protein binding family A member 1 | Cytoplasm | transporter |
| Q00623 | APOA1 | apolipoprotein A1 | Extracellular Space | transporter |
| Q00623 | APOA1 | apolipoprotein A1 | Extracellular Space | transporter |
| rs429358 | APOE | apolipoprotein E | Extracellular Space | transporter |
| rs1819162 | ATAD1 | ATPase family AAA domain containing 1 | Plasma Membrane | enzyme |
| P56480 | ATP5F1B | ATP synthase F1 subunit beta | Cytoplasm | transporter |
| P50516 | ATP6V1A | ATPase H+ transporting V1 subunit A | Plasma Membrane | transporter |
| AK082735 | C230098O21Rik | RIKEN cDNA C230098O21 gene | Other | other |
| Car14 | CA14 | carbonic anhydrase 14 | Plasma Membrane | enzyme |
| rs1376935 | CADM2 | cell adhesion molecule 2 | Plasma Membrane | other |
| rs34858520 | CALN1 | calneuron 1 | Cytoplasm | other |
| rs1043595 | CALU | calumenin | Cytoplasm | other |
| rs705692 | CAMTA1 | calmodulin binding transcription activator 1 | Other | other |
| O09165 | CASQ1 | calsequestrin 1 | Cytoplasm | other |
| Casq1 | CASQ1 | calsequestrin 1 | Cytoplasm | other |
| rs146555373 | CASR | calcium sensing receptor | Plasma Membrane | G-protein coupled receptor |
| rs13480409 | CCDC51 | coiled-coil domain containing 51 | Cytoplasm | transporter |
| rs13480409 | CCDC51 | coiled-coil domain containing 51 | Cytoplasm | transporter |
| rs13480409 | CCDC51 | coiled-coil domain containing 51 | Cytoplasm | transporter |
| rs13480409 | CCDC51 | coiled-coil domain containing 51 | Cytoplasm | transporter |
| P80317 | CCT6A | chaperonin containing TCP1 subunit 6A | Cytoplasm | other |
| P45591 | CFL2 | cofilin 2 | Extracellular Space | other |
| rs6329684 | CHRM3 | cholinergic receptor muscarinic 3 | Plasma Membrane | G-protein coupled receptor |
| rs6329684 | CHRM3 | cholinergic receptor muscarinic 3 | Plasma Membrane | G-protein coupled receptor |
| rs6329684 | CHRM3 | cholinergic receptor muscarinic 3 | Plasma Membrane | G-protein coupled receptor |
| rs6329684 | CHRM3 | cholinergic receptor muscarinic 3 | Plasma Membrane | G-protein coupled receptor |
| rs6329684 | CHRM3 | cholinergic receptor muscarinic 3 | Plasma Membrane | G-protein coupled receptor |
| rs6329684 | CHRM3 | cholinergic receptor muscarinic 3 | Plasma Membrane | G-protein coupled receptor |
| rs6329684 | CHRM3 | cholinergic receptor muscarinic 3 | Plasma Membrane | G-protein coupled receptor |
| rs6454672 | CNR1 | cannabinoid receptor 1 | Plasma Membrane | G-protein coupled receptor |
| rs30981553 | CNTNAP2 | contactin associated protein 2 | Plasma Membrane | other |
| Q8K1Z0 | COQ9 | coenzyme Q9 | Cytoplasm | other |
| rs3781411 | CTBP2 | C-terminal binding protein 2 | Nucleus | transcription regulator |
| rs13476352 | CUBN | cubilin | Plasma Membrane | transmembrane receptor |
| rs13476352 | CUBN | cubilin | Plasma Membrane | transmembrane receptor |
| rs2942357 | CUBN | cubilin | Plasma Membrane | transmembrane receptor |
| rs6022999 | CYP24A1 | cytochrome P450 family 24 subfamily A member 1 | Cytoplasm | enzyme |
| rs10135643 | DCAF5 | DDB1 and CUL4 associated factor 5 | Cytoplasm | other |
| rs34517439 | DNAJB4 | DnaJ heat shock protein family (Hsp40) member B4 | Nucleus | other |
| rs4747438 | DNAJC1 | DnaJ heat shock protein family (Hsp40) member C1 | Cytoplasm | other |
| rs7910002 | DNAJC1 | DnaJ heat shock protein family (Hsp40) member C1 | Cytoplasm | other |
| rs13476818 | DTD1 | D-aminoacyl-tRNA deacylase 1 | Cytoplasm | enzyme |
| Eepd1 | EEPD1 | endonuclease/exonuclease/phosphatase family domain containing 1 | Plasma Membrane | other |
| rs33245012 | EFL1 | elongation factor like GTPase 1 | Cytoplasm | translation regulator |
| rs25981 | EFNA5 | ephrin A5 | Plasma Membrane | kinase |
| rs4136370 | ELAVL2 | ELAV like RNA binding protein 2 | Cytoplasm | other |
| rs116550874 | ENO1 | enolase 1 | Cytoplasm | enzyme |
| P34914 | EPHX2 | epoxide hydrolase 2 | Cytoplasm | enzyme |
| rs31691517 | ERC1 | ELKS/RAB6-interacting/CAST family member 1 | Cytoplasm | other |
| Q921G7 | ETFDH | electron transfer flavoprotein dehydrogenase | Cytoplasm | enzyme |
| rs6955240 | EXOC4 | exocyst complex component 4 | Cytoplasm | transporter |
| P30416 | FKBP4 | FKBP prolyl isomerase 4 | Nucleus | enzyme |
| rs28032050 | FSIP2 | fibrous sheath interacting protein 2 | Cytoplasm | other |
| rs12438610 | GABRA5 | gamma-aminobutyric acid type A receptor subunit alpha5 | Plasma Membrane | ion channel |
| rs72707657 | GABRG3 | gamma-aminobutyric acid type A receptor subunit gamma3 | Plasma Membrane | ion channel |
| rs12902711 | GABRG3 | gamma-aminobutyric acid type A receptor subunit gamma3 | Plasma Membrane | ion channel |
| rs12595253 | GABRG3 | gamma-aminobutyric acid type A receptor subunit gamma3 | Plasma Membrane | ion channel |
| rs49863494 | GALNTL6 | polypeptide N-acetylgalactosaminyltransferase like 6 | Other | enzyme |
| rs27368335 | Gm31473 | predicted gene, 31473 | Other | other |
| rs27368335 | Gm31473 | predicted gene, 31473 | Other | other |
| rs3675993 | Gm34282 | predicted gene, 34282 | Other | other |
| rs13478156 | Gm35223 | predicted gene, 35223 | Other | other |
| rs30945756 | Gm38475 | predicted gene, 38475 | Other | other |
| rs32432654 | Gm40890 | predicted gene, 40890 | Other | other |
| rs3671328 | Gm4952 | predicted gene 4952 | Cytoplasm | other |
| rs3129981 | HCG20 | HLA complex group 20 | Other | other |
| Q91X72 | HPX | hemopexin | Extracellular Space | transporter |
| Ifi30 | IFI30 | IFI30 lysosomal thiol reductase | Cytoplasm | enzyme |
| rs1051393 | IFNAR2 | interferon alpha and beta receptor subunit 2 | Plasma Membrane | transmembrane receptor |
| Q60819 | IL15RA | interleukin 15 receptor subunit alpha | Plasma Membrane | transmembrane receptor |
| Ilf3 | ILF3 | interleukin enhancer binding factor 3 | Nucleus | transcription regulator |
| rs46617906 | IQGAP2 | IQ motif containing GTPase activating protein 2 | Cytoplasm | other |
| rs30816406 | KLHL12 | kelch like family member 12 | Cytoplasm | other |
| rs47227633 | KSR2 | kinase suppressor of ras 2 | Cytoplasm | kinase |
| rs10930438 | LOC101926913 | uncharacterized LOC101926913 | Other | other |
| rs35622985 | LOC101927314 | uncharacterized LOC101927314 | Other | other |
| rs10145335 | LOC105370655 | uncharacterized LOC105370655 | Other | other |
| P51885 | LUM | lumican | Extracellular Space | other |
| rs30241278 | MAP2 | microtubule associated protein 2 | Plasma Membrane | other |
| rs13284832 | MAPKAP1 | MAPK associated protein 1 | Cytoplasm | other |
| rs62055545 | MAPT-AS1 | MAPT antisense RNA 1 | Other | other |
| mmu-miR-497 | miR-16-5p (and other miRNAs w/seed AGCAGCA) |  | Cytoplasm | mature microRNA |
| mmu-miR-195 | miR-16-5p (and other miRNAs w/seed AGCAGCA) |  | Cytoplasm | mature microRNA |
| mmu-miR-324-3P | miR-324-3p (and other miRNAs w/seed CACUGCC) |  | Cytoplasm | mature microRNA |
| rs62020072 | MIR4713HG | MIR4713 host gene | Other | other |
| rs6909774 | MMS22L | MMS22 like, DNA repair protein | Nucleus | other |
| Mstn | MSTN | myostatin | Extracellular Space | growth factor |
| Mt1 | Mt1 | metallothionein 1 | Cytoplasm | other |
| Mtmr14 | MTMR14 | myotubularin related protein 14 | Cytoplasm | phosphatase |
| Mup2 | Mup1 (includes others) | major urinary protein 1 | Extracellular Space | other |
| Myh10 | MYH10 | myosin heavy chain 10 | Cytoplasm | enzyme |
| P09542 | MYL3 | myosin light chain 3 | Cytoplasm | other |
| Q9DCT2 | NDUFS3 | NADH:ubiquinone oxidoreductase core subunit S3 | Cytoplasm | enzyme |
| Nus1 | NUS1 | NUS1 dehydrodolichyl diphosphate synthase subunit | Cytoplasm | enzyme |
| Q60597 | OGDH | oxoglutarate dehydrogenase | Cytoplasm | enzyme |
| rs10887741 | PAPSS2 | 3'-phosphoadenosine 5'-phosphosulfate synthase 2 | Cytoplasm | enzyme |
| rs1980647 | PAPSS2 | 3'-phosphoadenosine 5'-phosphosulfate synthase 2 | Cytoplasm | enzyme |
| rs4934355 | PAPSS2 | 3'-phosphoadenosine 5'-phosphosulfate synthase 2 | Cytoplasm | enzyme |
| rs1358864 | PAPSS2 | 3'-phosphoadenosine 5'-phosphosulfate synthase 2 | Cytoplasm | enzyme |
| rs7903568 | PAPSS2 | 3'-phosphoadenosine 5'-phosphosulfate synthase 2 | Cytoplasm | enzyme |
| rs2077695 | PAPSS2 | 3'-phosphoadenosine 5'-phosphosulfate synthase 2 | Cytoplasm | enzyme |
| rs10218939 | PAPSS2 | 3'-phosphoadenosine 5'-phosphosulfate synthase 2 | Cytoplasm | enzyme |
| rs11202501 | PAPSS2 | 3'-phosphoadenosine 5'-phosphosulfate synthase 2 | Cytoplasm | enzyme |
| rs12412482 | PAPSS2 | 3'-phosphoadenosine 5'-phosphosulfate synthase 2 | Cytoplasm | enzyme |
| rs7908056 | PAPSS2 | 3'-phosphoadenosine 5'-phosphosulfate synthase 2 | Cytoplasm | enzyme |
| Q9Z2V4 | PCK1 | phosphoenolpyruvate carboxykinase 1 | Cytoplasm | kinase |
| Pcyt2 | PCYT2 | phosphate cytidylyltransferase 2, ethanolamine | Cytoplasm | enzyme |
| Q9D051 | PDHB | pyruvate dehydrogenase E1 subunit beta | Cytoplasm | enzyme |
| Q8BKZ9 | PDHX | pyruvate dehydrogenase complex component X | Cytoplasm | enzyme |
| Q3UV70 | PDP1 | pyruvate dehydrogenase phosphatase catalytic subunit 1 | Cytoplasm | phosphatase |
| P70266 | PFKFB1 | 6-phosphofructo-2-kinase/fructose-2,6-biphosphatase 1 | Cytoplasm | kinase |
| rs10851869 | PML | PML nuclear body scaffold | Nucleus | transcription regulator |
| rs5742915 | PML | PML nuclear body scaffold | Nucleus | transcription regulator |
| Q9Z2M7 | PMM2 | phosphomannomutase 2 | Cytoplasm | enzyme |
| rs2267668 | PPARD | peroxisome proliferator activated receptor delta | Nucleus | ligand-dependent nuclear receptor |
| rs2076168 | PPARD | peroxisome proliferator activated receptor delta | Nucleus | ligand-dependent nuclear receptor |
| rs8192678 | PPARGC1A | PPARG coactivator 1 alpha | Nucleus | transcription regulator |
| rs6361467 | Ppp2r5c | protein phosphatase 2, regulatory subunit B', gamma | Nucleus | phosphatase |
| rs3664044 | Ppp4r1l-ps | protein phosphatase 4, regulatory subunit 1-like, pseudogene | Other | other |
| Prcp | PRCP | prolylcarboxypeptidase | Cytoplasm | peptidase |
| O08709 | PRDX6 | peroxiredoxin 6 | Cytoplasm | enzyme |
| rs31923186 | PRKCB | protein kinase C beta | Cytoplasm | kinase |
| rs13478553 | PVRIG | PVR related immunoglobulin domain containing | Plasma Membrane | other |
| rs13482429 | RAI14 | retinoic acid induced 14 | Nucleus | transcription regulator |
| rs3677375 | RBBP5 | RB binding protein 5, histone lysine methyltransferase complex subunit | Nucleus | transcription regulator |
| rs13478157 | Rbm33 | RNA binding motif protein 33 | Other | other |
| P26043 | RDX | radixin | Cytoplasm | other |
| rs185231044 | RHBDL3 | rhomboid like 3 | Plasma Membrane | peptidase |
| Rpl13a | RPL13A | ribosomal protein L13a | Cytoplasm | other |
| S100a1 | S100A1 | S100 calcium binding protein A1 | Cytoplasm | other |
| Q8K2B3 | SDHA | succinate dehydrogenase complex flavoprotein subunit A | Cytoplasm | enzyme |
| Q00897 | SERPINA1 | serpin family A member 1 | Extracellular Space | other |
| Q00896 | SERPINA1 | serpin family A member 1 | Extracellular Space | other |
| P07759 | SERPINA3 | serpin family A member 3 | Extracellular Space | other |
| Sh3kbp1 | SH3KBP1 | SH3 domain containing kinase binding protein 1 | Cytoplasm | other |
| Q8BRU6 | SLC18A2 | solute carrier family 18 member A2 | Plasma Membrane | transporter |
| rs3792874 | SLC22A4 | solute carrier family 22 member 4 | Plasma Membrane | transporter |
| rs3792877 | SLC22A4 | solute carrier family 22 member 4 | Plasma Membrane | transporter |
| rs3792878 | SLC22A4 | solute carrier family 22 member 4 | Plasma Membrane | transporter |
| rs79173796 | SLC22A4 | solute carrier family 22 member 4 | Plasma Membrane | transporter |
| P14142 | SLC2A4 | solute carrier family 2 member 4 | Plasma Membrane | transporter |
| Slc2a4 | SLC2A4 | solute carrier family 2 member 4 | Plasma Membrane | transporter |
| rs363035 | SNAP25 | synaptosome associated protein 25 | Plasma Membrane | transporter |
| rs33841297 | SND1 | staphylococcal nuclease and tudor domain containing 1 | Nucleus | enzyme |
| rs3686133 | SP7 | Sp7 transcription factor | Nucleus | transcription regulator |
| Sparc | SPARC | secreted protein acidic and cysteine rich | Extracellular Space | other |
| rs3659852 | SPOCK3 | SPARC (osteonectin), cwcv and kazal like domains proteoglycan 3 | Extracellular Space | other |
| Q7TQ48 | SRL | sarcalumenin | Cytoplasm | other |
| rs4411372 | STK24 | serine/threonine kinase 24 | Cytoplasm | kinase |
| rs3715009 | SUSD1 | sushi domain containing 1 | Other | other |
| rs28225821 | SYNRG | synergin gamma | Cytoplasm | other |
| rs1858242 | TAFA1 | TAFA chemokine like family member 1 | Extracellular Space | other |
| Q921I1 | TF | transferrin | Extracellular Space | transporter |
| Trp53 | TP53 | tumor protein p53 | Nucleus | transcription regulator |
| rs11615016 | TPH2 | tryptophan hydroxylase 2 | Plasma Membrane | enzyme |
| Q1XH17 | TRIM72 | tripartite motif containing 72 | Cytoplasm | enzyme |
| Q9ERD7 | TUBB3 | tubulin beta 3 class III | Cytoplasm | other |
| Q9JMH6 | TXNRD1 | thioredoxin reductase 1 | Cytoplasm | enzyme |
| Q62465 | VAT1 | vesicle amine transport 1 | Plasma Membrane | transporter |
| Q64727 | VCL | vinculin | Plasma Membrane | enzyme |
| P20152 | VIM | vimentin | Cytoplasm | other |
| rs181053839 | WDPCP | WD repeat containing planar cell polarity effector | Plasma Membrane | other |
| rs31675929 | XKR4 | XK related 4 | Plasma Membrane | other |
| P68254 | YWHAQ | tyrosine 3-monooxygenase/tryptophan 5-monooxygenase activation protein theta | Cytoplasm | other |
| rs72737787 | ZCCHC7 | zinc finger CCHC-type containing 7 | Nucleus | other |
| rs9579775 | ZMYM2 | zinc finger MYM-type containing 2 | Nucleus | transcription regulator |
| rs9949626 | ZNF236-DT | ZNF236 divergent transcript | Other | other |
| rs13481855 | ZNF274 | zinc finger protein 274 | Nucleus | transcription regulator |
| rs13481855 | ZNF274 | zinc finger protein 274 | Nucleus | transcription regulator |

Mapped molecules = molecules arising from IPA dataset associated with target molecules from literature. To find source of target molecules, take symbol of molecule from IPA database (from any figures or table), locate it in column 2 above, find the associated identification of the molecule from the literature (column 1), and find that identified molecule in Supplemental Table 1.
