## Supplemental Table 3: All upstream regulators of brain molecules for "“It All Rolls Downstream: Upstream Control of Physical Activity Regulation”"

**Supplemental Table 3: Upstream Regulators of Brain Target Molecules**

| **Upstream Regulator** | **Molecule Type** | **p-value of overlap** | **# of Molecules** | **Target Molecules in Dataset** |
| --- | --- | --- | --- | --- |
| beta-estradiol | chemical - endogenous mammalian | 4.8E-08 | 44 | ACE, ADORA2A, AFF4, AHR, AKAP10, ANXA2, APBA1, APOE, BAZ1A, CHRM1, CHRM3, CKB, CNR1, CUBN, CYP24A1, CYP4F8, DNAJC1, DRD1, DRD2, EGR3, ENO1, FAM107A, FOXG1, GABRG3, GPR88, GSTM1, HSPA9, HTR4, ICAM5, IL15, IQGAP2, NPW, NTS, PAPSS2, PPARGC1A, PRKCB, SLC38A2, SNAP25, SST, TH, THRA, TLN2, TPH2, VDR |
| lipopolysaccharide | chemical drug | 0.00349 | 30 | ACE,ADORA2A,AHR,ANXA2,APOE,CASR,CUBN,DDN,EGR3,ENO1, FOXG1, GPR3, GPR34, IFNAR2, IL15, LGALS1, NTS, PAPSS2, PLA2G7, PML, PPARD, PPARGC1A, PRKCB, RORA, RXRG, SLC38A2, SLCO2A1, TH, THRA, VDR |
| dexamethasone | chemical drug | 0.0086 | 27 | ACE,APOE,CRHBP,CYP24A1,DACH1,EGR3,ENO1,FAM107A,FOXG1,GPR34, GSTM1, HSPA9, IFNAR2, IL15, LGALS1, MAP2, MSTN, NTS, PPARD, PPARGC1A, PRKCB, RORA, SLCO2A1, SST, THRA, UGT1A6, VDR |
| tretinoin | chemical - endogenous mammalian | 0.000839 | 25 | AHR,ANXA2,APOE,CNR1,CNTNAP2,CUBN,DRD2,IL15,LGALS1,MAP2, NRGN, PA2G7, PML, PPARD, PPARGC1A, PRKCB, RAI14, RXRG, SLC38A2, SCLO2A1, SP7,SST,TH,THRA,VDR |
| TP53 | transcription regulator | 0.00488 | 25 | ACE,ANXA2,APOE,CALU,CDH10,CKB,CYP24A1,DLX1,DNAJB4,EGR3, ENSA, GDA, GSTM1, KCNG1, KCNJ4, PHLDA3, PML, PPARD, PPARGC1A, PRKCB, PTPRV, SP7, SUCLG1, THRA, VDR |
| HTT | transcription regulator | 4.83E-10 | 24 | ACTN2, ADORA2A, APOE, CNR1, CPNE5, DRD1, DRD2, GLO1, GPR88, HSPA9, HTRA2, IL15, KCNJ4, MAP2, NTS, PPARD, PPARGC1A, PRKCB, RXRG, SNAP25, SST, TBR1, TH, THRA |
| TNF | cytokine | 0.0365 | 22 | ACE,ADORA2A,APOE,CLASP1,DLX1,DPYS,EGR3,GABRA5,GPR88,IFNAR2 |
| calcitriol | chemical drug | 1.2E-06 | 18 | AHR,ANXA2,ATAD1,CASR,CYP24A1,EFL1,FAM107A,HSPA9,IL15,KSR2 |
| CREB1 | transcription regulator | 2.34E-07 | 18 | ADORA2A,APOE,CALN1,CDH10,CRHBP,GABRA5,GDA,GPR3,NEURL1B,Nrgn |
| IL4 | cytokine | 0.00418 | 18 | ADORA2A,AHR,ANXA2,APOE,CKB,EFL1,H1-0,HTRA2,IFNAR2,IL15 |
| IFNG | cytokine | 0.0348 | 18 | ACE,ADORA2A,AHR,CNR1,CYP24A1,EGR3,ELAVL2,ENO1,FAM107A,IL15 |
| APP | other | 0.000956 | 17 | ACTN2,APOE,ATP6V1A,CKB,DRD2,ENO1,ENSA,GPR6,IL15,MAP2 |
| tetradecanoylphorbol acetate | chemical drug | 0.00745 | 17 | ACE,ADORA2A,CRHBP,CYP24A1,EGR3,EPHA4,GPR34,GSTM1,NTS,PPARD |
| SNCA | enzyme | 6.07E-08 | 16 | APOE,CRHBP,CTXN1,DDN,FAM107A,GDA,GLO1,ICAM5,KCNF1,KCNG1 |
| BDNF | growth factor | 2.01E-07 | 14 | ANXA2,CASQ1,CNR1,DRD2,EGR3,GDA,LAMP5,LGALS1,PRSS12,SNAP25 |
| levodopa | chemical - endogenous mammalian | 0.000433 | 14 | ADORA2A,CNR1,CTXN1,EFNA5,GSTM1,HTR1B,IFNAR2,MAPKAP1,RHBDL3,SH3GL2 |
| MAPT | other | 0.000075 | 14 | ACTR1A,ATP6V1A,CKB,DRD2,ENO1,ENSA,GLO1,GPR6,MAP2,SH3GL2 |
| decitabine | chemical drug | 0.00888 | 13 | ANXA2,APOE,CNTNAP2,CTBP2,CYP24A1,EGR3,HSPA9,HTR1B,LGALS1,MAP2 |
| Immunoglobulin | complex | 0.0286 | 13 | AHR,APOE,CYP4F8,EGR3,ENO1,FAM107A,GPR34,HTRA2,IL15,LGALS1 |
| D-glucose | chemical - endogenous mammalian | 0.00359 | 13 | ACE,CNR1,CYP24A1,DRD1,ENO1,KCNF1,LGALS1,NTS,PPARD,PPARGC1A |
| forskolin | chemical toxicant | 0.00347 | 13 | ACE,ADORA2A,ATP6V1A,CRHBP,EPHA4,GPR3,OCIAD2,PPARGC1A,SNAP25,SST |
| PSEN1 | peptidase | 3.57E-05 | 12 | APOE,ATP6V1A,CKB,ENO1,ENSA,GPR6,MAP2,SH3GL2,SLC38A2,SNAP25 |
| JAK1/2 | group | 3.79E-12 | 11 | CTXN1,DDN,FAM107A,GDA,ICAM5,KCNF1,KCNG1,Nrgn,RPRML,TBR1 |
| ESR2 | ligand-dependent nuclear receptor | 0.0295 | 11 | APOE,ARRDC4,CALU,CHRM1,DNAJC1,EGR3,ELAVL2,H1-0,IQGAP2,SLC38A2 |
| FOS | transcription regulator | 0.00383 | 11 | AFF4,CALU,DRD1,ENO1,GAK,NTS,OCIAD2,Ppp2r5c,TAFA1,TCEA1 |
| LEP | growth factor | 0.00114 | 11 | ACE,CYP24A1,DRD2,KIF3A,NPW,NTS,PPARGC1A,Scd4,SNAP25,SST |
| progesterone | chemical - endogenous mammalian | 0.0124 | 11 | ACE,APOE,CNR1,DNAJB4,DRD2,IL15,KCNG1,LGALS1,PLA2G7,TH |
| GLI1 | transcription regulator | 0.00607 | 10 | EGR3,ELAVL2,GDA,H1-0,IQGAP2,LGALS1,NEURL1B,PAPSS2,PPARGC1A,SP7 |
| SP1 | transcription regulator | 0.0111 | 10 | APOE,CKB,DRD1,DRD2,IL15,PPARD,PRKCB,SLC22A4,SNAP25,VDR |
| Insulin | group | 0.0352 | 10 | APOE,CHRM1,CHRM3,DNAJB4,HSPA9,PPARD,PPARGC1A,PRKCB,SLC38A2,SNAP25 |
| IL15 | cytokine | 0.00228 | 10 | AHR,APBA1,ENO1,ICAM5,IL15,PPARD,PPARGC1A,SORL1,TCEA1,VDR |
| PD98059 | chemical - kinase inhibitor | 0.011 | 10 | ACE,CYP24A1,EGR3,ENO1,MSTN,PPARD,SLC38A2,SP7,TH,VDR |
| CEBPB | transcription regulator | 0.0369 | 9 | CYP24A1,MMS22L,PPARD,PPARGC1A,RORA,SLC38A2,SST,UGT1A6,VDR |
| U0126 | chemical drug | 0.0264 | 9 | CYP24A1,EGR3,GPR34,PML,PPARD,PRKCB,RXRG,SP7,TH |
| ethanol | chemical - endogenous mammalian | 0.00623 | 9 | CNR1,DRD1,DRD2,IL15,LGALS1,MAP2,Nrgn,PPARGC1A,RORA |
| bexarotene | chemical drug | 1.35E-05 | 9 | APOE,DACH1,DRD1,EFNA5,EGR3,IL15,NTS,TBR1,TH |
| NFkB (complex) | complex | 0.0493 | 9 | AHR,APOE,DRD2,GPR34,HSPA9,IFNAR2,IL15,MSTN,SLC22A4 |
| dopamine | chemical - endogenous mammalian | 4.95E-05 | 8 | CNR1,CYP4F8,DRD1,DRD2,EGR3,IL15,MAP2,TH |
| RXRA | ligand-dependent nuclear receptor | 0.002 | 8 | APOE,CYP24A1,DRD2,PPARD,RORA,RXRG,TH,VDR |
| LY294002 | chemical drug | 0.0471 | 8 | APOE,CNR1,CUBN,EGR3,MAP2,NTS,PPARD,RXRG |
| SOD1 | enzyme | 0.00113 | 8 | APOE,CKB,CNR1,GABRG3,GAK,GSTM1,IL15,SP7 |
| CREBBP | transcription regulator | 0.0101 | 8 | ADORA2A,CKB,CYP24A1,DNAJB4,EGR3,PPARGC1A,SST,TH |
| PPARD | ligand-dependent nuclear receptor | 0.00167 | 8 | ADORA2A,APOE,ENO1,NPW,PPARD,PPARGC1A,RORA,SP7 |
| Ca2+ | chemical - endogenous mammalian | 0.00167 | 8 | ADORA2A,AHR,CASR,KCNG1,PPARGC1A,PRSS12,SST,VDR |
| GnRH analog | biologic drug | 0.00317 | 8 | ACP1,AKAP10,ENSA,GPR3,MAP2,PRCP,TCEA1,THRA |
| 8-bromo-cAMP | chemical reagent | 0.0382 | 8 | ACE,CKB,DRD2,IL15,IQGAP2,PPARD,PPARGC1A,TLN2 |
| FGF2 | growth factor | 0.0331 | 8 | ACE,CHRM3,ENO1,EPHA4,RORA,SP7,SST,TH |
| VDR | transcription regulator | 0.000777 | 8 | ACE,CASR,CUBN,CYP24A1,KSR2,LGALS1,TPH2,VDR |
| Vegf | group | 0.04 | 8 | ACE,ADORA2A,AHR,EGR3,EPHA4,PPARD,PPARGC1A,PRKCB |
| ASCL1 | transcription regulator | 6.13E-06 | 7 | DLX1,DLX2,FOXG1,LAMP5,SNAP25,TBR1,TH |
| tazemetostat | chemical drug | 0.00883 | 7 | CNTNAP2,CTBP2,HSPA9,LGALS1,MAP2,TCEA1,ZMYM2 |
| SP2509 | chemical reagent | 0.00786 | 7 | CNTNAP2,CTBP2,HSPA9,LGALS1,MAP2,TCEA1,ZMYM2 |
| Creb | group | 0.0124 | 7 | CDH10,EGR3,GDA,NTS,PPARGC1A,SST,TH |
| tetrodotoxin | chemical drug | 7.92E-05 | 7 | CADM2,CDH10,CPNE4,CPNE5,GDA,PPARGC1A,SORL1 |
| Mek | group | 0.0136 | 7 | APOE,EGR3,GDA,IL15,PPARGC1A,SORL1,SP7 |
| MECP2 | transcription regulator | 0.00056 | 7 | APOE,EFNA5,FOXG1,GABRA5,KCNF1,TH,TPH2 |
| cholecalciferol | chemical - endogenous mammalian | 5.74E-05 | 7 | ANXA2,CUBN,CYP24A1,PPARGC1A,PRKCB,SP7,VDR |
| PDX1 | transcription regulator | 0.000577 | 7 | ANXA2,ARRDC4,CKB,MAP2,PAPSS2,SST,XKR4 |
| PDGF BB | complex | 0.0146 | 7 | AHR,EGR3,GDA,MAP2,PPARD,RXRG,TH |
| cycloheximide | chemical reagent | 0.0237 | 7 | AHR,CYP24A1,DRD2,KSR2,SST,THRA,VDR |
| glucocorticoid | chemical drug | 0.00722 | 7 | AHR,CNR1,CYP24A1,PPARGC1A,TH,TPH2,VDR |
| Z-LLL-CHO | chemical - protease inhibitor | 0.0172 | 7 | AHR,CASR,DRD2,ENO1,HSPA9,SP7,VDR |
| IgG | complex | 0.0117 | 7 | AHR,APOE,EGR3,IL15,LGALS1,PPARGC1A,STK24 |
| H89 | chemical drug | 0.000211 | 7 | ACE,ADORA2A,EGR3,NTS,PPARD,PPARGC1A,SNAP25 |
| SRF | transcription regulator | 0.0457 | 6 | EGR3,GDA,IL15,MSTN,SND1,ZMYM2 |
| methamphetamine | chemical drug | 0.00103 | 6 | DRD2,GDA,LGALS1,NTS,TH,VDR |
| ZFHX3 | transcription regulator | 5.08E-05 | 6 | DRD1,PLA2G7,RORA,SP9,SST,TBR1 |
| ESRRA | transcription regulator | 0.0146 | 6 | CKB,EGR3,ENO1,PPARGC1A,PRCP,THRA |
| aflatoxin B1 | chemical - endogenous non-mammalian | 0.0215 | 6 | CHRM3,ELAVL2,PHLDA3,THRA,UGT1A6,VDR |
| GRIN3A | ion channel | 0.000189 | 6 | CADM2,CDH10,CPNE4,CPNE5,GDA,SORL1 |
| LEPR | transmembrane receptor | 0.00309 | 6 | APOE,CNR1,EXOC4,PPARD,PPARGC1A,SNAP25 |
| CNR1 | G-protein coupled receptor | 0.000905 | 6 | APOE,CNR1,DRD2,GLO1,MSTN,RXRG |
| NGF | growth factor | 0.0132 | 6 | ANXA2,DRD2,EPHA4,MAP2,SNAP25,TH |
| NCOA2 | transcription regulator | 0.00142 | 6 | AHR,GLO1,GPR34,RORA,THRA,VDR |
| NR1I2 | ligand-dependent nuclear receptor | 0.00301 | 6 | AHR,CYP24A1,GSTM1,PAPSS2,SORL1,UGT1A6 |
| streptozocin | chemical drug | 0.0211 | 6 | ADORA2A,CASR,DRD1,PPARD,PPARGC1A,SST |
| cocaine | chemical drug | 0.00309 | 6 | ADORA2A,CASR,CNR1,DRD2,NTS,TH |
| LMNA | other | 0.0129 | 6 | ADAMTS10,ARRDC4,MAPKAP1,SLC38A2,TH,ZMYM2 |
| nicotine | chemical drug | 0.00714 | 6 | ACE,AHR,CHRM3,ENO1,PPARD,TH |
| SCD | enzyme | 0.0125 | 5 | PPARD,PPARGC1A,Scd4,THRA,VDR |
| fenofibrate | chemical drug | 0.0465 | 5 | MSTN,PPARGC1A,RORA,SUCLG1,THRA |
| HDAC4 | transcription regulator | 0.00409 | 5 | GABRA5,GABRG3,PRKCB,SH3GL2,SNAP25 |
| FMR1 | translation regulator | 0.0062 | 5 | EGR3,ENO1,GABRA5,ICAM5,SNAP25 |
| cyclic AMP | chemical - endogenous mammalian | 0.0484 | 5 | DRD1,IL15,PPARGC1A,SST,TH |
| GDNF | growth factor | 0.000181 | 5 | DRD1,DRD2,LGALS1,SST,TH |
| FGF8 | growth factor | 0.000655 | 5 | DLX1,DLX2,FOXG1,TH,VDR |
| ADCYAP1 | other | 0.0308 | 5 | CNTNAP2,EGR3,SNAP25,SST,UGT1A6 |
| CD 437 | chemical drug | 0.0172 | 5 | CNR1,ENO1,GPR6,KIF3A,LGALS1 |
| 6-hydroxydopamine | chemical toxicant | 0.000755 | 5 | CNR1,DRD2,HSPA9,SST,TH |
| Esrra | transcription regulator | 0.00161 | 5 | CNR1,DRD2,ENO1,PPARGC1A,THRA |
| PRKAG3 | kinase | 0.00741 | 5 | ATP6V1A,CLASP1,IL15,PHLDA3,PPARGC1A |
| MAPK8 | kinase | 0.0108 | 5 | APOE,GSTM1,PPARD,PPARGC1A,VDR |
| TO-901317 | chemical reagent | 0.0313 | 5 | APOE,ENO1,RAI14,Scd4,THRA |
| levothyroxine | chemical - endogenous mammalian | 0.000219 | 5 | APOE,DRD2,Nrgn,RORA,THRA |
| carbon tetrachloride | chemical toxicant | 0.0358 | 5 | APOE,CNR1,Cyp2d22,GSTM1,PPARD |
| kainic acid | chemical toxicant | 0.012 | 5 | APOE,CNR1,CUBN,EGR3,GABRA5 |
| FEV | transcription regulator | 0.000438 | 5 | ANXA2,EFNA5,NTS,SST,TPH2 |
| 1-methyl-4-phenyl-1,2,3,6-tetrahydropyridine | chemical toxicant | 0.00108 | 5 | ANXA2,CNR1,EGR3,MAP2,TH |
| methapyrilene | chemical drug | 0.00352 | 5 | ANXA2,APOE,GSTM1,PRKCB,RXRG |
| metformin | chemical drug | 0.0453 | 5 | AHR,GLO1,MSTN,PHLDA3,PPARGC1A |
| PRNP | other | 0.000314 | 5 | AHR,APOE,HSPA9,SNAP25,TH |
| FGF1 | growth factor | 0.00264 | 5 | AHR,APOE,DLX2,GSTM1,TH |
| BMP7 | growth factor | 0.0289 | 5 | ADORA2A,DRD2,PPARGC1A,SP7,THRA |
| isobutylmethylxanthine | chemical toxicant | 0.0218 | 5 | ACE,CRHBP,PPARD,PPARGC1A,TH |
| Nr1h | group | 0.00589 | 5 | ACE,APOE,IL15,PML,PPARGC1A |
| nitric oxide | chemical - endogenous mammalian | 0.00637 | 5 | ACE,AHR,GSTM1,PPARGC1A,RXRG |
| Pka catalytic subunit | group | 4.16E-05 | 4 | PPARGC1A,SST,TH,VDR |
| APC | enzyme | 0.0399 | 4 | PPARD,PRKCB,SST,SUCLG1 |
| ATF2 | transcription regulator | 0.00496 | 4 | NTS,PPARGC1A,SST,TH |
| Sb202190 | chemical drug | 0.0184 | 4 | MAPKAP1,PPARGC1A,SP7,VDR |
| L-glutamic acid | chemical - endogenous mammalian | 0.0188 | 4 | MAP2,PPARGC1A,Scd4,SORL1 |
| DSCAM | other | 0.00514 | 4 | KCNJ4,MAP2,NEURL1B,PRKCB |
| S100A9 | other | 0.0365 | 4 | HTR4,PRSS12,RXRG,UGT1A6 |
| S100A8 | other | 0.0442 | 4 | HTR4,PRSS12,RXRG,UGT1A6 |
| sulforafan | chemical drug | 0.0231 | 4 | GSTM1,MSTN,PML,UGT1A6 |
| LAMA4 | enzyme | 0.00616 | 4 | GSTM1,IFNAR2,IL15,TLN2 |
| HNRNPA2B1 | other | 0.0226 | 4 | GABRA5,NTS,PPARGC1A,SNAP25 |
| mono-(2-ethylhexyl)phthalate | chemical toxicant | 0.0352 | 4 | ENO1,PPARD,PPARGC1A,SUCLG1 |
| CLPP | peptidase | 0.0022 | 4 | ENO1,HSPA9,PPARGC1A,SUCLG1 |
| REST | transcription regulator | 0.0327 | 4 | EFNA5,SNAP25,TPH2,XKR4 |
| diphtheria toxin | chemical - endogenous non-mammalian | 0.0109 | 4 | DRD1,DRD2,LGALS1,PLA2G7 |
| PLX5622 | chemical drug | 0.0231 | 4 | DRD1,DRD2,IL15,TBR1 |
| MRTFA | transcription regulator | 0.048 | 4 | DNAJB4,EGR3,MAP2,SPOCK3 |
| CX3CL1 | cytokine | 0.0197 | 4 | DLX1,DLX2,SNAP25,TH |
| EGR2 | transcription regulator | 0.0308 | 4 | DDN,EPHA4,MAP2,RORA |
| pregnenolone carbonitrile | chemical drug | 0.000557 | 4 | CYP24A1,SLCO2A1,SORL1,UGT1A6 |
| alitretinoin | chemical drug | 0.0358 | 4 | CYP24A1,DRD2,PPARGC1A,RXRG |
| 1,25-dihydroxyvitamin D | chemical drug | 0.00137 | 4 | CUBN,CYP24A1,MSTN,VDR |
| Calmodulin | group | 0.000959 | 4 | CPNE5,CTXN1,ICAM5,RAB40B |
| morphine | chemical drug | 0.0236 | 4 | CNR1,DRD1,DRD2,TH |
| KLF11 | transcription regulator | 0.0159 | 4 | CHRM1,CHRM3,DRD2,PPARGC1A |
| topotecan | chemical drug | 0.0495 | 4 | CAMTA1,CNTNAP2,EGR3,ENO1 |
| RGS4 | enzyme | 0.00477 | 4 | CALN1,PPARGC1A,PRSS12,VDR |
| FLCN | other | 0.0167 | 4 | ATP6V1A,CKB,ENO1,PPARGC1A |
| Pka | complex | 0.0207 | 4 | APOE,NTS,PPARGC1A,SST |
| TFAP2A | transcription regulator | 0.0167 | 4 | APOE,DRD1,GLO1,PPARD |
| PML | transcription regulator | 0.0333 | 4 | APOE,DNAJB4,PML,PPARGC1A |
| Rxr | group | 0.0097 | 4 | APOE,CYP24A1,EFNA5,TPH2 |
| Ngf | group | 0.00199 | 4 | ANXA2,MAP2,PPARGC1A,TH |
| Pln | other | 0.00079 | 4 | ANXA2,CALU,ENO1,PRKCB |
| PLN | transporter | 0.00122 | 4 | ANXA2,CALU,ENO1,PRKCB |
| nitrofurantoin | chemical drug | 0.0333 | 4 | ANXA2,APOE,GSTM1,PRKCB |
| POR | enzyme | 0.0435 | 4 | AHR,GSTM1,LGALS1,PLA2G7 |
| Hdac | group | 0.0339 | 4 | AHR,CYP24A1,EGR3,PPARGC1A |
| TNFSF13B | cytokine | 0.0236 | 4 | ADORA2A,EFNA5,IL15,VDR |
| haloperidol | chemical drug | 0.00408 | 4 | ADORA2A,DRD2,NTS,TH |
| ZNF503 | other | 4.82E-05 | 4 | ADORA2A,DRD1,DRD2,GPR6 |
| SNAI1 | transcription regulator | 0.0308 | 4 | ACTR1A,RORA,SP7,VDR |
| MEF2C | transcription regulator | 0.0167 | 4 | ACTN2,CKB,PPARGC1A,SP7 |
| losartan potassium | chemical drug | 0.0221 | 4 | ACE,MAP2,PPARGC1A,TH |
| isoproterenol | chemical drug | 0.0385 | 4 | ACE,CYP4F8,GPR3,TH |
| CREM | transcription regulator | 0.0296 | 4 | ACE,APOE,SST,TH |
| thyroid hormone | chemical - endogenous mammalian | 0.0464 | 4 | ACE,APOE,PML,SST |
| POU4F1 | transcription regulator | 0.0474 | 3 | RTN4RL2,SNAP25,TH |
| NPY | other | 0.0119 | 3 | PPARGC1A,SST,TH |
| CHKB | kinase | 0.00456 | 3 | PPARD,PPARGC1A,RXRG |
| PIN1 | enzyme | 0.022 | 3 | PML,PPARD,PPARGC1A |
| JUND | transcription regulator | 0.0234 | 3 | NTS,SST,TH |
| phenylephrine | chemical drug | 0.0288 | 3 | MSTN,PRKCB,THRA |
| ascorbic acid | chemical - endogenous mammalian | 0.0404 | 3 | IL15,SP7,TH |
| Rhox5 | transcription regulator | 0.00146 | 3 | IFNAR2,PPARGC1A,SLC22A4 |
| fluoxetine | chemical drug | 0.0234 | 3 | HTR4,TH,TPH2 |
| PDLIM2 | other | 0.0313 | 3 | GSTM1,PHLDA3,SH3GL2 |
| TRAF2 | enzyme | 0.0234 | 3 | GPR34,PPARGC1A,SNRNP25 |
| DSCAML1 | other | 0.028 | 3 | GPR3,MAP2,PRKCB |
| 2,4,5,2',4',5'-hexachlorobiphenyl | chemical toxicant | 0.0474 | 3 | EXOC4,Ppp2r5c,SST |
| ELOVL3 | enzyme | 0.0145 | 3 | ENO1,IQGAP2,PLA2G7 |
| Hif1 | complex | 0.018 | 3 | ENO1,HTRA2,LGALS1 |
| peoniflorin | chemical drug | 0.000393 | 3 | ENO1,GLO1,SNAP25 |
| VIP | other | 0.0454 | 3 | EGR3,LGALS1,TH |
| sodium arsenite | chemical drug | 0.0145 | 3 | EGR3,ENO1,TH |
| DTNBP1 | other | 0.000393 | 3 | DRD2,SNAP25,SST |
| SLC6A3 | transporter | 0.000285 | 3 | DRD2,NTS,TH |
| amphetamine | chemical drug | 0.0249 | 3 | DRD2,NTS,TH |
| risperidone | chemical drug | 0.000972 | 3 | DRD2,NTS,PPARGC1A |
| clozapine | chemical drug | 0.00999 | 3 | DRD2,NTS,PPARGC1A |
| NEUROG3 | transcription regulator | 0.0433 | 3 | DRD1,NTS,SST |
| quinolinic acid | chemical - endogenous mammalian | 0.00514 | 3 | DRD1,DRD2,TH |
| GNA15 | enzyme | 0.0496 | 3 | DPYS,GDA,MAP2 |
| MYO6 | other | 0.0227 | 3 | DNAJB4,EGR3,ERC1 |
| TAF4 | transcription regulator | 0.033 | 3 | DLX2,DRD2,PPARGC1A |
| IHH | enzyme | 0.0322 | 3 | DLX1,ENO1,GPR34 |
| POU4F2 | transcription regulator | 0.00264 | 3 | DLX1,DLX2,TH |
| PPP4R3A | other | 0.000457 | 3 | DLX1,DLX2,SP9 |
| Rb | group | 0.0234 | 3 | DLX1,DLX2,PPARGC1A |
| phorbol esters | chemical - other | 0.0288 | 3 | CYP24A1,PRKCB,VDR |
| GW501516 | chemical drug | 0.0433 | 3 | CYP24A1,PPARD,PPARGC1A |
| triadimefon | chemical toxicant | 0.00577 | 3 | CYP24A1,GABRG3,UGT1A6 |
| propiconazole | chemical reagent | 0.00306 | 3 | CYP24A1,GABRG3,GSTM1 |
| seocalcitol | chemical drug | 0.0109 | 3 | CUBN,CYP24A1,VDR |
| PAX5-ELN | fusion gene/product | 0.00352 | 3 | CRHBP,EFNA5,HTR1B |
| 2-arachidonoylglycerol | chemical - endogenous mammalian | 0.000238 | 3 | CNR1,PPARD,PPARGC1A |
| NR4A2 | ligand-dependent nuclear receptor | 0.0322 | 3 | CNR1,DRD2,TH |
| sodium tungstate | chemical drug | 0.00176 | 3 | CKB,ENO1,SNAP25 |
| TFE3 | transcription regulator | 0.00999 | 3 | CKB,ENO1,PPARGC1A |
| SLC18A3 | transporter | 0.000197 | 3 | CHRM1,CHRM3,DRD2 |
| birabresib | chemical drug | 0.0119 | 3 | CDH10,PDE9A,RAI14 |
| vitamin D | chemical drug | 0.00999 | 3 | CASR,CYP24A1,VDR |
| VitaminD3-VDR-RXR | complex | 0.00456 | 3 | CASR,CYP24A1,PPARD |
| ASPSCR1-TFE3 | fusion gene/product | 0.0357 | 3 | APOE,KCNJ4,PPARGC1A |
| beta-naphthoflavone | chemical toxicant | 0.00609 | 3 | APOE,GSTM1,UGT1A6 |
| COP1 | enzyme | 0.0193 | 3 | APOE,EGR3,PPARGC1A |
| MBD2 | transcription regulator | 0.0124 | 3 | APOE,EFNA5,SST |
| MBD3 | other | 0.022 | 3 | APOE,EFNA5,PPARD |
| Sn50 peptide | chemical toxicant | 0.0114 | 3 | APOE,DRD2,IL15 |
| ZIC2 | transcription regulator | 3.07E-05 | 3 | APOE,DRD1,EPHA4 |
| verapamil | chemical drug | 0.00912 | 3 | ANXA2,CYP24A1,ENO1 |
| LONP1 | peptidase | 0.0206 | 3 | ANXA2,CALU,HSPA9 |
| plicamycin | chemical drug | 0.0474 | 3 | AHR,SNAP25,VDR |
| BCL11B | transcription regulator | 0.0151 | 3 | AHR,RORA,TBR1 |
| estradiol benzoate | chemical drug | 0.0213 | 3 | AHR,PPARD,SST |
| NR1I3 | ligand-dependent nuclear receptor | 0.0454 | 3 | AHR,GSTM1,PAPSS2 |
| CGS 21680 | chemical reagent | 0.00352 | 3 | ADORA2A,EGR3,PPARGC1A |
| MYL2 | other | 0.00226 | 3 | ACTN2,ANXA2,LGALS1 |
| propranolol | chemical drug | 0.00352 | 3 | ACE,PPARGC1A,TH |
| N-nitro-L-arginine methyl ester | chemical drug | 0.0385 | 3 | ACE,PPARD,PPARGC1A |
| okadaic acid | chemical toxicant | 0.0322 | 3 | ACE,CYP24A1,SP7 |
| MAPK3 | kinase | 0.0423 | 3 | ACE,CYP24A1,PPARGC1A |
| NR1H2 | ligand-dependent nuclear receptor | 0.0414 | 3 | ACE,APOE,THRA |
| ketoconazole | chemical drug | 0.0188 | 2 | TH,VDR |
| magnesium | chemical - endogenous mammalian | 0.0188 | 2 | TH,VDR |
| CRHR2 | G-protein coupled receptor | 0.012 | 2 | TH,TPH2 |
| citalopram | chemical drug | 0.00143 | 2 | TH,TPH2 |
| PRKAR1A | kinase | 0.0297 | 2 | SST,VDR |
| NTF3 | growth factor | 0.0164 | 2 | SST,TH |
| ABL1 | kinase | 0.0375 | 2 | SP7,SST |
| MAPK13 | kinase | 0.011 | 2 | PPARGC1A,VDR |
| MAPK11 | kinase | 0.0226 | 2 | PPARGC1A,VDR |
| MAPK12 | kinase | 0.011 | 2 | PPARGC1A,VDR |
| clenbuterol | chemical drug | 0.00745 | 2 | PPARGC1A,VDR |
| KISS1 | other | 0.0152 | 2 | PPARGC1A,TH |
| ADM2 | other | 0.0101 | 2 | PPARGC1A,TH |
| pCPT-cAMP | chemical - kinase inhibitor | 0.00666 | 2 | PPARGC1A,SST |
| PKNOX1 | transcription regulator | 0.0176 | 2 | PPARGC1A,SST |
| CHIR 99021 | chemical drug | 0.0327 | 2 | PPARGC1A,SP7 |
| agmatine | chemical - endogenous mammalian | 0.011 | 2 | PPARGC1A,SLC22A4 |
| PLIN5 | other | 0.0358 | 2 | PPARGC1A,Scd4 |
| TCF7 | transcription regulator | 0.0495 | 2 | PPARD,RORA |
| natriuretic peptide derivative | biologic drug | 0.000157 | 2 | PPARD,PPARGC1A |
| PLCL2 | enzyme | 0.00331 | 2 | PPARD,PPARGC1A |
| PRKD | group | 0.024 | 2 | PPARD,PPARGC1A |
| SERTAD2 | transcription regulator | 0.00519 | 2 | PPARD,PPARGC1A |
| PLCL1 | enzyme | 0.00666 | 2 | PPARD,PPARGC1A |
| guanidinopropionic acid | chemical - endogenous non-mammalian | 0.0495 | 2 | PPARD,PPARGC1A |
| UBE2I | enzyme | 0.0312 | 2 | PML,PPARGC1A |
| NTS | other | 0.011 | 2 | NTS,TH |
| chlorpyrifos | chemical toxicant | 0.00143 | 2 | NTS,TH |
| bromocriptine | chemical drug | 0.0342 | 2 | NTS,TH |
| ATF1 | transcription regulator | 0.0375 | 2 | NTS,SST |
| CDK5 | kinase | 0.0267 | 2 | MAP2,TH |
| pargyline | chemical drug | 0.00277 | 2 | MAP2,TH |
| 7-nitroindazole | chemical reagent | 0.00666 | 2 | MAP2,TH |
| CDK2 | kinase | 0.0442 | 2 | MAP2,SNAP25 |
| aroclor 1254 | chemical toxicant | 0.0059 | 2 | MAP2,Nrgn |
| SUB1 | transcription regulator | 0.0188 | 2 | KCNG1,KCNJ4 |
| CYP27B1 | enzyme | 0.0408 | 2 | IL15,VDR |
| TLR1 | transmembrane receptor | 0.00917 | 2 | IL15,VDR |
| NKX2-2 | transcription regulator | 0.00917 | 2 | IL15,SST |
| HHEX | transcription regulator | 0.0253 | 2 | IL15,SST |
| iodoacetic acid | chemical toxicant | 0.00389 | 2 | IL15,RORA |
| PHLPP1 | enzyme | 0.011 | 2 | IL15,PRKCB |
| SENP3 | peptidase | 0.0442 | 2 | IL15,PML |
| HES3 | transcription regulator | 0.0391 | 2 | IFNAR2,SST |
| BAK1 | other | 0.0226 | 2 | IFNAR2,IL15 |
| Peg13 | other | 0.0176 | 2 | HTR1B,TPH2 |
| Sch-23390 | chemical drug | 0.012 | 2 | HSPA9,NTS |
| APOC3 | transporter | 0.00389 | 2 | HSPA9,HTRA2 |
| TERC | other | 0.0267 | 2 | H1-0,THRA |
| PXR ligand-PXR-Retinoic acid-RXRα | complex | 0.0391 | 2 | GSTM1,PAPSS2 |
| mercuric chloride | chemical toxicant | 0.0282 | 2 | GSTM1,HSPA9 |
| UM101 | chemical drug | 0.0176 | 2 | GPR88,VDR |
| ZC3H12C | other | 0.0226 | 2 | GPR34,PLA2G7 |
| acetyl-L-carnitine | chemical - endogenous mammalian | 0.00829 | 2 | GLO1,PPARGC1A |
| CUL4B | other | 0.0442 | 2 | GLO1,HSPA9 |
| ketamine | chemical drug | 0.0141 | 2 | GABRA5,TH |
| pubchem compound 135889696 | chemical reagent | 0.000774 | 2 | FOXG1,MAP2 |
| ethyl 2-{[(6,7-dimethyl-3-oxo-1,2,3,4-tetrahydro-2-quinoxalinyl)acetyl]amino}-4,5-dimethyl-3-thiophenecarboxylate | chemical reagent | 0.00452 | 2 | FOXG1,MAP2 |
| MSX2 | transcription regulator | 0.0375 | 2 | EPHA4,SP7 |
| PCGEM1 | other | 0.0375 | 2 | ENO1,SUCLG1 |
| GAPDH | enzyme | 0.0327 | 2 | ENO1,SLC38A2 |
| SFMBT1 | transcription regulator | 0.012 | 2 | ENO1,MAP2 |
| dichlorovinylcysteine | chemical toxicant | 0.0188 | 2 | ENO1,HSPA9 |
| HOXA13 | transcription regulator | 0.0131 | 2 | ENO1,EPHA4 |
| PLC | group | 0.00745 | 2 | EGR3,TH |
| polyunsaturated fatty acids | chemical - endogenous non-mammalian | 0.00829 | 2 | DRD2,TH |
| GHRH | other | 0.0141 | 2 | DRD2,SST |
| NGFR | transmembrane receptor | 0.0327 | 2 | DRD2,SNAP25 |
| CP-55940 | chemical reagent | 0.0312 | 2 | DRD2,PRKCB |
| fluphenazine | chemical drug | 0.00108 | 2 | DRD2,NTS |
| remoxipride | chemical drug | 5.26E-05 | 2 | DRD2,NTS |
| DRD2 | G-protein coupled receptor | 0.0425 | 2 | DRD1,KCNJ4 |
| SKF-38393 | chemical reagent | 0.00745 | 2 | DRD1,HSPA9 |
| Klf16 | transcription regulator | 5.26E-05 | 2 | DRD1,DRD2 |
| flupenthixol | chemical drug | 0.00108 | 2 | DRD1,DRD2 |
| manganese | chemical - endogenous mammalian | 0.00666 | 2 | DRD1,DRD2 |
| GSX2 | transcription regulator | 0.00389 | 2 | DLX1,DLX2 |
| PPP4R3B | other | 0.00745 | 2 | DLX1,DLX2 |
| OLIG1 | transcription regulator | 0.000774 | 2 | DLX1,DLX2 |
| RYK | kinase | 0.000774 | 2 | DLX1,DLX2 |
| SLC6A4 | transporter | 0.0342 | 2 | CYP4F8,HTR1B |
| 25-hydroxyvitamin D | chemical drug | 5.26E-05 | 2 | CYP24A1,VDR |
| HR | transcription regulator | 0.0442 | 2 | CYP24A1,VDR |
| calcifediol | chemical - endogenous mammalian | 0.0164 | 2 | CYP24A1,VDR |
| PHEX | peptidase | 0.00389 | 2 | CYP24A1,SP7 |
| histone deacetylase | complex | 0.0342 | 2 | CYP24A1,PPARGC1A |
| TSPYL5 | other | 0.012 | 2 | CNTNAP2,DTD1 |
| Foxp1 | transcription regulator | 0.0327 | 2 | CNR1,PPARGC1A |
| ALDH1A1 | enzyme | 0.00666 | 2 | CNR1,PPARD |
| reserpine | chemical drug | 0.0059 | 2 | CNR1,NTS |
| lysophosphatidylinositol | chemical - endogenous mammalian | 0.0477 | 2 | CNR1,IL15 |
| Hif | complex | 0.0059 | 2 | CKB,TH |
| CL 316243 | chemical drug | 0.0358 | 2 | CKB,PPARGC1A |
| grape seed extract | chemical drug | 0.0226 | 2 | CKB,ENO1 |
| thioridazine | chemical drug | 0.00917 | 2 | CHRM3,NTS |
| CHRM1 | G-protein coupled receptor | 0.00389 | 2 | CHRM1,EGR3 |
| picrotoxin | chemical toxicant | 0.0164 | 2 | CHRM1,CHRM3 |
| icatibant | biologic drug | 0.00389 | 2 | CHRM1,CHRM3 |
| P2RY14 | G-protein coupled receptor | 0.0176 | 2 | CASR,SST |
| GABBR1 | G-protein coupled receptor | 0.0059 | 2 | CASR,SP7 |
| CASZ1 | enzyme | 0.0312 | 2 | CASQ1,HTR1B |
| mir-9 | microRNA | 0.0164 | 2 | CAMTA1,FOXG1 |
| IND S7 | chemical - kinase inhibitor | 0.0176 | 2 | CALU,ENO1 |
| MEL S3 | chemical - kinase inhibitor | 0.0188 | 2 | CALU,ENO1 |
| SETDB1 | enzyme | 0.0425 | 2 | CADM2,IL15 |
| DHX9 | enzyme | 0.00917 | 2 | APOE,SST |
| APOB | transporter | 0.0267 | 2 | APOE,SP7 |
| sucrose | chemical - endogenous mammalian | 0.0327 | 2 | APOE,SNAP25 |
| XRCC6 | enzyme | 0.00745 | 2 | APOE,SLC38A2 |
| LPIN1 | phosphatase | 0.02 | 2 | APOE,PPARGC1A |
| LCAT | enzyme | 0.0131 | 2 | APOE,PPARGC1A |
| COL5A1 | other | 0.0282 | 2 | APOE,LGALS1 |
| MBD1 | transcription regulator | 0.012 | 2 | APOE,EFNA5 |
| NR2C2 | ligand-dependent nuclear receptor | 0.0358 | 2 | APOE,CYP24A1 |
| carbamylcholine | chemical drug | 0.0375 | 2 | APOE,CHRM3 |
| linsidomine | chemical drug | 0.0226 | 2 | ANXA2,PPARD |
| S100A10 | other | 0.00183 | 2 | ANXA2,HTR4 |
| CACNA1C | ion channel | 0.00143 | 2 | ANXA2,ENO1 |
| 2-mercaptoacetate | chemical drug | 0.011 | 2 | ANXA2,ENO1 |
| amlodipine | chemical drug | 0.00666 | 2 | ANXA2,ENO1 |
| UBA1 | enzyme | 0.0131 | 2 | ANXA2,APOE |
| IL23 | complex | 0.0495 | 2 | AHR,RORA |
| BATF | transcription regulator | 0.0188 | 2 | AHR,RORA |
| vitamin A | chemical - endogenous mammalian | 0.024 | 2 | AHR,KCNG1 |
| oleanolic acid | chemical - endogenous non-mammalian | 0.0282 | 2 | AHR,GLO1 |
| TOX | transcription regulator | 0.0131 | 2 | AHR,EGR3 |
| PRKDC | kinase | 0.0188 | 2 | AHR,APOE |
| omeprazole | chemical drug | 0.0477 | 2 | ADORA2A,SST |
| 8-(3-chlorostyryl)caffeine | chemical reagent | 0.000313 | 2 | ADORA2A,NTS |
| MYT1 | transcription regulator | 0.00829 | 2 | ACP1,TH |
| oltipraz | chemical drug | 0.0213 | 2 | ACE,UGT1A6 |
| OLR1 | transmembrane receptor | 0.0477 | 2 | ACE,RXRG |
| Pki | group | 0.00183 | 2 | ACE,NTS |
| MDK | growth factor | 0.0282 | 2 | ACE,LGALS1 |
| 24-hydroxycholesterol | chemical - endogenous mammalian | 0.00519 | 2 | ACE,APOE |
| 1,2-dioctanoyl-sn-glycerol | chemical reagent | 0.0358 | 1 | VDR |
| CYP2R1 | enzyme | 0.00727 | 1 | VDR |
| ITIH4 | other | 0.00727 | 1 | VDR |
| liarozole | chemical drug | 0.0288 | 1 | VDR |
| lactose | chemical - endogenous mammalian | 0.0358 | 1 | VDR |
| SLC6A2 | transporter | 0.0498 | 1 | TPH2 |
| TMEM259 | other | 0.0145 | 1 | TMUB2 |
| RNF185 | enzyme | 0.0145 | 1 | TMUB2 |
| fentanyl | chemical drug | 0.0358 | 1 | TLN2 |
| DRD3 | G-protein coupled receptor | 0.0358 | 1 | TH |
| BBS1 | other | 0.0217 | 1 | TH |
| TAAR1 | G-protein coupled receptor | 0.0145 | 1 | TH |
| SPR | enzyme | 0.0145 | 1 | TH |
| PITX3 | transcription regulator | 0.0498 | 1 | TH |
| SNCB | other | 0.0498 | 1 | TH |
| PHOX2A | transcription regulator | 0.0429 | 1 | TH |
| PTPRN | phosphatase | 0.0358 | 1 | TH |
| PHOX2B | transcription regulator | 0.0429 | 1 | TH |
| PTPRN2 | phosphatase | 0.0429 | 1 | TH |
| GTS 21 | chemical drug | 0.0145 | 1 | TH |
| URB597 | chemical drug | 0.0288 | 1 | TH |
| U-69593 | chemical reagent | 0.00727 | 1 | TH |
| idazoxan | chemical drug | 0.0358 | 1 | TH |
| nortriptyline | chemical drug | 0.0145 | 1 | TH |
| bupropion | chemical drug | 0.0358 | 1 | TH |
| BRF110 | chemical reagent | 0.0217 | 1 | TH |
| myristoylated protein kinase C peptide inhibitor | biologic drug | 0.0288 | 1 | TH |
| iprindole | chemical drug | 0.00727 | 1 | TH |
| mazindol | chemical drug | 0.0145 | 1 | TH |
| methyllycaconitine | chemical toxicant | 0.0429 | 1 | TH |
| dextromethorphan | chemical drug | 0.0288 | 1 | TH |
| FEZF2 | transcription regulator | 0.0358 | 1 | TBR1 |
| NRXN2 | transporter | 0.00727 | 1 | TAFA1 |
| Nrxn3 | other | 0.00727 | 1 | TAFA1 |
| dibutyryl cGMP | chemical reagent | 0.0358 | 1 | SST |
| 1-O-hexadecyl-2-N-methylcarbamol-sn-glycerol-3-phosphocholine | chemical - endogenous mammalian | 0.0288 | 1 | SST |
| Neurotrophin | group | 0.0429 | 1 | SST |
| ALX3 | transcription regulator | 0.0288 | 1 | SST |
| FK-962 | chemical drug | 0.00727 | 1 | SST |
| CHAT | enzyme | 0.0217 | 1 | SST |
| sumatriptan | chemical drug | 0.0429 | 1 | SST |
| gastric acid | chemical - endogenous mammalian | 0.00727 | 1 | SST |
| androsta-1,4,6-triene-3,17-dione | chemical reagent | 0.0429 | 1 | SST |
| advanced glycation end product 3 | chemical - endogenous mammalian | 0.0358 | 1 | SP7 |
| DIPQUO | chemical reagent | 0.0358 | 1 | SP7 |
| AMELY | growth factor | 0.0429 | 1 | SP7 |
| RIOX1 | enzyme | 0.0498 | 1 | SP7 |
| EZH1 | enzyme | 0.0498 | 1 | SP7 |
| miR-637 (and other miRNAs w/seed CUGGGGG) | mature microRNA | 0.0145 | 1 | SP7 |
| OGN | growth factor | 0.0498 | 1 | SP7 |
| HTR4 | G-protein coupled receptor | 0.0429 | 1 | SP7 |
| EDIL3 | other | 0.0358 | 1 | SP7 |
| DSPP | other | 0.0429 | 1 | SP7 |
| HIVEP2 | transcription regulator | 0.0429 | 1 | SP7 |
| FZD2 | G-protein coupled receptor | 0.0498 | 1 | SP7 |
| FZD6 | G-protein coupled receptor | 0.0429 | 1 | SP7 |
| phenylamil | chemical reagent | 0.0429 | 1 | SP7 |
| chitosan | chemical - endogenous mammalian | 0.0217 | 1 | SP7 |
| CLASP2 | other | 0.0288 | 1 | SNAP25 |
| ZFP36L2 | transcription regulator | 0.0498 | 1 | SNAP25 |
| forchlorfenuron | chemical reagent | 0.00727 | 1 | SNAP25 |
| DNAJC5 | other | 0.00727 | 1 | SNAP25 |
| APPL1 | other | 0.0288 | 1 | SNAP25 |
| RAB3B | enzyme | 0.0217 | 1 | SNAP25 |
| VPS35 | transporter | 0.0145 | 1 | SLC38A2 |
| PPP2R2A | phosphatase | 0.0429 | 1 | SLC22A4 |
| PPP2R5B | phosphatase | 0.0358 | 1 | SLC22A4 |
| HSD17B10 | enzyme | 0.00727 | 1 | SH3GL2 |
| phytanic acid | chemical - endogenous mammalian | 0.0498 | 1 | RXRG |
| DNAJA2 | enzyme | 0.0217 | 1 | RORA |
| FGD5-AS1 | other | 0.0288 | 1 | RORA |
| DNAJB1 | transcription regulator | 0.0145 | 1 | RORA |
| bavachalcone | chemical reagent | 0.0145 | 1 | RORA |
| BMS-214662 | chemical drug | 0.0145 | 1 | PRKCB |
| nardostachys chinensis extract | chemical reagent | 0.00727 | 1 | PRKCB |
| 3,4-dideoxyglucosone-3-ene | chemical - endogenous mammalian | 0.00727 | 1 | PRKCB |
| enzastaurin | chemical drug | 0.0498 | 1 | PRKCB |
| epalrestat | chemical drug | 0.0217 | 1 | PRKCB |
| cicletanine | chemical drug | 0.00727 | 1 | PRKCB |
| ingenol mebutate | chemical drug | 0.0498 | 1 | PRKCB |
| 4-CMTB | chemical reagent | 0.0217 | 1 | PPARGC1A |
| Katp Channel (family) | group | 0.00727 | 1 | PPARGC1A |
| Nfatc | group | 0.0358 | 1 | PPARGC1A |
| GP7 | chemical reagent | 0.0429 | 1 | PPARGC1A |
| KCP | other | 0.0145 | 1 | PPARGC1A |
| (S)-3-hydroxy-2-methylpropanoic acid | chemical - endogenous mammalian | 0.0145 | 1 | PPARGC1A |
| AMG-9810 | chemical reagent | 0.00727 | 1 | PPARGC1A |
| Mitochondrial complex 1 | complex | 0.0145 | 1 | PPARGC1A |
| Tug1 | other | 0.00727 | 1 | PPARGC1A |
| KLF14 | transcription regulator | 0.0217 | 1 | PPARGC1A |
| YTHDF2 | other | 0.0429 | 1 | PPARGC1A |
| SLC25A33 | transporter | 0.0145 | 1 | PPARGC1A |
| FNIP1 | other | 0.0498 | 1 | PPARGC1A |
| ACOT13 | enzyme | 0.0429 | 1 | PPARGC1A |
| TXLNG | other | 0.0358 | 1 | PPARGC1A |
| ACOT11 | enzyme | 0.0288 | 1 | PPARGC1A |
| alisporivir | biologic drug | 0.0145 | 1 | PPARGC1A |
| OSTN | other | 0.00727 | 1 | PPARGC1A |
| HAO1 | enzyme | 0.0358 | 1 | PPARGC1A |
| MYBBP1A | transcription regulator | 0.0358 | 1 | PPARGC1A |
| GW 6471 | chemical reagent | 0.0358 | 1 | PPARGC1A |
| SLN | other | 0.0217 | 1 | PPARGC1A |
| ABCC9 | ion channel | 0.0288 | 1 | PPARGC1A |
| cyclic des-acyl ghrelin (6-13) | biologic drug | 0.0145 | 1 | PPARGC1A |
| Cyp2a12/Cyp2a22 | enzyme | 0.0288 | 1 | PPARGC1A |
| oligomycin A | chemical - endogenous non-mammalian | 0.0145 | 1 | PPARGC1A |
| pterosin B | chemical reagent | 0.0288 | 1 | PPARGC1A |
| CGP 12177 | chemical reagent | 0.0217 | 1 | PPARGC1A |
| arginine methyltransferase inhibitor-1 | chemical reagent | 0.0429 | 1 | PPARGC1A |
| BRL 37344 | chemical reagent | 0.0429 | 1 | PPARGC1A |
| trimetazidine | chemical drug | 0.0288 | 1 | PPARGC1A |
| cAMP-dependent protein kinase | complex | 0.0145 | 1 | PPARGC1A |
| K ATP Channel | complex | 0.0217 | 1 | PPARGC1A |
| FGFR3-TACC3 | fusion gene/product | 0.00727 | 1 | PPARGC1A |
| CRTC1-MAML2 | fusion gene/product | 0.0217 | 1 | PPARGC1A |
| adiporon | chemical reagent | 0.0498 | 1 | PPARGC1A |
| GSK3235025 | chemical reagent | 0.0288 | 1 | PPARGC1A |
| 10,12-tricosadiynoic acid | chemical reagent | 0.0429 | 1 | PPARGC1A |
| MHY1485 | chemical reagent | 0.0145 | 1 | PPARGC1A |
| alpha-keto-beta-methylvaleric acid | chemical - endogenous mammalian | 0.0498 | 1 | PPARGC1A |
| pyrrolidonecarboxylic acid | chemical - endogenous mammalian | 0.0498 | 1 | PPARGC1A |
| RSAD2 | enzyme | 0.0429 | 1 | PPARD |
| afamelanotide | biologic drug | 0.0288 | 1 | PPARD |
| d18:1/48:2 omega-O-linoleoyl-ceramide | chemical - endogenous mammalian | 0.0217 | 1 | PPARD |
| IVNS1ABP | other | 0.0217 | 1 | PML |
| ZRL5P4 | chemical reagent | 0.0145 | 1 | PML |
| platelet activating factor-C16 | chemical - endogenous mammalian | 0.0288 | 1 | PLA2G7 |
| resolvin D5 | chemical - endogenous mammalian | 0.0429 | 1 | PLA2G7 |
| SLCO1C1 | transporter | 0.0288 | 1 | Nrgn |
| NUMB/NUMBL | group | 0.00727 | 1 | MSTN |
| ostarine | chemical drug | 0.0217 | 1 | MSTN |
| SENP2 | peptidase | 0.0498 | 1 | MSTN |
| olomoucine | chemical - kinase inhibitor | 0.0498 | 1 | MAP2 |
| pregnanolone | chemical - endogenous mammalian | 0.00727 | 1 | MAP2 |
| coconut oil | chemical drug | 0.0498 | 1 | MAP2 |
| ZNF536 | transcription regulator | 0.0358 | 1 | MAP2 |
| tri-o-cresyl phosphate | chemical reagent | 0.0288 | 1 | MAP2 |
| 1-methyl-D-tryptophan | chemical drug | 0.00727 | 1 | MAP2 |
| GALNS | enzyme | 0.0288 | 1 | MAP2 |
| Dst | other | 0.0145 | 1 | MAP2 |
| opaganib | chemical drug | 0.0217 | 1 | MAP2 |
| MME | peptidase | 0.0288 | 1 | MAP2 |
| betel quid extract | chemical reagent | 0.00727 | 1 | MAP2 |
| SKF-83959 | chemical reagent | 0.0217 | 1 | MAP2 |
| 3beta-methoxypregnenolone | chemical drug | 0.00727 | 1 | MAP2 |
| CYM-5520 | chemical reagent | 0.0288 | 1 | MAP2 |
| ergothioneine | chemical - endogenous non-mammalian | 0.00727 | 1 | MAP2 |
| E-64c | chemical - protease inhibitor | 0.00727 | 1 | MAP2 |
| TH17 Cytokine | group | 0.0217 | 1 | LGALS1 |
| PCAT6 | other | 0.00727 | 1 | KLHL12 |
| HTR6 | G-protein coupled receptor | 0.0217 | 1 | KIF3A |
| SSTR3 | G-protein coupled receptor | 0.0288 | 1 | KIF3A |
| CpG ODN 1555 | chemical reagent | 0.0358 | 1 | IL15 |
| bis(3',5')-cyclic diguanylic acid | chemical - endogenous non-mammalian | 0.0498 | 1 | IL15 |
| PF-4418948 | chemical drug | 0.0498 | 1 | IL15 |
| KLF12 | transcription regulator | 0.0498 | 1 | IL15 |
| atacicept | biologic drug | 0.00727 | 1 | IL15 |
| TUSC2 | other | 0.0217 | 1 | IL15 |
| di-n-propyl disulfide | chemical reagent | 0.0288 | 1 | GSTM1 |
| SR-3420 | chemical reagent | 0.00727 | 1 | GPR3 |
| CAY10499 | chemical reagent | 0.0145 | 1 | GPR3 |
| atglistatin | chemical reagent | 0.00727 | 1 | GPR3 |
| ACADS | enzyme | 0.0217 | 1 | GLO1 |
| nandrolone decanoate | chemical drug | 0.0288 | 1 | GABRA5 |
| testosterone cypionate | chemical drug | 0.0288 | 1 | GABRA5 |
| methyltestosterone | chemical drug | 0.0358 | 1 | GABRA5 |
| ELAVL2 | other | 0.0217 | 1 | FOXG1 |
| CHRD | other | 0.0429 | 1 | FOXG1 |
| RIPPLY2 | other | 0.0217 | 1 | EPHA4 |
| POU3F4 | transcription regulator | 0.0217 | 1 | EPHA4 |
| INSL5 | other | 0.0358 | 1 | ENO1 |
| PCCA-DT | other | 0.0288 | 1 | ENO1 |
| ethyl protocatechuate | chemical reagent | 0.0358 | 1 | ENO1 |
| 1,4,5-IP3 | chemical - endogenous mammalian | 0.0217 | 1 | EGR3 |
| edratide | biologic drug | 0.0358 | 1 | EGR3 |
| RAB4 | group | 0.00727 | 1 | DRD2 |
| CC2D1A | transcription regulator | 0.0498 | 1 | DRD2 |
| Mir124a-1hg | other | 0.0217 | 1 | DRD2 |
| tiapride | chemical drug | 0.00727 | 1 | DRD2 |
| GTF2F2 | transcription regulator | 0.0217 | 1 | DRD2 |
| Bc1-ps1 | translation regulator | 0.00727 | 1 | DRD2 |
| COMT | enzyme | 0.0498 | 1 | DRD2 |
| GRK6 | kinase | 0.0429 | 1 | DRD2 |
| DRD4 | G-protein coupled receptor | 0.0498 | 1 | DRD2 |
| Smptb | other | 0.0145 | 1 | DRD2 |
| amisulpride | chemical drug | 0.00727 | 1 | DRD2 |
| heroin | chemical drug | 0.0429 | 1 | DRD2 |
| droperidol | chemical drug | 0.00727 | 1 | DRD2 |
| S-alpha-methyl-4-carboxyphenylglycine | chemical reagent | 0.00727 | 1 | DRD1 |
| SLC1A1 | transporter | 0.0429 | 1 | DRD1 |
| PDCD6IP | other | 0.0358 | 1 | DRD1 |
| NEFM | other | 0.00727 | 1 | DRD1 |
| 6-methyl-2-(phenylethynyl)pyridine | chemical reagent | 0.0498 | 1 | DRD1 |
| (+)-butaclamol | chemical reagent | 0.0145 | 1 | DRD1 |
| FOXQ1 | transcription regulator | 0.0429 | 1 | DACH1 |
| 1-alpha,24(R),25-trihydroxyvitamin D3 | chemical - endogenous mammalian | 0.0145 | 1 | CYP24A1 |
| 24R,25-dihydroxyvitamin D3 | chemical - endogenous mammalian | 0.0145 | 1 | CYP24A1 |
| MN1 | other | 0.0429 | 1 | CYP24A1 |
| GC | transporter | 0.0217 | 1 | CYP24A1 |
| phosphorus | chemical reagent | 0.0288 | 1 | CYP24A1 |
| DP-001 | chemical drug | 0.0288 | 1 | CYP24A1 |
| 1beta,25-dihydroxyvitamin D3 | chemical drug | 0.0145 | 1 | CYP24A1 |
| 2-hydroxy-imino phenylpyruvic acid | chemical reagent | 0.0145 | 1 | CTBP2 |
| 2-keto-4-methylthiobutyric acid | chemical - endogenous mammalian | 0.0145 | 1 | CTBP2 |
| Becn2 | other | 0.00727 | 1 | CNR1 |
| lorglumide | chemical reagent | 0.0288 | 1 | CNR1 |
| GPRASP1 | transporter | 0.00727 | 1 | CNR1 |
| cannabinoid | chemical drug | 0.0358 | 1 | CNR1 |
| ASB9 | transcription regulator | 0.00727 | 1 | CKB |
| mir-483 | microRNA | 0.0358 | 1 | CKB |
| 4-diphenylacetoxy-1,1-dimethylpiperidinium | chemical reagent | 0.00727 | 1 | CHRM3 |
| thioperamide | chemical reagent | 0.00727 | 1 | CHRM3 |
| Astra 1397 | chemical reagent | 0.00727 | 1 | CHRM3 |
| quinuclidinyl benzilate | chemical reagent | 0.00727 | 1 | CHRM3 |
| atropine | chemical drug | 0.0498 | 1 | CHRM3 |
| N-methylscopolamine | chemical drug | 0.0145 | 1 | CHRM3 |
| pirenzepine | chemical drug | 0.0217 | 1 | CHRM1 |
| GABBR2 | G-protein coupled receptor | 0.0498 | 1 | CASR |
| RNF19A | enzyme | 0.00727 | 1 | CASR |
| cinacalcet | chemical drug | 0.0145 | 1 | CASR |
| TRDN | other | 0.0288 | 1 | CASQ1 |
| miR-9-3p (and other miRNAs w/seed UAAAGCU) | mature microRNA | 0.0429 | 1 | CAMTA1 |
| LGH447 | chemical drug | 0.0429 | 1 | ATP6V1A |
| MLXIP | transcription regulator | 0.0145 | 1 | ARRDC4 |
| 4-aminopyrazolo(3,4-d)pyrimidine | chemical reagent | 0.0145 | 1 | APOE |
| hyodeoxycholic acid | chemical - endogenous mammalian | 0.0217 | 1 | APOE |
| 24(S),25-epoxycholesterol | chemical - endogenous mammalian | 0.0498 | 1 | APOE |
| triamcinolone | chemical drug | 0.0498 | 1 | APOE |
| DNAJA4 | other | 0.00727 | 1 | APOE |
| AFM | transporter | 0.0145 | 1 | APOE |
| Pzp | other | 0.0429 | 1 | APOE |
| lynestrenol | chemical drug | 0.0145 | 1 | APOE |
| 2-[[4-[(e)-styryl]phenoxy]methyl]oxirane | chemical reagent | 0.0288 | 1 | APOE |
| ALYREF | transcription regulator | 0.0145 | 1 | APOE |
| non-esterified fatty acid | chemical - endogenous mammalian | 0.0498 | 1 | APOE |
| gemcabene | chemical drug | 0.0288 | 1 | APOE |
| 20alpha-hydroxycholesterol | chemical - endogenous mammalian | 0.0498 | 1 | APOE |
| hydrocortisone phosphate | chemical drug | 0.00727 | 1 | APOE |
| 22-hydroxycholesterol | chemical - endogenous mammalian | 0.0145 | 1 | APOE |
| YES1 | kinase | 0.0217 | 1 | ANXA2 |
| PFN1 | other | 0.0358 | 1 | ANXA2 |
| TDO2 | enzyme | 0.0498 | 1 | AHR |
| 2,2-(2-chlorophenyl-4'-chlorophenyl)-1,1-dichloroethene | chemical - endogenous mammalian | 0.0145 | 1 | AHR |
| PD 150606 | chemical - protease inhibitor | 0.0498 | 1 | AHR |
| 8-phenyltheophylline | chemical reagent | 0.00727 | 1 | ADORA2A |
| darusentan | chemical drug | 0.0358 | 1 | ACE |
| PITHD1 | other | 0.0498 | 1 | ACE |
| temocapril | chemical reagent | 0.0217 | 1 | ACE |
| olmesartan medoxomil | chemical drug | 0.0429 | 1 | ACE |
| quinapril | chemical drug | 0.0429 | 1 | ACE |
| eugenol | chemical - endogenous non-mammalian | 0.0498 | 1 | ACE |
| trigonelline | chemical - endogenous mammalian | 0.0429 | 1 | ACE |
| D,L-propargylglycine | chemical reagent | 0.0498 | 1 | ACE |
| fosinopril | chemical drug | 0.0217 | 1 | ACE |
