## Supplemental Table 4: All upstream regulators of muscle molecules for "“It All Rolls Downstream: Upstream Control of Physical Activity Regulation”"

**Supplemental Table 4: Upstream Regulators of Muscle Target Molecules**

| **Upstream Regulator** | **Molecule Type** | **p-value of overlap** | **# of**  **molecules** | **Target Molecules in Dataset** |
| --- | --- | --- | --- | --- |
| beta-estradiol | chemical - endogenous mammalian | 1.79E-09 | 41 | AARS1,ACADVL,ACE,AFF4,AHR,AKAP10,ANXA4,APBA1,APOA1,APOE, ATP5F1B, CA14, CHRM3, CNR1, CUBN, BYP24A1, DNAJC1, ENEO1, GABRG3, IFI30, ILF3, IQGAP2, LUM, MYH10, OGDH, PAPSS2, PMM2, PPARGC1A, PRKCB, RDX, SERPINA1, SERPINA3, SLCA4, SNAP25, SPARC, TF, TP53, TPH2, TXRD1, VCL, VIM |
| TP53 | transcription regulator | 1E-08 | 34 | ACADVL,ACE,ALDH1A1,ANXA4,ANXA6,APOA1,APOE,CALU,CYP24A1,DNAJB4, FKBP4, IFI30, MYH10, MYL3, NDUFS3, OGDH, PCK1, PDHB, PDHX, PDP1, PFKFB1, PML, PPARD, PPARGC1A, PRDX6, PRKCB, SDHA, SERPINA3, SLCA4, SP7, TP53, TUBB3, VCL, VIM |
| lipopolysaccharide | chemical drug | 1.45E-05 | 32 | AARS1,ACE,AHR,ALDH1A1,ANXA5,APOA1,APOE,CASR,CCT6A,CUBN, ENO1,EPHX2, IFI30, IFNAR2, IL15RA, MT1, MTMR14, MYL3, PAPSS2, PML, PPARD, PPARGC1A, PRKCB, SDHA, SERPINA1, SERPINA3, SPARC, TF, TP53, VCL, VIM, YWHAQ |
| tretinoin | chemical - endogenous mammalian | 3.16E-06 | 27 | AHR,ALDH1A1,ANXA5,ANXA6,APOE,CNR1,CNTNAP2,CUBN,HPX,ILF3, MAP2, MT1, PCK1,PML, PPARD, PPARGC1A, PRKCB, RAI14, SERPINA1, SLC2A4, SP7, SPARC, TF, TP53, TUBB3, VCL, VIM |
| dexamethasone | chemical drug | 0.000388 | 27 | ACE,ALDH1A1,APOA1,APOE,CYP24A1,ENO1,FKBP4,IFNAR2,ILF3,MAP2, MSTN, MT1, PCK1, PMM2, PPARD, PPARGC1A, PRDX6, PRKCB, SDHA, SERPINA1, SERPINA3, SLC2A4, SPARC, TF, TP53, VCL, VIM |
| TNF | cytokine | 0.000118 | 10 | ACADVL,ACE,APOA1,APOE,GABRA5,IFNAR2,IL15RA,MAPKAP1,MSTN,MYH10 |
| HNF4A | transcription regulator | 0.00424 | 10 | ACTR1A,AHSG,ALDH1A1,ANXA5,APOA1,APOE,CFL2,DNAJB4,ETFDH,HPX |
| TGFB1 | growth factor | 0.00173 | 10 | ACE,ADAMTS10,AHR,APOE,CCT6A,DNAJB4,ENO1,IFI30,MSTN,MYL3 |
| IFNG | cytokine | 0.000117 | 10 | ACE,AHR,CNR1,CYP24A1,ELAVL2,ENO1,IFI30,IL15RA,MAP2,MYH10 |
| MYC | transcription regulator | 3.05E-05 | 10 | AHR,ANXA4,ANXA5,ANXA6,CNTNAP2,ENO1,IQGAP2,LUM,Mt1,PCK1 |
| Insulin | group | 7.1E-09 | 10 | APOA1,APOE,CA14,CCT6A,CHRM3,DNAJB4,Mt1,PCK1,PDHB,PPARD |
| D-glucose | chemical - endogenous mammalian | 1.98E-08 | 10 | AARS1,ACE,CNR1,CYP24A1,ENO1,Mt1,OGDH,PCK1,PDHX,PPARD |
| HRAS | enzyme | 1.72E-07 | 10 | ANXA6,ATP5F1B,CALU,CYP24A1,IFI30,IQGAP2,MYH10,MYL3,PCK1,PML |
| HTT | transcription regulator | 2.45E-07 | 10 | ACTN2,APOA1,APOE,ATP5F1B,CNR1,FKBP4,MAP2,Mt1,Mup1 (includes others),NDUFS3 |
| trichostatin A | chemical drug | 0.000024 | 10 | ACOT8,AHR,ALDH1A1,ANXA6,APOE,CNTNAP2,CTBP2,ILF3,MAP2,PMM2 |
| ESR1 | ligand-dependent nuclear receptor | 0.00197 | 10 | ACE,ANXA4,APOA1,APOE,ATP6V1A,CFL2,DNAJC1,ENO1,FKBP4,HPX |
| CTNNB1 | transcription regulator | 3.45E-05 | 10 | ACADVL,AHR,ALDH1A1,CNR1,CYP24A1,ILF3,Mup1 (includes others),PCK1,PML,PPARD |
| calcitriol | chemical drug | 2.76E-07 | 10 | AHR,APOA1,ATAD1,CASR,COQ9,CYP24A1,EFL1,KSR2,MMS22L,PPARD |
| dihydrotestosterone | chemical - endogenous mammalian | 3.58E-07 | 10 | ACADVL,ACE,APOE,CNTNAP2,ENO1,FKBP4,IQGAP2,MSTN,Mt1,Mup1 (includes others) |
| APP | other | 0.000261 | 10 | ACTN2,ANXA5,ANXA6,APOE,ATP5F1B,ATP6V1A,CFL2,ENO1,MAP2,Mt1 |
| IL1B | cytokine | 0.00138 | 10 | ACE,ALDH1A1,ANXA4,APOE,CASR,CYP24A1,IFNAR2,IL15RA,Mt1,PPARGC1A |
| MAPT | other | 2.64E-07 | 10 | AARS1,ACTR1A,ANXA5,ANXA6,ATP5F1B,ATP6V1A,CFL2,ENO1,MAP2,NDUFS3 |
| CREB1 | transcription regulator | 1.7E-06 | 10 | APOE,CALN1,GABRA5,MYH10,PCK1,PPARGC1A,PRCP,SH3KBP1,SLC18A2,SLC2A4 |
| EGF | growth factor | 3.33E-05 | 10 | APOA1,PFKFB1,PPARD,SERPINA1,SERPINA3,SLC2A4,SNAP25,SPARC,TF,TP53 |
| SP1 | transcription regulator | 1.04E-05 | 10 | APOA1,APOE,ATP5F1B,Mt1,PCK1,PDHX,PPARD,PRKCB,SLC22A4,SNAP25 |
| AR | ligand-dependent nuclear receptor | 0.000118 | 10 | ALDH1A1,CALU,FKBP4,IFNAR2,MSTN,MYL3,PCK1,SDHA,SERPINA3,SLC22A4 |
| pirinixic acid | chemical toxicant | 4.94E-07 | 10 | ACADVL,ACOT8,AHSG,APOA1,APOE,CCT6A,ETFDH,HPX,Mup1 (includes others),PDHB |
| LEP | growth factor | 1.67E-06 | 10 | ACADVL,ACE,APOA1,CYP24A1,EPHX2,ETFDH,HPX,PCK1,PPARGC1A,SLC2A4 |
| decitabine | chemical drug | 0.0015 | 10 | APOE,CNTNAP2,CTBP2,CYP24A1,ILF3,LUM,MAP2,SPARC,TP53,TUBB3 |
| RELA | transcription regulator | 5.86E-06 | 10 | AHR,APOE,CASR,IL15RA,LUM,PCK1,PRDX6,SH3KBP1,SLC2A4,SP7 |
| Immunoglobulin | complex | 0.00575 | 10 | AHR,APOE,ATP5F1B,ENO1,IFI30,IL15RA,OGDH,PCK1,PDHB,PPARGC1A |
| IL6 | cytokine | 0.0012 | 10 | AHR,APOA1,APOE,HPX,MAP2,Mt1,SERPINA1,SERPINA3,SLC2A4,SP7 |
| STAT3 | transcription regulator | 0.000468 | 10 | AHR,AHSG,ALDH1A1,IFI30,MAP2,Mt1,PCK1,PML,PPARGC1A,SERPINA1 |
| tetradecanoylphorbol acetate | chemical drug | 0.0277 | 10 | ACE,CYP24A1,ILF3,PCK1,PPARD,PRKCB,SERPINA1,SERPINA3,SNAP25,SPARC |
| AGT | growth factor | 0.00361 | 10 | ACE,CNR1,EFNA5,LUM,MAP2,MYH10,PPARGC1A,Ppp2r5c,SERPINA3,SPARC |
| PD98059 | chemical - kinase inhibitor | 4.63E-05 | 10 | ACE,APOA1,CYP24A1,ENO1,MSTN,PCK1,PFKFB1,PPARD,SERPINA1,SP7 |
| forskolin | chemical toxicant | 0.000524 | 10 | ACE,APOA1,ATP6V1A,Mt1,PCK1,PPARGC1A,SLC18A2,SLC2A4,SNAP25,STK24 |
| SMARCA4 | transcription regulator | 0.000114 | 10 | ACE,AHR,APOA1,CA14,IFI30,IL15RA,LUM,Mt1,PDP1,SLC2A4 |
| INSR | kinase | 2.86E-05 | 10 | ACADVL,ALDH1A1,APOA1,ATP5F1B,ETFDH,HPX,IFNAR2,OGDH,PCK1,PDHB |
| PPARA | ligand-dependent nuclear receptor | 5.12E-06 | 10 | ACADVL,ACOT8,APOA1,APOE,ATP5F1B,HPX,Mup1 (includes others),PCK1,PPARD,PPARGC1A |
| GABA | chemical - endogenous mammalian | 9.25E-06 | 10 | APOE,ATP5F1B,CHRM3,MYH10,PAPSS2,PDHB,PMM2,PRDX6,RDX,TP53 |
| CEBPB | transcription regulator | 0.000304 | 10 | ALDH1A1,CYP24A1,HPX,MMS22L,PCK1,PPARD,PPARGC1A,SERPINA1,TF,TP53 |
| L-triiodothyronine | chemical - endogenous mammalian | 0.000129 | 10 | AHSG,ALDH1A1,APOA1,CNR1,Mup1 (includes others),PCK1,PDHB,PPARGC1A,SLC18A2,SLC2A4 |
| NR3C1 | ligand-dependent nuclear receptor | 0.00198 | 10 | ACTN2,APOE,FKBP4,IL15RA,Mt1,PCK1,PRKCB,SERPINA1,SH3KBP1,TP53 |
| PPARG | ligand-dependent nuclear receptor | 5.88E-05 | 10 | ACOT8,APOA1,APOE,MAP2,PCK1,PDHB,PPARD,PPARGC1A,SERPINA1,SLC2A4 |
| GLI1 | transcription regulator | 8.43E-05 | 10 | ACADVL,ANXA6,CFL2,ELAVL2,IQGAP2,PAPSS2,PPARGC1A,SDHA,SERPINA1,SP7 |
| sirolimus | chemical drug | 0.000367 | 10 | ACADVL,ANXA5,ATP6V1A,ENO1,Mt1,PML,PPARD,PPARGC1A,RPL13A,SLC2A4 |
| LEPR | transmembrane receptor | 2.34E-08 | 10 | ANXA5,APOA1,APOE,CNR1,EXOC4,HPX,Mup1 (includes others),PPARD,PPARGC1A,SLC2A4 |
| methylprednisolone | chemical drug | 0.00154 | 10 | ALDH1A1,CCT6A,EPHX2,FKBP4,Mt1,PCYT2,SERPINA1,SERPINA3,SPARC,TP53 |
| palmitic acid | chemical - endogenous mammalian | 9.94E-07 | 10 | AHR,AHSG,CNR1,EPHX2,Mt1,PCK1,PPARD,PPARGC1A,SLC2A4,SP7 |
| FGF2 | growth factor | 0.000224 | 10 | ACE,CHRM3,ENO1,SERPINA1,SP7,SPARC,TF,TP53,TUBB3,VIM |
| mono-(2-ethylhexyl)phthalate | chemical toxicant | 5.28E-09 | 10 | ACADVL,ATP5F1B,ENO1,OGDH,PCK1,PDHB,PPARD,PPARGC1A,SDHA,SLC2A4 |
| rosiglitazone | chemical drug | 0.000167 | 10 | ACADVL,AHSG,APOE,CYP24A1,EPHX2,ETFDH,Mup1 (includes others),PCK1,PPARGC1A,SDHA |
| levodopa | chemical - endogenous mammalian | 0.00684 | 10 | CNR1,EFNA5,IFNAR2,MAPKAP1,PCYT2,RHBDL3,SP7,SPOCK3,TP53,VAT1 |
| DMD | other | 7.37E-07 | 10 | CAMTA1,CASQ1,CCT6A,CFL2,MSTN,Mup1 (includes others),PPARGC1A,SERPINA1,SLC2A4,VIM |
| PSEN1 | peptidase | 0.000128 | 10 | APOE,ATP5F1B,ATP6V1A,CFL2,ENO1,MAP2,PRDX6,SNAP25,TP53,TUBB3 |
| ADIPOQ | other | 2.91E-06 | 10 | APOA1,Mup1 (includes others),PCK1,PDHB,PFKFB1,PPARD,PPARGC1A,SLC2A4,SP7,TP53 |
| SMAD3 | transcription regulator | 2.82E-05 | 10 | APOA1,LUM,MSTN,PPARD,PPARGC1A,SLC2A4,SP7,SPARC,TF,VIM |
| KRAS | enzyme | 0.0286 | 10 | AHR,CYP24A1,ENO1,IQGAP2,PPARD,RAI14,SERPINA3,SPARC,TP53,VIM |
| glucocorticoid | chemical drug | 1.44E-05 | 10 | AHR,CNR1,CYP24A1,Mt1,Mup1 (includes others),PCK1,PPARGC1A,SERPINA3,TP53,TPH2 |
| NFkB (complex) | complex | 0.00541 | 10 | AHR,APOE,IFNAR2,IL15RA,MSTN,SERPINA3,SLC22A4,SLC2A4,TP53,VIM |
| FOS | transcription regulator | 0.00256 | 10 | AFF4,ANXA4,CALU,ENO1,EPHX2,PCYT2,Ppp2r5c,TAFA1,TP53,VIM |
| metribolone | chemical reagent | 0.000587 | 10 | ACTN2,ATP5F1B,ENO1,ETFDH,IQGAP2,PDHB,PDHX,PMM2,PRDX6,TP53 |
| 8-bromo-cAMP | chemical reagent | 0.00113 | 10 | ACE,APOA1,IQGAP2,MYH10,PCK1,PPARD,PPARGC1A,SLC2A4,TP53,VCL |
| OSM | cytokine | 0.00196 | 10 | ACE,AHR,IL15RA,MAP2,MYH10,OGDH,SERPINA1,SERPINA3,TP53,VIM |
| PPARGC1A | transcription regulator | 0.000362 | 10 | ACADVL,ATP5F1B,ENO1,LUM,PCK1,PPARGC1A,SDHA,SERPINA3,SLC2A4,TP53 |
| cisplatin | chemical drug | 0.0274 | 10 | ACADVL,ALDH1A1,ANXA4,ANXA5,Mt1,PPARGC1A,RPL13A,SPARC,TP53,VIM |
| PTEN | phosphatase | 0.0356 | 9 | IFI30,NDUFS3,PDHB,PPARGC1A,PRKCB,SDHA,SPARC,TP53,VIM |
| MTOR | kinase | 0.000327 | 9 | ENO1,MAP2,PDHX,PPARD,PPARGC1A,SERPINA1,SLC2A4,SND1,TP53 |
| FOXO1 | transcription regulator | 0.0023 | 9 | ATP5F1B,CNR1,MYL3,PCK1,PDHB,PPARGC1A,PRCP,SLC2A4,TP53 |
| PML | transcription regulator | 6.47E-07 | 9 | ANXA4,APOA1,APOE,DNAJB4,PML,PPARGC1A,TP53,TXNRD1,VIM |
| butyric acid | chemical - endogenous mammalian | 0.00189 | 9 | ALDH1A1,ANXA5,IFI30,IFNAR2,PCK1,PPARGC1A,PRKCB,SLC2A4,TP53 |
| SOX2 | transcription regulator | 0.00314 | 9 | AHR,ALDH1A1,CASR,S100A1,SERPINA1,TF,TUBB3,TXNRD1,VIM |
| IGF1 | growth factor | 0.00958 | 9 | ACTN2,MSTN,RDX,SLC2A4,SP7,TF,TP53,TUBB3,VIM |
| AMPK | complex | 4.32E-08 | 9 | ACE,MSTN,PCK1,PDHB,PPARGC1A,SDHA,SLC2A4,TP53,VIM |
| hydrogen peroxide | chemical - endogenous mammalian | 0.00571 | 9 | ACE,ATP6V1A,CNR1,MAPKAP1,PPARGC1A,SPARC,TP53,VIM,YWHAQ |
| progesterone | chemical - endogenous mammalian | 0.0215 | 9 | ACE,ALDH1A1,APOE,CNR1,DNAJB4,FKBP4,PCK1,TP53,VIM |
| HIF1A | transcription regulator | 0.00584 | 9 | ACE,AHR,APOE,ATP5F1B,ENO1,SDHA,SLC2A4,TP53,VIM |
| PPARD | ligand-dependent nuclear receptor | 7.76E-05 | 9 | ACADVL,APOA1,APOE,ENO1,PPARD,PPARGC1A,SLC2A4,SP7,TP53 |
| nitrofurantoin | chemical drug | 6.47E-07 | 9 | ACADVL,ANXA5,APOE,HPX,Mt1,PRKCB,RPL13A,SERPINA1,TP53 |
| testosterone | chemical - endogenous mammalian | 0.0023 | 8 | MSTN,Mt1,PRKCB,RAI14,TF,TP53,TUBB3,TXNRD1 |
| SMARCB1 | transcription regulator | 4.58E-05 | 8 | IFNAR2,IL15RA,Mup1 (includes others),PCK1,PFKFB1,S100A1,SPARC,TP53 |
| SB203580 | chemical drug | 0.00938 | 8 | ENO1,MSTN,PCK1,PPARD,PPARGC1A,SPARC,TP53,VIM |
| U0126 | chemical drug | 0.022 | 8 | CYP24A1,PML,PPARD,PRKCB,SP7,TF,TP53,VIM |
| ADCYAP1 | other | 0.00009 | 8 | CNTNAP2,LUM,PRDX6,SERPINA3,SLC18A2,SNAP25,SPARC,TXNRD1 |
| tazemetostat | chemical drug | 0.000618 | 8 | CNTNAP2,CTBP2,ILF3,MAP2,TP53,TUBB3,VCL,ZMYM2 |
| SP2509 | chemical reagent | 0.000534 | 8 | CNTNAP2,CTBP2,ILF3,MAP2,TP53,TUBB3,VCL,ZMYM2 |
| RORA | ligand-dependent nuclear receptor | 4.73E-05 | 8 | APOE,Mt1,Mup1 (includes others),PCK1,PPARGC1A,PRDX6,SLC2A4,VIM |
| curcumin | chemical drug | 0.00212 | 8 | APOE,MAP2,OGDH,PPARGC1A,SLC2A4,TP53,TUBB3,VIM |
| TO-901317 | chemical reagent | 9.27E-05 | 8 | APOE,ENO1,Mt1,Mup1 (includes others),PCK1,RAI14,SERPINA3,SLC2A4 |
| SOD1 | enzyme | 0.000286 | 8 | APOE,CNR1,GABRG3,Mt1,PRDX6,RDX,SP7,TP53 |
| LY294002 | chemical drug | 0.0161 | 8 | APOE,CNR1,CUBN,MAP2,PPARD,SLC2A4,TP53,VIM |
| NR4A1 | ligand-dependent nuclear receptor | 0.000368 | 8 | APOE,ATP5F1B,ENO1,Mt1,OGDH,PCK1,PDHB,SLC2A4 |
| carbon tetrachloride | chemical toxicant | 0.00012 | 8 | APOA1,APOE,CNR1,Mt1,PPARD,TF,TP53,VIM |
| methapyrilene | chemical drug | 1.77E-06 | 8 | ANXA5,APOE,Mt1,PDP1,PRKCB,RPL13A,TP53,TXNRD1 |
| camptothecin | chemical drug | 0.0361 | 8 | ANXA4,APOE,IL15RA,IQGAP2,Mt1,RDX,SLC2A4,TP53 |
| AHR | ligand-dependent nuclear receptor | 0.01 | 8 | AHSG,ENO1,EPHX2,Mt1,PCK1,TP53,VCL,VIM |
| FSH | complex | 0.00302 | 8 | AHR,PCK1,SLC2A4,SNAP25,STK24,TF,TP53,VCL |
| POU5F1 | transcription regulator | 0.00268 | 8 | AHR,ENO1,PDHX,SERPINA1,TF,TP53,TXNRD1,VIM |
| acetaminophen | chemical drug | 0.00009 | 8 | ACOT8,AHR,ALDH1A1,Mt1,PRDX6,PRKCB,TF,TXNRD1 |
| VDR | transcription regulator | 0.000193 | 8 | ACE,CASR,CUBN,CYP24A1,KSR2,SERPINA1,TPH2,VCL |
| STAT1 | transcription regulator | 0.00376 | 8 | ACE,APOA1,APOE,IFI30,IL15RA,PPARD,SERPINA3,TP53 |
| NFE2L2 | transcription regulator | 0.00806 | 8 | ACE,AHR,Mt1,PPARGC1A,PRKCB,SERPINA3,TP53,TXNRD1 |
| CPT1B | enzyme | 4.15E-05 | 8 | ACADVL,ATP5F1B,NDUFS3,OGDH,PCK1,PPARGC1A,SDHA,SLC2A4 |
| CLPP | peptidase | 1.45E-08 | 8 | ACADVL,ATP5F1B,ENO1,ETFDH,OGDH,PDHB,PPARGC1A,SDHA |
| WNT3A | cytokine | 0.00126 | 8 | ACADVL,AHR,CYP24A1,SDHA,SP7,SPARC,TUBB3,VIM |
| SREBF1 | transcription regulator | 0.000266 | 8 | AARS1,IFI30,PCK1,SERPINA1,SERPINA3,SLC22A4,TF,TP53 |
| CD3 | complex | 0.0361 | 8 | AARS1,AHR,APBA1,HPX,IL15RA,OGDH,PCYT2,TXNRD1 |
| Akt | group | 0.0108 | 7 | MAP2,PCK1,PPARD,PPARGC1A,SLC2A4,TP53,VIM |
| AKT1 | kinase | 0.00215 | 7 | ENO1,MSTN,MYH10,PCK1,PPARGC1A,TP53,VIM |
| streptozocin | chemical drug | 0.00189 | 7 | CASR,PCK1,PPARD,PPARGC1A,SLC2A4,TP53,VIM |
| NR3C2 | ligand-dependent nuclear receptor | 2.75E-05 | 7 | APOE,FKBP4,MAP2,PDHB,SERPINA3,TP53,TUBB3 |
| cyclosporin A | biologic drug | 0.0182 | 7 | APOE,CASR,HPX,IQGAP2,MYH10,TP53,VIM |
| TCR | complex | 0.00749 | 7 | APOE,ATP5F1B,ENO1,PRDX6,RPL13A,SDHA,SERPINA3 |
| FOXA2 | transcription regulator | 0.000646 | 7 | APOA1,Mt1,PCK1,PDHX,SERPINA1,SLC2A4,TF |
| ethanol | chemical - endogenous mammalian | 0.0186 | 7 | APOA1,CNR1,MAP2,PCK1,PPARGC1A,SERPINA1,TP53 |
| APOE | transporter | 0.00606 | 7 | APOA1,APOE,FKBP4,MAP2,PPARD,PRKCB,SERPINA3 |
| CREBBP | transcription regulator | 0.0107 | 7 | ANXA6,CYP24A1,DNAJB4,PCK1,PPARGC1A,SERPINA3,SLC18A2 |
| PRL | cytokine | 0.00272 | 7 | ANXA5,ENO1,SERPINA3,SND1,SPARC,TP53,VIM |
| BDNF | growth factor | 0.00408 | 7 | ANXA5,CASQ1,CNR1,SNAP25,SPARC,TUBB3,VIM |
| SIRT1 | transcription regulator | 0.0173 | 7 | ANXA4,CNR1,PCK1,PML,PPARGC1A,SP7,TP53 |
| EDN1 | cytokine | 0.000235 | 7 | ANXA4,ANXA5,ANXA6,PRKCB,TP53,VCL,VIM |
| sulforafan | chemical drug | 2.87E-05 | 7 | ALDH1A1,MSTN,Mt1,PML,TP53,TXNRD1,VIM |
| gentamicin | chemical drug | 0.00499 | 7 | ALDH1A1,DNAJB4,EPHX2,ETFDH,S100A1,TP53,VAT1 |
| RXRA | ligand-dependent nuclear receptor | 0.00249 | 7 | ALDH1A1,APOA1,APOE,CYP24A1,PCK1,PPARD,SLC2A4 |
| HNF1A | transcription regulator | 0.026 | 7 | AHSG,ANXA4,HPX,OGDH,PCK1,PDHX,SERPINA1 |
| FOXO3 | transcription regulator | 0.00531 | 7 | AHR,MSTN,Mt1,PPARGC1A,PRCP,TP53,VIM |
| PDGF BB | complex | 0.00492 | 7 | AHR,MAP2,Mt1,PPARD,SERPINA3,TP53,YWHAQ |
| HGF | growth factor | 0.0323 | 7 | AHR,IL15RA,PCK1,PPARD,SP7,TP53,VIM |
| Z-LLL-CHO | chemical - protease inhibitor | 0.00589 | 7 | AHR,CASR,ENO1,FKBP4,SLC2A4,SP7,TP53 |
| IL15 | cytokine | 0.0211 | 7 | AHR,APBA1,ENO1,IL15RA,PPARD,PPARGC1A,TXNRD1 |
| metformin | chemical drug | 0.00104 | 7 | AHR,AHSG,MSTN,PCK1,PPARGC1A,SLC2A4,TP53 |
| GnRH analog | biologic drug | 0.00376 | 7 | ACP1,AKAP10,MAP2,PRCP,TP53,VAT1,VCL |
| DNMT3A | enzyme | 0.000179 | 7 | ACE,CNR1,Mt1,MYL3,PRKCB,S100A1,SPARC |
| Nr1h | group | 4.19E-05 | 7 | ACE,ALDH1A1,APOE,PML,PPARGC1A,SLC2A4,TP53 |
| nitric oxide | chemical - endogenous mammalian | 4.72E-05 | 7 | ACE,AHR,Mt1,PPARGC1A,TP53,TXNRD1,VCL |
| Vegf | group | 0.0378 | 7 | ACE,AHR,IL15RA,PPARD,PPARGC1A,PRKCB,VIM |
| TEAD1 | transcription regulator | 4.72E-05 | 7 | ACADVL,ATP5F1B,ETFDH,NDUFS3,PPARD,PPARGC1A,SDHA |
| ESRRA | transcription regulator | 0.00116 | 7 | ACADVL,ATP5F1B,ENO1,ETFDH,PDP1,PPARGC1A,PRCP |
| fenofibrate | chemical drug | 0.00109 | 7 | ACADVL,APOA1,MSTN,PDHB,PPARGC1A,SLC2A4,TP53 |
| methotrexate | chemical drug | 0.000727 | 7 | ACADVL,ANXA4,Mt1,PCK1,PPARD,PRKCB,TP53 |
| SCD | enzyme | 0.000134 | 7 | ACADVL,AHSG,ALDH1A1,PCK1,PPARD,PPARGC1A,SLC2A4 |
| tetrachlorodibenzodioxin | chemical toxicant | 0.0326 | 7 | AARS1,AHR,CALN1,PCK1,PRKCB,SLC2A4,TP53 |
| NRG1 | growth factor | 0.00583 | 6 | PFKFB1,SDHA,SLC2A4,TF,VCL,VIM |
| melatonin | chemical - endogenous mammalian | 0.000465 | 6 | Mt1,SLC2A4,SP7,SPARC,TP53,VIM |
| RUNX1 | transcription regulator | 0.00593 | 6 | Mt1,MYH10,PRKCB,SLC22A4,SP7,TP53 |
| HDAC4 | transcription regulator | 0.000217 | 6 | GABRA5,GABRG3,MYH10,PRKCB,SLC2A4,SNAP25 |
| 15-deoxy-delta-12,14 -PGJ 2 | chemical - endogenous mammalian | 0.0032 | 6 | FKBP4,SERPINA1,SLC2A4,TP53,TUBB3,TXNRD1 |
| ANGPT2 | growth factor | 0.0019 | 6 | DNAJB4,EPHX2,FKBP4,TP53,TUBB3,VIM |
| RNA polymerase II | complex | 0.00714 | 6 | CYP24A1,ENO1,KSR2,Mt1,PCK1,RPL13A |
| Esrra | transcription regulator | 6.62E-05 | 6 | ATP5F1B,CNR1,ENO1,NDUFS3,PCK1,PPARGC1A |
| CD 437 | chemical drug | 0.00139 | 6 | ATP5F1B,CCT6A,CNR1,ENO1,RPL13A,TP53 |
| RICTOR | other | 0.00738 | 6 | ATP5F1B,ATP6V1A,NDUFS3,PRKCB,RPL13A,SDHA |
| LDLR | transporter | 0.00326 | 6 | APOE,PCK1,PCYT2,PPARGC1A,SLC2A4,SP7 |
| bucladesine | chemical toxicant | 0.0215 | 6 | APOE,Mt1,PCK1,SLC2A4,SNAP25,TF |
| cholesterol | chemical - endogenous mammalian | 0.00333 | 6 | APOE,MAP2,PCYT2,PPARD,PPARGC1A,SLC2A4 |
| MECP2 | transcription regulator | 0.00105 | 6 | APOE,EFNA5,GABRA5,Mt1,TPH2,VIM |
| HDAC1 | transcription regulator | 0.0075 | 6 | APOA1,Mt1,PCK1,PPARGC1A,TP53,TUBB3 |
| NR1H4 | ligand-dependent nuclear receptor | 0.00271 | 6 | APOA1,APOE,PCK1,PPARGC1A,SDHA,SERPINA1 |
| NR1H3 | ligand-dependent nuclear receptor | 0.00277 | 6 | APOA1,APOE,Mt1,PFKFB1,PPARGC1A,SLC2A4 |
| EZH2 | transcription regulator | 0.0388 | 6 | ANXA6,CHRM3,CNR1,SERPINA1,SP7,TP53 |
| P38 MAPK | group | 0.0247 | 6 | ANXA5,PCK1,PML,PPARGC1A,SP7,TP53 |
| KLF4 | transcription regulator | 0.0142 | 6 | ALDH1A1,SERPINA1,TF,TP53,TXNRD1,VIM |
| MYCN | transcription regulator | 0.0149 | 6 | ALDH1A1,EFNA5,RPL13A,SPARC,TP53,VIM |
| aflatoxin B1 | chemical - endogenous non-mammalian | 0.00838 | 6 | ALDH1A1,CHRM3,ELAVL2,SPARC,TP53,TXNRD1 |
| E. coli B5 lipopolysaccharide | chemical - endogenous non-mammalian | 0.00479 | 6 | AHSG,APOA1,IL15RA,Mup1 (includes others),PPARGC1A,RDX |
| Pln | other | 1.08E-06 | 6 | AHSG,APOA1,CALU,ENO1,PRKCB,VIM |
| PLN | transporter | 2.15E-06 | 6 | AHSG,APOA1,CALU,ENO1,PRKCB,VIM |
| pioglitazone | chemical drug | 0.00126 | 6 | AHSG,APOA1,APOE,PPARD,PPARGC1A,SLC2A4 |
| simvastatin | chemical drug | 0.00218 | 6 | AHSG,APOA1,APOE,MYH10,SLC2A4,VIM |
| benzo(a)pyrene | chemical toxicant | 0.00218 | 6 | AHR,APOA1,SERPINA1,TF,TP53,TXNRD1 |
| cycloheximide | chemical reagent | 0.0279 | 6 | AHR,APOA1,CYP24A1,KSR2,Mt1,TP53 |
| NR1I2 | ligand-dependent nuclear receptor | 0.00105 | 6 | AHR,ALDH1A1,CYP24A1,Mup1 (includes others),PAPSS2,TF |
| NR1I3 | ligand-dependent nuclear receptor | 4.67E-05 | 6 | AHR,ALDH1A1,APOA1,Mt1,PAPSS2,PCK1 |
| fatty acid | chemical - endogenous mammalian | 9.17E-05 | 6 | AHR,AHSG,APOA1,PPARD,PPARGC1A,SLC2A4 |
| SNAI1 | transcription regulator | 0.000409 | 6 | ACTR1A,CFL2,SP7,SPARC,TF,VIM |
| KDM1A | enzyme | 0.0239 | 6 | ACP1,IQGAP2,MAP2,SNAP25,VCL,VIM |
| STK11 | kinase | 0.00949 | 6 | ACE,CCT6A,ENO1,LUM,PPARGC1A,TXNRD1 |
| FN1 | enzyme | 0.00177 | 6 | ACE,APOE,RDX,SPARC,VCL,VIM |
| genistein | chemical drug | 0.0355 | 6 | ACE,APOA1,ATP6V1A,Mt1,OGDH,TP53 |
| nicotine | chemical drug | 0.0026 | 6 | ACE,AHR,CHRM3,ENO1,PPARD,TP53 |
| INS | other | 0.000465 | 6 | ACADVL,APOA1,PCK1,PPARGC1A,SLC2A4,TP53 |
| POR | enzyme | 0.000734 | 6 | ACADVL,AHR,ALDH1A1,ANXA5,Mt1,SERPINA3 |
| PTP4A1 | phosphatase | 7.63E-05 | 6 | AARS1,ANXA5,SDHA,SPARC,TP53,VIM |
| FOXC2 | transcription regulator | 0.000207 | 5 | PPARGC1A,SLC2A4,SP7,TP53,VIM |
| BMP4 | growth factor | 0.0139 | 5 | PPARD,PRKCB,SP7,SPARC,TP53 |
| lysophosphatidic acid | chemical - other | 0.000321 | 5 | PCK1,PFKFB1,TF,TP53,VIM |
| cyclic AMP | chemical - endogenous mammalian | 0.0226 | 5 | PCK1,PFKFB1,PPARGC1A,TF,TP53 |
| OGT | enzyme | 0.000497 | 5 | OGDH,PDHX,PPARGC1A,TP53,VIM |
| NOS2 | enzyme | 0.00547 | 5 | MYL3,PPARGC1A,PRDX6,SERPINA3,SLC2A4 |
| FST | other | 9.38E-05 | 5 | MSTN,Mup1 (includes others),PPARGC1A,SERPINA1,SLC2A4 |
| Sb202190 | chemical drug | 0.00138 | 5 | MAPKAP1,PPARGC1A,SP7,TP53,VIM |
| DICER1 | enzyme | 0.0398 | 5 | MAP2,SERPINA1,SPARC,TP53,VIM |
| PDX1 | transcription regulator | 0.00609 | 5 | MAP2,Mt1,PAPSS2,SDHA,XKR4 |
| IL10RA | transmembrane receptor | 0.0384 | 5 | IL15RA,LUM,PCK1,PPARGC1A,SPARC |
| CXCL12 | cytokine | 0.00795 | 5 | IFNAR2,SP7,TF,TP53,VIM |
| HNRNPA2B1 | other | 0.00182 | 5 | GABRA5,PPARGC1A,SERPINA1,SLC2A4,SNAP25 |
| sodium arsenite | chemical drug | 5.94E-05 | 5 | ENO1,PCK1,SLC18A2,SLC2A4,TP53 |
| RORC | ligand-dependent nuclear receptor | 0.00649 | 5 | ENO1,Mt1,Mup1 (includes others),PCK1,SPARC |
| REST | transcription regulator | 0.00301 | 5 | EFNA5,SNAP25,TPH2,TUBB3,XKR4 |
| PGR | ligand-dependent nuclear receptor | 0.0346 | 5 | DNAJB4,PAPSS2,SERPINA1,SNAP25,VCL |
| MRTFA | transcription regulator | 0.00512 | 5 | DNAJB4,MAP2,MYH10,SPOCK3,VCL |
| RAS | group | 0.00117 | 5 | CYP24A1,PML,PPARD,TP53,VIM |
| cholecalciferol | chemical - endogenous mammalian | 0.00121 | 5 | CUBN,CYP24A1,PPARGC1A,PRKCB,SP7 |
| GNA12 | enzyme | 0.000106 | 5 | CFL2,RDX,TXNRD1,VCL,VIM |
| DNMT3B | enzyme | 0.00406 | 5 | CASQ1,CNR1,MYL3,PRKCB,S100A1 |
| topotecan | chemical drug | 0.00535 | 5 | CAMTA1,CNTNAP2,ENO1,TP53,VCL |
| ETV6-RUNX1 | fusion gene/product | 0.0402 | 5 | CALN1,IFI30,IFNAR2,IL15RA,PML |
| arsenic trioxide | chemical drug | 0.0494 | 5 | ATP6V1A,PFKFB1,PML,TP53,TXNRD1 |
| PRKAG3 | kinase | 0.00309 | 5 | ATP6V1A,IFI30,PPARGC1A,RPL13A,SH3KBP1 |
| HDAC5 | transcription regulator | 0.000158 | 5 | ATP5F1B,PPARGC1A,SLC2A4,TP53,TUBB3 |
| Perm1 | other | 1.62E-06 | 5 | ATP5F1B,NDUFS3,PDHB,PPARGC1A,SLC2A4 |
| torin1 | chemical reagent | 0.00559 | 5 | ATP5F1B,ENO1,PDHB,RPL13A,VIM |
| UQCC3 | other | 0.00167 | 5 | ATP5F1B,ATP6V1A,ENO1,NDUFS3,SDHA |
| IDH1 | enzyme | 2.81E-05 | 5 | APOE,PCK1,SLC2A4,TP53,VIM |
| oleic acid | chemical - endogenous mammalian | 0.00194 | 5 | APOE,Mt1,PCK1,PPARGC1A,SLC2A4 |
| bexarotene | chemical drug | 0.00609 | 5 | APOE,EFNA5,SLC18A2,SPARC,TUBB3 |
| Rxr | group | 0.000585 | 5 | APOE,CYP24A1,EFNA5,PCK1,TPH2 |
| kainic acid | chemical toxicant | 0.00512 | 5 | APOE,CNR1,CUBN,GABRA5,TP53 |
| NFKB1 | transcription regulator | 0.0354 | 5 | APOE,CASR,IFNAR2,SP7,TP53 |
| MITF | transcription regulator | 0.0198 | 5 | APOE,CA14,PPARGC1A,TP53,VAT1 |
| THRA | ligand-dependent nuclear receptor | 0.00211 | 5 | APOA1,PCK1,PPARGC1A,SLC2A4,TP53 |
| actinomycin D | biologic drug | 0.0141 | 5 | APOA1,CYP24A1,Mt1,TF,TP53 |
| mir-8 | microRNA | 0.00691 | 5 | APOA1,CFL2,MYH10,TP53,VIM |
| estrogen | chemical drug | 0.0302 | 5 | APOA1,APOE,SERPINA3,SPARC,TP53 |
| MAPK8 | kinase | 0.00457 | 5 | APOA1,APOE,PPARD,PPARGC1A,TP53 |
| SP3 | transcription regulator | 0.00927 | 5 | APBA1,SPARC,TP53,TXNRD1,VIM |
| IL5 | cytokine | 0.0291 | 5 | ANXA6,ENO1,IFI30,LUM,VIM |
| MKNK1 | kinase | 0.00416 | 5 | ANXA5,SNAP25,SPARC,TF,VIM |
| cadmium chloride | chemical toxicant | 0.000253 | 5 | ANXA5,Mt1,Mup1 (includes others),TP53,VIM |
| CX3CR1 | G-protein coupled receptor | 0.00397 | 5 | ANXA5,EXOC4,PDHB,TF,WDPCP |
| PAX3-FOXO1 | fusion gene/product | 0.0269 | 5 | ANXA5,CCT6A,ENO1,MYH10,VIM |
| alitretinoin | chemical drug | 0.00342 | 5 | ANXA5,APOA1,CYP24A1,PPARGC1A,VIM |
| CST5 | other | 0.0245 | 5 | ANXA5,ANXA6,DNAJB4,IQGAP2,VIM |
| IL1A | cytokine | 0.0176 | 5 | ALDH1A1,SERPINA1,SERPINA3,SPARC,VIM |
| PKD1 | ion channel | 0.00826 | 5 | ALDH1A1,PCK1,PML,PRKCB,SP7 |
| LIPE | enzyme | 0.00182 | 5 | ALDH1A1,PCK1,PFKFB1,PPARGC1A,SLC2A4 |
| elaidic acid | chemical - endogenous mammalian | 0.000229 | 5 | ALDH1A1,APOE,ATP5F1B,HPX,PCYT2 |
| ARNT | transcription regulator | 0.00309 | 5 | AHR,ENO1,NDUFS3,TP53,VIM |
| KLF6 | transcription regulator | 0.0235 | 5 | AHR,ANXA5,IFI30,IFNAR2,IL15RA |
| EPO | cytokine | 0.0354 | 5 | ADAMTS10,PRKCB,TF,TP53,TPH2 |
| MEF2C | transcription regulator | 0.00121 | 5 | ACTN2,PPARGC1A,SLC2A4,SP7,VIM |
| fluoride | chemical - endogenous mammalian | 0.000134 | 5 | ACTN2,MYH10,SP7,TUBB3,YWHAQ |
| IGF1R | transmembrane receptor | 0.0342 | 5 | ACP1,ATP5F1B,CALU,SP7,TP53 |
| N-nitro-L-arginine methyl ester | chemical drug | 0.000351 | 5 | ACE,PPARD,PPARGC1A,SLC2A4,TP53 |
| isobutylmethylxanthine | chemical toxicant | 0.00962 | 5 | ACE,Mt1,PPARD,PPARGC1A,SDHA |
| thyroid hormone | chemical - endogenous mammalian | 0.00489 | 5 | ACE,APOE,PCK1,PML,VIM |
| mifepristone | chemical drug | 0.0479 | 5 | ACE,APOE,Mt1,SNAP25,TP53 |
| MAP4K4 | kinase | 0.000685 | 5 | ACADVL,OGDH,PAPSS2,PDHX,SLC2A4 |
| ATP5IF1 | other | 3.85E-05 | 5 | ACADVL,ETFDH,PCK1,SDHA,TXNRD1 |
| PPARGC1B | transcription regulator | 0.000068 | 5 | ACADVL,ATP5F1B,ENO1,PCK1,SLC2A4 |
| heparin | chemical - endogenous mammalian | 0.000113 | 5 | ACADVL,APOE,SDHA,SP7,VIM |
| HNF1B | transcription regulator | 0.000826 | 5 | ACADVL,ANXA4,LUM,SERPINA1,SPARC |
| MYL2 | other | 2.17E-06 | 5 | ACADVL,ACTN2,ANXA5,ANXA6,HPX |
| glutamine | chemical - endogenous mammalian | 0.000518 | 5 | AARS1,PCK1,PPARGC1A,TP53,VIM |
| crizotinib | chemical drug | 6.61E-05 | 4 | PPARGC1A,SDHA,SPARC,VIM |
| wortmannin | chemical drug | 0.04 | 4 | PPARD,SLC2A4,TP53,VIM |
| S-nitroso-N-acetyl-DL-penicillamine | chemical reagent | 0.000439 | 4 | PPARD,PPARGC1A,SLC2A4,TP53 |
| SFRP1 | transmembrane receptor | 0.000314 | 4 | PCK1,SLC2A4,TP53,VIM |
| PRKAA | group | 0.000337 | 4 | PCK1,PPARGC1A,TP53,VIM |
| GCG | other | 0.00156 | 4 | PCK1,PPARGC1A,SLC2A4,TP53 |
| MLXIPL | transcription regulator | 0.0116 | 4 | PCK1,PPARGC1A,RPL13A,SLC2A4 |
| APC | enzyme | 0.0206 | 4 | OGDH,PPARD,PRKCB,TP53 |
| PRKN | enzyme | 0.00227 | 4 | NDUFS3,PRDX6,SDHA,TP53 |
| RBM20 | other | 0.00428 | 4 | MYL3,PRDX6,RPL13A,SPARC |
| NRF1 | transcription regulator | 0.000833 | 4 | Mt1,SDHA,TP53,VIM |
| Hbb-b1 | transporter | 0.00227 | 4 | Mt1,NDUFS3,SDHA,TF |
| HFE | transmembrane receptor | 0.00581 | 4 | Mt1,Mup1 (includes others),SERPINA3,TF |
| diethylnitrosamine | chemical toxicant | 0.0116 | 4 | Mt1,Mup1 (includes others),PPARGC1A,TP53 |
| AICAR | chemical - endogenous mammalian | 0.00501 | 4 | MSTN,PPARD,PPARGC1A,TP53 |
| DSCAM | other | 0.00245 | 4 | MAP2,PRKCB,TUBB3,VIM |
| Ngf | group | 0.000925 | 4 | MAP2,PPARGC1A,TP53,TUBB3 |
| L-glutamic acid | chemical - endogenous mammalian | 0.00939 | 4 | MAP2,PPARGC1A,TF,TP53 |
| bisphenol A | chemical - endogenous mammalian | 0.0306 | 4 | GABRG3,SLC2A4,TP53,TPH2 |
| FOXA1 | transcription regulator | 0.0174 | 4 | FKBP4,PCK1,PFKFB1,TF |
| BMP2 | growth factor | 0.0465 | 4 | EPHX2,SP7,SPARC,TUBB3 |
| KDM8 | enzyme | 0.000925 | 4 | ENO1,OGDH,PDHB,SDHA |
| PCGEM1 | other | 0.00011 | 4 | ENO1,OGDH,PDHB,SDHA |
| desmopressin | biologic drug | 0.00235 | 4 | ENO1,MYH10,SLC2A4,YWHAQ |
| ELOVL3 | enzyme | 0.00079 | 4 | ENO1,IFI30,IQGAP2,LUM |
| FMR1 | translation regulator | 0.0148 | 4 | ENO1,GABRA5,SNAP25,TF |
| FEV | transcription regulator | 0.00177 | 4 | EFNA5,SLC18A2,TPH2,VIM |
| SOX4 | transcription regulator | 0.0356 | 4 | EFNA5,IFI30,Mt1,VIM |
| FGFR1 | kinase | 0.00363 | 4 | DNAJC1,SPARC,TUBB3,VIM |
| MRTFB | transcription regulator | 0.0191 | 4 | DNAJB4,MAP2,MYH10,VCL |
| lactacystin | chemical - protease inhibitor | 0.0315 | 4 | CYP24A1,SP7,TP53,TXNRD1 |
| GW501516 | chemical drug | 0.00376 | 4 | CYP24A1,PPARD,PPARGC1A,TP53 |
| deferoxamine | chemical drug | 0.04 | 4 | CYP24A1,ENO1,PFKFB1,TP53 |
| 1,25-dihydroxyvitamin D | chemical drug | 0.000632 | 4 | CUBN,CYP24A1,MSTN,VIM |
| 2-arachidonoylglycerol | chemical - endogenous mammalian | 2.61E-06 | 4 | CNR1,PCK1,PPARD,PPARGC1A |
| 1-methyl-4-phenyl-1,2,3,6-tetrahydropyridine | chemical toxicant | 0.00363 | 4 | CNR1,EPHX2,MAP2,TP53 |
| KLF11 | transcription regulator | 0.00786 | 4 | CHRM3,PCK1,PPARGC1A,SERPINA1 |
| TRIM24 | transcription regulator | 0.00456 | 4 | CASR,CYP24A1,MYH10,TP53 |
| TERT | enzyme | 0.0301 | 4 | CALU,IQGAP2,TP53,VIM |
| SUZ12 | enzyme | 0.00871 | 4 | CALU,CHRM3,CNR1,ILF3 |
| tetrodotoxin | chemical drug | 0.00986 | 4 | CADM2,LUM,Mt1,PPARGC1A |
| corticosterone | chemical - endogenous mammalian | 0.0174 | 4 | CA14,FKBP4,PPARGC1A,TPH2 |
| FLCN | other | 0.00828 | 4 | ATP6V1A,ENO1,PPARGC1A,SPARC |
| ESRRG | ligand-dependent nuclear receptor | 0.00156 | 4 | ATP5F1B,ENO1,PCK1,PDHB |
| ST1926 | chemical drug | 0.0157 | 4 | ATP5F1B,CCT6A,ENO1,TP53 |
| IND S7 | chemical - kinase inhibitor | 2.14E-05 | 4 | ATP5F1B,CALU,ENO1,VIM |
| MEL S3 | chemical - kinase inhibitor | 2.47E-05 | 4 | ATP5F1B,CALU,ENO1,VIM |
| EIF2B5 | translation regulator | 2.83E-05 | 4 | APOE,SERPINA3,TF,VIM |
| VCAN | other | 0.0174 | 4 | APOE,MYH10,TP53,VCL |
| docosahexaenoic acid | chemical drug | 0.023 | 4 | APOE,MSTN,PCK1,SLC2A4 |
| ASPSCR1-TFE3 | fusion gene/product | 0.00284 | 4 | APOE,IFI30,PPARGC1A,VAT1 |
| MBD3 | other | 0.00143 | 4 | APOE,EFNA5,Mt1,PPARD |
| CNR1 | G-protein coupled receptor | 0.013 | 4 | APOE,CNR1,MSTN,SLC2A4 |
| 4-methylnitrosoamino-1-(3-pyridinyl)-1-butanone | chemical toxicant | 0.000157 | 4 | APOA1,SERPINA1,TF,TP53 |
| phenobarbital | chemical drug | 0.0017 | 4 | APOA1,PAPSS2,PCK1,TP53 |
| 1,4-bis[2-(3,5-dichloropyridyloxy)]benzene | chemical toxicant | 0.00548 | 4 | APOA1,Mt1,PAPSS2,PCK1 |
| sodium tungstate | chemical drug | 4.17E-05 | 4 | APOA1,ENO1,PDHB,SNAP25 |
| CD36 | transmembrane receptor | 0.00351 | 4 | APOA1,ATP5F1B,PPARGC1A,SLC2A4 |
| ABCA1 | transporter | 0.000361 | 4 | APOA1,APOE,PPARGC1A,SLC2A4 |
| GW3965 | chemical reagent | 0.00669 | 4 | APOA1,APOE,PCYT2,SLC2A4 |
| chenodeoxycholic acid | chemical - endogenous mammalian | 0.00201 | 4 | APOA1,APOE,PCK1,TXNRD1 |
| GW 4064 | chemical toxicant | 0.000748 | 4 | APOA1,APOE,PCK1,PPARGC1A |
| APOA1 | transporter | 0.000468 | 4 | APOA1,APOE,ATP5F1B,PPARGC1A |
| CD38 | enzyme | 0.0174 | 4 | ANXA6,IFI30,PPARGC1A,VIM |
| ouabain | chemical drug | 0.000144 | 4 | ANXA5,TP53,VIM,YWHAQ |
| Jnk | group | 0.0412 | 4 | ANXA5,PPARD,SP7,TP53 |
| staurosporine | chemical drug | 0.00615 | 4 | ANXA5,PCK1,SERPINA3,TP53 |
| UBA1 | enzyme | 1.13E-05 | 4 | ANXA5,APOE,TP53,VIM |
| TWIST1 | transcription regulator | 0.0177 | 4 | ALDH1A1,TF,TP53,VIM |
| VHL | transcription regulator | 0.0116 | 4 | ALDH1A1,SPARC,TP53,VIM |
| ID1 | transcription regulator | 0.00108 | 4 | ALDH1A1,PPARGC1A,TF,VIM |
| LRP6 | transmembrane receptor | 0.000386 | 4 | ALDH1A1,PPARGC1A,SP7,VIM |
| DIO2 | enzyme | 0.0145 | 4 | ALDH1A1,LUM,PDP1,PPARGC1A |
| TCF | group | 0.00209 | 4 | ALDH1A1,CYP24A1,SERPINA1,SERPINA3 |
| FGF19 | growth factor | 0.00193 | 4 | ALDH1A1,APOE,SERPINA1,SERPINA3 |
| Hmgn3 | other | 3.24E-05 | 4 | AHSG,APOA1,Mup1 (includes others),SERPINA1 |
| cigarette smoke | chemical toxicant | 0.04 | 4 | AHR,SERPINA3,TP53,TXNRD1 |
| ITK | kinase | 0.00177 | 4 | AHR,PRKCB,TF,TP53 |
| NCOA1 | transcription regulator | 0.0127 | 4 | AHR,PCK1,PMM2,PPARD |
| HAVCR1 | other | 0.00388 | 4 | AHR,MYH10,SPARC,TP53 |
| ATP | chemical - endogenous mammalian | 0.00669 | 4 | AHR,CASR,TP53,TUBB3 |
| PRNP | other | 0.00136 | 4 | AHR,APOE,SNAP25,TP53 |
| MAP2K1 | kinase | 0.0296 | 4 | AHR,APOE,ENO1,SERPINA1 |
| aspirin | chemical drug | 0.021 | 4 | AHR,APOA1,PPARD,TP53 |
| Pkc(s) | group | 0.0423 | 4 | AHR,APOA1,CYP24A1,S100A1 |
| miR-16-5p (and other miRNAs w/seed AGCAGCA) | mature microRNA | 0.0383 | 4 | ACTR1A,CFL2,DNAJB4,DTD1 |
| mir-802 | microRNA | 0.0116 | 4 | ACP1,CFL2,RDX,VIM |
| ERG | transcription regulator | 0.0465 | 4 | ACOT8,CNR1,SPARC,VIM |
| JAK2 | kinase | 0.0136 | 4 | ACE,PRKCB,TF,TP53 |
| H89 | chemical drug | 0.0167 | 4 | ACE,PPARD,PPARGC1A,SNAP25 |
| losartan potassium | chemical drug | 0.0111 | 4 | ACE,MAP2,PPARGC1A,TUBB3 |
| aldosterone | chemical - endogenous mammalian | 0.026 | 4 | ACE,CYP24A1,PPARGC1A,VIM |
| NR1H2 | ligand-dependent nuclear receptor | 0.00351 | 4 | ACE,APOE,PFKFB1,SLC2A4 |
| cholic acid | chemical - endogenous mammalian | 0.00108 | 4 | ACE,APOA1,PCK1,PPARGC1A |
| okadaic acid | chemical toxicant | 0.00245 | 4 | ACE,APOA1,CYP24A1,SP7 |
| MAPK3 | kinase | 0.00363 | 4 | ACE,APOA1,CYP24A1,PPARGC1A |
| CREM | transcription regulator | 0.0151 | 4 | ACE,ANXA4,APOE,SLC2A4 |
| PNPLA2 | enzyme | 0.00156 | 4 | ACADVL,PCK1,PPARGC1A,TP53 |
| cyclopropanecarboxylic acid | chemical reagent | 6.61E-05 | 4 | ACADVL,PCK1,PDHB,TP53 |
| ACOX1 | enzyme | 0.0108 | 4 | ACADVL,Mup1 (includes others),SERPINA1,TP53 |
| bezafibrate | chemical drug | 0.00284 | 4 | ACADVL,APOA1,APOE,PPARGC1A |
| gemfibrozil | chemical drug | 0.000878 | 4 | ACADVL,APOA1,APOE,PPARD |
| UCP1 | transporter | 0.0164 | 4 | AARS1,NDUFS3,PCK1,SND1 |
| mir-29 | microRNA | 0.0199 | 3 | SPARC,TP53,VIM |
| DKK1 | growth factor | 0.00893 | 3 | SP7,SPARC,TP53 |
| mir-221 | microRNA | 0.00115 | 3 | SLC2A4,TP53,TUBB3 |
| PPP3CA | phosphatase | 0.0178 | 3 | SLC2A4,TF,VIM |
| CCN5 | growth factor | 0.00381 | 3 | SLC2A4,SPARC,VIM |
| Nuclear factor 1 | group | 0.000803 | 3 | SERPINA3,SLC2A4,TP53 |
| TGFA | growth factor | 0.0131 | 3 | SERPINA1,TP53,VIM |
| arachidonic acid | chemical - endogenous mammalian | 0.025 | 3 | PRKCB,SLC2A4,VIM |
| cuprizone | chemical toxicant | 0.0263 | 3 | PRDX6,SERPINA3,VIM |
| PC-SPES | chemical drug | 0.0199 | 3 | PRDX6,SDHA,VIM |
| SIRT3 | enzyme | 0.00733 | 3 | PPARGC1A,TP53,VIM |
| PRKAA2 | kinase | 0.0495 | 3 | PPARGC1A,TP53,VIM |
| HIPK2 | kinase | 0.0104 | 3 | PPARGC1A,TP53,VIM |
| caffeine | chemical drug | 0.00827 | 3 | PPARGC1A,TP53,VIM |
| VEGFB | growth factor | 0.00445 | 3 | PPARGC1A,TP53,TXNRD1 |
| ADORA2A | G-protein coupled receptor | 0.0495 | 3 | PPARGC1A,SPARC,TUBB3 |
| MSTN | growth factor | 0.00928 | 3 | PPARGC1A,SP7,VIM |
| MEF2D | transcription regulator | 0.0104 | 3 | PPARGC1A,SLC2A4,VIM |
| ADIPOR1 | transmembrane receptor | 0.00106 | 3 | PPARGC1A,SLC2A4,TP53 |
| DDIT3 | transcription regulator | 0.0404 | 3 | PPARGC1A,SERPINA1,VIM |
| SURF1 | enzyme | 0.00007 | 3 | PPARGC1A,SDHA,SLC2A4 |
| TNFSF12 | cytokine | 0.0256 | 3 | PPARGC1A,SDHA,SLC2A4 |
| MAP2K3 | kinase | 0.0086 | 3 | PPARGC1A,PRDX6,SLC2A4 |
| berberine | chemical drug | 0.0358 | 3 | PPARD,TP53,VIM |
| maslinic acid | chemical - endogenous non-mammalian | 0.0204 | 3 | PPARD,PRKCB,TP53 |
| PLCL2 | enzyme | 4.27E-05 | 3 | PPARD,PPARGC1A,SPARC |
| PLCL1 | enzyme | 0.000129 | 3 | PPARD,PPARGC1A,SPARC |
| PRKD | group | 0.000966 | 3 | PPARD,PPARGC1A,SLC2A4 |
| guanidinopropionic acid | chemical - endogenous non-mammalian | 0.00305 | 3 | PPARD,PPARGC1A,SDHA |
| NDRG1 | kinase | 0.00703 | 3 | PML,TP53,VIM |
| PRKCA | kinase | 0.038 | 3 | PML,SP7,TP53 |
| UBE2I | enzyme | 0.00146 | 3 | PML,PPARGC1A,SLC2A4 |
| PIN1 | enzyme | 0.0127 | 3 | PML,PPARD,PPARGC1A |
| E2f | group | 0.038 | 3 | PFKFB1,RBBP5,TP53 |
| ARNTL | transcription regulator | 0.0115 | 3 | PDP1,SLC2A4,TP53 |
| SMYD1 | transcription regulator | 0.00381 | 3 | PDHB,PPARGC1A,SDHA |
| fisetin | chemical drug | 0.00196 | 3 | PCK1,TP53,VIM |
| NEDD9 | other | 0.0158 | 3 | PCK1,TF,VIM |
| epicatechin | chemical drug | 0.0204 | 3 | PCK1,SNAP25,SPARC |
| FGF21 | growth factor | 0.0119 | 3 | PCK1,PPARGC1A,VIM |
| dorsomorphin | chemical - kinase inhibitor | 0.0111 | 3 | PCK1,PPARGC1A,TP53 |
| SOCS3 | phosphatase | 0.0204 | 3 | PCK1,PPARGC1A,TP53 |
| FOXA3 | transcription regulator | 0.00158 | 3 | PCK1,PPARGC1A,TF |
| ATF2 | transcription regulator | 0.0183 | 3 | PCK1,PPARGC1A,TF |
| IRS1 | enzyme | 0.0436 | 3 | PCK1,PPARGC1A,SLC2A4 |
| ZNF746 | transcription regulator | 0.000011 | 3 | PCK1,PPARGC1A,SDHA |
| retinol | chemical drug | 0.000327 | 3 | PCK1,PPARD,TF |
| ATF3 | transcription regulator | 0.0486 | 3 | PAPSS2,PCK1,TP53 |
| BMP6 | growth factor | 0.0238 | 3 | MYH10,TXNRD1,VCL |
| MASTL | kinase | 0.00287 | 3 | MYH10,STK24,VCL |
| methylmercury | chemical toxicant | 0.00703 | 3 | Mt1,SPARC,YWHAQ |
| TSH | complex | 0.00733 | 3 | Mt1,PPARD,SLC2A4 |
| USF1 | transcription regulator | 0.014 | 3 | Mt1,PCK1,TP53 |
| HBA1/HBA2 | transporter | 0.00224 | 3 | Mt1,NDUFS3,SDHA |
| MMP9 | peptidase | 0.0322 | 3 | MSTN,SERPINA1,VIM |
| phenylephrine | chemical drug | 0.0168 | 3 | MSTN,PRKCB,SLC2A4 |
| NR4A3 | ligand-dependent nuclear receptor | 0.0163 | 3 | MSTN,PDP1,PPARGC1A |
| ethyl 2-{[(6,7-dimethyl-3-oxo-1,2,3,4-tetrahydro-2-quinoxalinyl)acetyl]amino}-4,5-dimethyl-3-thiophenecarboxylate | chemical reagent | 0.00007 | 3 | MAP2,TP53,TUBB3 |
| SPARC | other | 0.0295 | 3 | MAP2,SPARC,VIM |
| C1QA | other | 0.000966 | 3 | MAP2,SERPINA3,VIM |
| DSCAML1 | other | 0.0163 | 3 | MAP2,PRKCB,TUBB3 |
| GNA15 | enzyme | 0.0295 | 3 | MAP2,Mt1,SERPINA3 |
| BIRC5 | other | 0.00445 | 3 | LUM,TP53,VIM |
| MIF | cytokine | 0.0427 | 3 | IL15RA,TP53,TPH2 |
| EML4-ALK | fusion gene/product | 0.0149 | 3 | IL15RA,SPARC,TP53 |
| MET | kinase | 0.0477 | 3 | IL15RA,PPARGC1A,VIM |
| Ifn | group | 0.0358 | 3 | IL15RA,PML,TP53 |
| PLAU | peptidase | 0.00893 | 3 | IFNAR2,TP53,VIM |
| MACROH2A1 | other | 0.0127 | 3 | IFNAR2,SLC2A4,SPARC |
| Rhox5 | transcription regulator | 0.000803 | 3 | IFNAR2,PPARGC1A,SLC22A4 |
| CHKB | kinase | 0.00254 | 3 | ETFDH,PPARD,PPARGC1A |
| CCN2 | growth factor | 0.0282 | 3 | ENO1,SPARC,TP53 |
| KDM4B | enzyme | 0.000373 | 3 | ENO1,SDHA,VIM |
| TXNIP | other | 0.00565 | 3 | ENO1,PDP1,PPARGC1A |
| PKM | kinase | 0.014 | 3 | ENO1,PDHB,PPARGC1A |
| zinc | chemical drug | 0.0131 | 3 | ENO1,Mt1,TP53 |
| SFMBT1 | transcription regulator | 0.000327 | 3 | ENO1,MAP2,TUBB3 |
| KAT2A | enzyme | 0.0373 | 3 | EFNA5,PCK1,PPARGC1A |
| MYO6 | other | 0.0131 | 3 | DNAJB4,ERC1,OGDH |
| RRP1B | transcription regulator | 0.0275 | 3 | CYP24A1,HPX,RPL13A |
| verapamil | chemical drug | 0.00515 | 3 | CYP24A1,ENO1,TXNRD1 |
| cadmium | chemical toxicant | 0.00999 | 3 | CUBN,Mt1,TP53 |
| SOD2 | enzyme | 0.0238 | 3 | CTBP2,MAP2,TP53 |
| NR4A2 | ligand-dependent nuclear receptor | 0.0188 | 3 | CNR1,SLC18A2,SLC2A4 |
| Foxp1 | transcription regulator | 0.00158 | 3 | CNR1,PCK1,PPARGC1A |
| ALDH1A1 | enzyme | 0.000129 | 3 | CNR1,PCK1,PPARD |
| DNMT1 | enzyme | 0.0188 | 3 | CNR1,CYP24A1,Mt1 |
| thioridazine | chemical drug | 0.000214 | 3 | CHRM3,SERPINA3,TP53 |
| collagen type i (family) | group | 0.000593 | 3 | CFL2,LUM,SPARC |
| YWHAG | other | 0.00007 | 3 | CCT6A,PRDX6,TP53 |
| interferon beta-1a | biologic drug | 0.0373 | 3 | CCT6A,ENO1,SERPINA1 |
| CASR | G-protein coupled receptor | 0.0477 | 3 | CASR,SERPINA1,VCL |
| VitaminD3-VDR-RXR | complex | 0.00254 | 3 | CASR,CYP24A1,PPARD |
| CHD4 | enzyme | 0.00963 | 3 | CASQ1,ELAVL2,TP53 |
| GRIN3A | ion channel | 0.0302 | 3 | CADM2,LUM,Mt1 |
| TFEB | transcription regulator | 0.0188 | 3 | ATP6V1A,IFI30,PPARGC1A |
| PLA2R1 | transmembrane receptor | 0.000593 | 3 | ATP5F1B,NDUFS3,SDHA |
| ACLY | enzyme | 0.000474 | 3 | ATP5F1B,NDUFS3,PPARGC1A |
| TLE3 | other | 0.0144 | 3 | ATP5F1B,ETFDH,NDUFS3 |
| PARP | group | 2.35E-05 | 3 | APOE,TP53,VIM |
| Collagen type I (complex) | complex | 0.00795 | 3 | APOE,TP53,VIM |
| APOB | transporter | 0.00115 | 3 | APOE,SLC2A4,SP7 |
| daidzein | chemical drug | 0.0193 | 3 | APOE,OGDH,TP53 |
| XRCC5 | enzyme | 2.35E-05 | 3 | APOE,Mt1,TP53 |
| ascorbic acid | chemical - endogenous mammalian | 0.0238 | 3 | APOA1,SP7,SPARC |
| RETN | other | 0.00618 | 3 | APOA1,SLC2A4,TP53 |
| ZFHX3 | transcription regulator | 0.0163 | 3 | APOA1,SERPINA1,TUBB3 |
| LPL | enzyme | 0.00423 | 3 | APOA1,PPARGC1A,SLC2A4 |
| HSD11B1 | enzyme | 0.00158 | 3 | APOA1,PPARGC1A,SDHA |
| Mapk | group | 0.0263 | 3 | APOA1,PPARD,TUBB3 |
| linoleic acid | chemical - endogenous mammalian | 0.00963 | 3 | APOA1,PPARD,SLC2A4 |
| MEX3A | other | 0.00146 | 3 | APOA1,PCK1,PPARGC1A |
| NR2F2 | ligand-dependent nuclear receptor | 0.00764 | 3 | APOA1,PCK1,PPARGC1A |
| NR0B2 | ligand-dependent nuclear receptor | 0.0419 | 3 | APOA1,PCK1,PPARGC1A |
| NR5A2 | ligand-dependent nuclear receptor | 0.0436 | 3 | APOA1,Mt1,TP53 |
| MAT1A | enzyme | 0.000286 | 3 | APOA1,Mt1,PRDX6 |
| RXRG | ligand-dependent nuclear receptor | 0.00565 | 3 | APOA1,CYP24A1,PCK1 |
| LCAT | enzyme | 0.000373 | 3 | APOA1,APOE,PPARGC1A |
| MYRF | transcription regulator | 0.00795 | 3 | ANXA6,S100A1,SLC22A4 |
| UCHL1 | peptidase | 0.0107 | 3 | ANXA6,ATP5F1B,TP53 |
| inosine | chemical - endogenous mammalian | 0.00361 | 3 | ANXA5,Mt1,VIM |
| PSEN2 | peptidase | 0.0232 | 3 | ANXA5,ANXA6,TP53 |
| RGS4 | enzyme | 0.0178 | 3 | ANXA4,CALN1,PPARGC1A |
| MUC1 | other | 0.0149 | 3 | ALDH1A1,TXNRD1,VIM |
| 3-methyladenine | chemical toxicant | 0.00515 | 3 | ALDH1A1,TP53,VIM |
| ZEB1 | transcription regulator | 0.021 | 3 | ALDH1A1,TF,VIM |
| MAOA | enzyme | 0.00106 | 3 | ALDH1A1,SLC18A2,VIM |
| EN1 | transcription regulator | 0.000107 | 3 | ALDH1A1,SLC18A2,TUBB3 |
| SLC13A1 | transporter | 0.0173 | 3 | ALDH1A1,Mt1,PCK1 |
| propiconazole | chemical reagent | 0.0017 | 3 | ALDH1A1,CYP24A1,GABRG3 |
| triadimefon | chemical toxicant | 0.00323 | 3 | ALDH1A1,CYP24A1,GABRG3 |
| CSNK2A1 | kinase | 0.000327 | 3 | AHR,TP53,VIM |
| arsenite | chemical toxicant | 0.0315 | 3 | AHR,TP53,TXNRD1 |
| plicamycin | chemical drug | 0.0282 | 3 | AHR,SNAP25,TP53 |
| TGFBR1 | kinase | 0.0351 | 3 | AHR,PRKCB,VIM |
| MTORC1 | complex | 0.00423 | 3 | AHR,PPARGC1A,SLC2A4 |
| estradiol benzoate | chemical drug | 0.0123 | 3 | AHR,PPARD,SLC2A4 |
| LRP5 | transmembrane receptor | 0.00323 | 3 | AHR,ALDH1A1,SP7 |
| TGFB2 | growth factor | 0.0322 | 3 | AHR,ALDH1A1,CYP24A1 |
| semaxinib | chemical drug | 0.0495 | 3 | AHR,AHSG,EPHX2 |
| SOX9 | transcription regulator | 0.0404 | 3 | ADAMTS10,ALDH1A1,SP7 |
| mir-34 | microRNA | 0.0221 | 3 | ACTR1A,TP53,TUBB3 |
| RTN4 | other | 0.00733 | 3 | ACTR1A,MAP2,YWHAQ |
| QKI | other | 0.0115 | 3 | ACTN2,PPARGC1A,VIM |
| SLC27A2 | transporter | 0.00209 | 3 | ACOT8,ALDH1A1,ETFDH |
| PP2/AG1879 tyrosine kinase inhibitor | chemical drug | 0.014 | 3 | ACE,TP53,VIM |
| sodium bisulfide | chemical reagent | 0.00515 | 3 | ACE,TP53,TXNRD1 |
| azoxymethane | chemical toxicant | 0.0131 | 3 | ACE,PRKCB,TP53 |
| GIP | other | 0.0054 | 3 | ACE,PCK1,PPARGC1A |
| calcimycin | chemical reagent | 0.0322 | 3 | ACE,CASR,TP53 |
| bisindolylmaleimide I | chemical drug | 0.0295 | 3 | ACE,APOA1,Mt1 |
| RNF31 | enzyme | 0.00963 | 3 | ACE,APOA1,IL15RA |
| CYP19A1 | enzyme | 0.0256 | 3 | ACADVL,SLC2A4,TP53 |
| PLIN5 | other | 0.00182 | 3 | ACADVL,PPARGC1A,TXNRD1 |
| KLF15 | transcription regulator | 0.00445 | 3 | ACADVL,PPARGC1A,SLC2A4 |
| PCK1 | kinase | 8.71E-05 | 3 | ACADVL,PCK1,SLC2A4 |
| alisertib | chemical drug | 5.52E-05 | 3 | ACADVL,ETFDH,PPARGC1A |
| eicosapentenoic acid | chemical drug | 0.0221 | 3 | ACADVL,APOE,TP53 |
| LPIN1 | phosphatase | 0.000728 | 3 | ACADVL,APOE,PPARGC1A |
| clofibrate | chemical drug | 0.021 | 3 | ACADVL,ACOT8,Mt1 |
| BAG1 | other | 0.0256 | 2 | VCL,VIM |
| MRPS18B | other | 0.000713 | 2 | TUBB3,VIM |
| DUOXA1 | other | 0.00151 | 2 | TUBB3,VIM |
| HDAC6 | transcription regulator | 0.0315 | 2 | TPH2,VIM |
| vincristine | chemical drug | 0.0315 | 2 | TP53,YWHAQ |
| MEG3 | other | 0.00301 | 2 | TP53,VIM |
| PKNOX2 | transcription regulator | 0.00614 | 2 | TP53,VIM |
| TRIM37 | enzyme | 0.0212 | 2 | TP53,VIM |
| MTDH | transcription regulator | 0.0267 | 2 | TP53,VIM |
| KLF17 | transcription regulator | 0.00394 | 2 | TP53,VIM |
| FZD8 | G-protein coupled receptor | 0.00498 | 2 | TP53,VIM |
| WDR5 | transcription regulator | 0.00808 | 2 | TP53,VIM |
| CAT | enzyme | 0.0434 | 2 | TP53,VIM |
| MIR124 | group | 0.0191 | 2 | TP53,VIM |
| MDM4 | transcription regulator | 0.0144 | 2 | TP53,VIM |
| miR-1285-3p (and other miRNAs w/seed CUGGGCA) | mature microRNA | 0.000342 | 2 | TP53,VIM |
| HEXIM1 | transcription regulator | 0.0103 | 2 | TP53,VIM |
| SRSF1 | other | 0.0153 | 2 | TP53,VIM |
| CLCA2 | ion channel | 0.0022 | 2 | TP53,VIM |
| HSPA9 | other | 0.0244 | 2 | TP53,VIM |
| S100A4 | other | 0.0303 | 2 | TP53,VIM |
| IL32 | cytokine | 0.0448 | 2 | TP53,VIM |
| CUL7 | enzyme | 0.000948 | 2 | TP53,VIM |
| SDCBP | enzyme | 0.00951 | 2 | TP53,VIM |
| GFAP | other | 0.00614 | 2 | TP53,VIM |
| PCGF2 | transcription regulator | 0.0244 | 2 | TP53,VIM |
| NOX4 | enzyme | 0.042 | 2 | TP53,VIM |
| PARK7 | enzyme | 0.0172 | 2 | TP53,VIM |
| MTUS1 | other | 0.000512 | 2 | TP53,VIM |
| HDAC8 | transcription regulator | 0.00444 | 2 | TP53,VIM |
| methyl methanesulfonate | chemical toxicant | 0.0462 | 2 | TP53,VIM |
| carboplatin | chemical drug | 0.042 | 2 | TP53,VIM |
| calpeptin | chemical reagent | 0.00394 | 2 | TP53,VIM |
| mir-30 | microRNA | 0.0366 | 2 | TP53,VIM |
| CALR | transcription regulator | 0.034 | 2 | TP53,VCL |
| PTK2B | kinase | 0.00555 | 2 | TP53,VCL |
| MAFG | transcription regulator | 0.00951 | 2 | TP53,TXNRD1 |
| trichloroethylene | chemical toxicant | 0.0462 | 2 | SPARC,YWHAQ |
| benzene | chemical toxicant | 0.0462 | 2 | SPARC,YWHAQ |
| GLIS2 | transcription regulator | 0.0172 | 2 | SPARC,VIM |
| ABCB4 | transporter | 0.0379 | 2 | SPARC,VIM |
| miR-338-3p (miRNAs w/seed CCAGCAU) | mature microRNA | 0.0191 | 2 | SPARC,VIM |
| NVP-TAE684 | chemical drug | 0.0212 | 2 | SPARC,VIM |
| DNAJB6 | transcription regulator | 0.00444 | 2 | SPARC,VIM |
| CCR1 | G-protein coupled receptor | 0.0118 | 2 | SPARC,VIM |
| HTATIP2 | transcription regulator | 0.00808 | 2 | SPARC,TP53 |
| ITGB4 | transmembrane receptor | 0.00555 | 2 | SPARC,TP53 |
| zoledronic acid | chemical drug | 0.0315 | 2 | SPARC,TP53 |
| MSX2 | transcription regulator | 0.0256 | 2 | SP7,VIM |
| POSTN | other | 0.0279 | 2 | SP7,VIM |
| TSC22D3 | transcription regulator | 0.0406 | 2 | SP7,TP53 |
| EDIL3 | other | 0.000342 | 2 | SP7,TP53 |
| WWOX | enzyme | 0.00614 | 2 | SP7,TP53 |
| ABL1 | kinase | 0.0256 | 2 | SP7,TP53 |
| miR-128-3p (and other miRNAs w/seed CACAGUG) | mature microRNA | 0.00498 | 2 | SNAP25,TP53 |
| FA2H | enzyme | 0.00121 | 2 | SLC2A4,VIM |
| miR-223-3p (miRNAs w/seed GUCAGUU) | mature microRNA | 0.0127 | 2 | SLC2A4,VIM |
| tamsulosin | chemical drug | 0.000512 | 2 | SLC2A4,VIM |
| Pkg | group | 0.00808 | 2 | SLC2A4,VCL |
| CaMKII | complex | 0.00741 | 2 | SLC2A4,TP53 |
| CAST | peptidase | 0.00614 | 2 | SLC2A4,TP53 |
| RPS6KB1 | kinase | 0.0267 | 2 | SLC2A4,TP53 |
| AIFM1 | enzyme | 0.00346 | 2 | SLC2A4,TP53 |
| UCHL3 | peptidase | 0.00121 | 2 | SLC2A4,TP53 |
| 2,3-bis(4-hydroxyphenyl)-propionitrile | chemical reagent | 0.0406 | 2 | SLC2A4,TP53 |
| dichloroacetic acid | chemical drug | 0.00741 | 2 | SLC2A4,TP53 |
| pyruvaldehyde | chemical - endogenous mammalian | 0.0223 | 2 | SLC2A4,TP53 |
| sodium orthovanadate | chemical reagent | 0.0181 | 2 | SLC2A4,TP53 |
| zalcitabine | chemical drug | 0.00346 | 2 | SLC2A4,TP53 |
| XAV939 | chemical reagent | 0.0202 | 2 | SLC2A4,SP7 |
| SSTR2 | G-protein coupled receptor | 0.00951 | 2 | SLC18A2,VIM |
| amitriptyline | chemical drug | 0.00951 | 2 | SERPINA3,TP53 |
| RNF2 | transcription regulator | 0.042 | 2 | SERPINA1,TP53 |
| mir-145 | microRNA | 0.0353 | 2 | SDHA,TP53 |
| CHEK2 | kinase | 0.00498 | 2 | SDHA,TP53 |
| NCSTN | peptidase | 0.0191 | 2 | S100A1,VIM |
| mir-31 | microRNA | 0.0144 | 2 | RDX,SP7 |
| palbociclib | chemical drug | 0.0491 | 2 | PRKCB,TP53 |
| CDK6 | kinase | 0.00394 | 2 | PRKCB,TP53 |
| dehydrocostus lactone | chemical - endogenous non-mammalian | 0.0022 | 2 | PRKCB,TP53 |
| CP-55940 | chemical reagent | 0.0212 | 2 | PRKCB,RPL13A |
| 1,2-dimethylhydrazine | chemical toxicant | 0.0256 | 2 | PRDX6,TP53 |
| CDK9 | kinase | 0.042 | 2 | PPARGC1A,VIM |
| TWIST2 | transcription regulator | 0.034 | 2 | PPARGC1A,VIM |
| SERPINF1 | other | 0.0462 | 2 | PPARGC1A,VIM |
| PD 169316 | chemical drug | 0.00676 | 2 | PPARGC1A,VIM |
| naringenin | chemical - endogenous non-mammalian | 0.0256 | 2 | PPARGC1A,VIM |
| taurine | chemical - endogenous mammalian | 0.0223 | 2 | PPARGC1A,VIM |
| cyclic GMP | chemical - endogenous mammalian | 0.00394 | 2 | PPARGC1A,VCL |
| dasatinib | chemical drug | 0.0202 | 2 | PPARGC1A,TP53 |
| conjugated linoleic acid | chemical drug | 0.0291 | 2 | PPARGC1A,TP53 |
| MAPK13 | kinase | 0.00741 | 2 | PPARGC1A,TP53 |
| AURKA | kinase | 0.00808 | 2 | PPARGC1A,TP53 |
| CAMK4 | kinase | 0.0223 | 2 | PPARGC1A,TP53 |
| PIM1 | kinase | 0.0462 | 2 | PPARGC1A,TP53 |
| JARID2 | transcription regulator | 0.0103 | 2 | PPARGC1A,TP53 |
| DDIT4 | other | 0.0135 | 2 | PPARGC1A,TP53 |
| PIM2 | kinase | 0.00614 | 2 | PPARGC1A,TP53 |
| nitroprusside | chemical drug | 0.0392 | 2 | PPARGC1A,TP53 |
| CHIR 99021 | chemical drug | 0.0223 | 2 | PPARGC1A,SP7 |
| MEF2 | group | 0.0162 | 2 | PPARGC1A,SLC2A4 |
| Calcineurin A | group | 0.0172 | 2 | PPARGC1A,SLC2A4 |
| P2RY6 | G-protein coupled receptor | 0.011 | 2 | PPARGC1A,SLC2A4 |
| CFB | peptidase | 0.0162 | 2 | PPARGC1A,SLC2A4 |
| CCNC | other | 0.011 | 2 | PPARGC1A,SLC2A4 |
| MYBBP1A | transcription regulator | 0.000342 | 2 | PPARGC1A,SLC2A4 |
| mir-33 | microRNA | 0.00808 | 2 | PPARGC1A,SLC2A4 |
| H2AX | transcription regulator | 0.0327 | 2 | PPARGC1A,SLC2A4 |
| PKNOX1 | transcription regulator | 0.0118 | 2 | PPARGC1A,SLC2A4 |
| HSPB8 | kinase | 0.0127 | 2 | PPARGC1A,SLC2A4 |
| FKBP5 | enzyme | 0.0191 | 2 | PPARGC1A,SLC2A4 |
| CL 316243 | chemical drug | 0.0244 | 2 | PPARGC1A,SLC2A4 |
| stearic acid | chemical - endogenous mammalian | 0.0144 | 2 | PPARGC1A,SLC2A4 |
| agmatine | chemical - endogenous mammalian | 0.00741 | 2 | PPARGC1A,SLC22A4 |
| Cdk | group | 0.0162 | 2 | PPARGC1A,SDHA |
| MED30 | transcription regulator | 0.00614 | 2 | PPARGC1A,SDHA |
| digoxin | chemical drug | 0.00301 | 2 | PPARD,TP53 |
| MAP2K4 | kinase | 0.0462 | 2 | PPARD,TP53 |
| CHRNA7 | transmembrane receptor | 0.0181 | 2 | PPARD,TP53 |
| FABP4 | transporter | 0.0135 | 2 | PPARD,TP53 |
| MAP3K1 | kinase | 0.0406 | 2 | PPARD,TP53 |
| thioctic acid | chemical drug | 0.0491 | 2 | PPARD,TP53 |
| anandamide | chemical - endogenous mammalian | 0.0223 | 2 | PPARD,TP53 |
| telmisartan | chemical drug | 0.0476 | 2 | PPARD,SLC2A4 |
| natriuretic peptide derivative | biologic drug | 0.000104 | 2 | PPARD,PPARGC1A |
| SERTAD2 | transcription regulator | 0.00346 | 2 | PPARD,PPARGC1A |
| UBE3A | enzyme | 0.00951 | 2 | PML,TP53 |
| MECOM | transcription regulator | 0.0144 | 2 | PML,SPARC |
| galactose | chemical - endogenous mammalian | 0.00394 | 2 | PDP1,TP53 |
| SMAD6 | transcription regulator | 0.00444 | 2 | PCK1,SP7 |
| RBP4 | other | 0.00878 | 2 | PCK1,SLC2A4 |
| glucagon | biologic drug | 0.0462 | 2 | PCK1,SLC2A4 |
| 8-chlorophenylthio-adenosine 3',5'-cyclic monophosphate | chemical reagent | 0.0366 | 2 | PCK1,PPARGC1A |
| ZBTB20 | transcription regulator | 0.0448 | 2 | PCK1,PPARGC1A |
| CRTC2 | other | 0.0144 | 2 | PCK1,PPARGC1A |
| ELOVL5 | enzyme | 0.0118 | 2 | PCK1,PPARGC1A |
| Pka catalytic subunit | group | 0.011 | 2 | PCK1,PPARGC1A |
| RYR2 | ion channel | 0.0127 | 2 | PCK1,PPARGC1A |
| IRS2 | enzyme | 0.0279 | 2 | PCK1,PPARGC1A |
| lactic acid | chemical - endogenous mammalian | 0.0256 | 2 | PCK1,PPARGC1A |
| GC-GCR dimer | complex | 0.0135 | 2 | PCK1,PFKFB1 |
| SOX6 | transcription regulator | 0.00951 | 2 | MYL3,TP53 |
| PRKACA | kinase | 0.0303 | 2 | MYH10,PCK1 |
| HRH3 | G-protein coupled receptor | 0.0256 | 2 | Mup1 (includes others),SERPINA3 |
| PLA2G2A | enzyme | 0.00498 | 2 | Mup1 (includes others),PCK1 |
| Gm15807/Hmgn5 | other | 0.0153 | 2 | Mup1 (includes others),MYL3 |
| dimethylnitrosamine | chemical toxicant | 0.0327 | 2 | Mt1,VIM |
| diquat | chemical toxicant | 0.000713 | 2 | Mt1,TXNRD1 |
| dicumarol | chemical drug | 3.46E-05 | 2 | Mt1,TP53 |
| 3-methylcholanthrene | chemical toxicant | 0.042 | 2 | Mt1,TP53 |
| H-7 | chemical - kinase inhibitor | 0.0244 | 2 | Mt1,TP53 |
| iron | chemical - endogenous mammalian | 0.042 | 2 | Mt1,TF |
| STS | enzyme | 0.0144 | 2 | Mt1,SLC2A4 |
| CYB5R4 | enzyme | 0.00614 | 2 | Mt1,PPARGC1A |
| SRD5A1 | enzyme | 0.0103 | 2 | Mt1,PCK1 |
| mercuric chloride | chemical toxicant | 0.0191 | 2 | Mt1,Mup1 (includes others) |
| suramin | chemical drug | 0.0144 | 2 | MSTN,TP53 |
| miR-7a-5p (and other miRNAs w/seed GGAAGAC) | mature microRNA | 0.0135 | 2 | MAPKAP1,VIM |
| mir-7 | microRNA | 0.00951 | 2 | MAPKAP1,VIM |
| ROR1 | kinase | 0.0153 | 2 | MAP2,VIM |
| pubchem compound 135889696 | chemical reagent | 0.000512 | 2 | MAP2,TUBB3 |
| CHD8 | enzyme | 0.00394 | 2 | MAP2,TUBB3 |
| ZNF536 | transcription regulator | 0.000342 | 2 | MAP2,TUBB3 |
| CGP 42112 | chemical reagent | 0.00151 | 2 | MAP2,TUBB3 |
| CDK1 | kinase | 0.0118 | 2 | MAP2,TP53 |
| acteoside | chemical - endogenous non-mammalian | 0.0127 | 2 | MAP2,TP53 |
| paroxetine | chemical drug | 0.00878 | 2 | MAP2,TP53 |
| CDK2 | kinase | 0.0303 | 2 | MAP2,SNAP25 |
| NCL | other | 0.0118 | 2 | ILF3,TP53 |
| TNFAIP2 | other | 0.00394 | 2 | IL15RA,VIM |
| FKBP4 | enzyme | 0.00498 | 2 | IL15RA,PRDX6 |
| lapatinib/pazopanib | chemical drug | 0.00614 | 2 | IFNAR2,TP53 |
| HES3 | transcription regulator | 0.0267 | 2 | IFI30,IFNAR2 |
| K+ | chemical - endogenous mammalian | 0.0353 | 2 | GABRA5,TP53 |
| gamma-secretase inhibitor compound E | chemical reagent | 0.0135 | 2 | FKBP4,TUBB3 |
| WNT4 | cytokine | 0.0233 | 2 | FKBP4,SP7 |
| isoprenaline | chemical drug | 0.0315 | 2 | EPHX2,SLC2A4 |
| CARM1 | transcription regulator | 0.0223 | 2 | EPHX2,PCK1 |
| arsenic | chemical toxicant | 0.0135 | 2 | EPHX2,Mt1 |
| grape seed extract | chemical drug | 0.0153 | 2 | ENO1,VIM |
| SNHG11 | other | 0.00301 | 2 | ENO1,VIM |
| ENO1 | enzyme | 0.00151 | 2 | ENO1,VIM |
| PCCA-DT | other | 0.000206 | 2 | ENO1,VIM |
| NQO1 | enzyme | 0.0366 | 2 | ENO1,TP53 |
| peoniflorin | chemical drug | 0.00614 | 2 | ENO1,SNAP25 |
| MED13 | transcription regulator | 0.0315 | 2 | ENO1,SLC2A4 |
| 3,5-diiodothyronine | chemical - endogenous mammalian | 0.00184 | 2 | ENO1,SLC2A4 |
| T3-TR-RXR | complex | 0.0244 | 2 | ENO1,PCK1 |
| DMP1 | other | 0.0291 | 2 | ELAVL2,Mup1 (includes others) |
| MN1 | other | 0.000512 | 2 | CYP24A1,TP53 |
| calcifediol | chemical - endogenous mammalian | 0.011 | 2 | CYP24A1,TP53 |
| dichlororibofuranosylbenzimidazole | chemical toxicant | 0.00555 | 2 | CYP24A1,TP53 |
| PHEX | peptidase | 0.00259 | 2 | CYP24A1,SP7 |
| histone deacetylase | complex | 0.0233 | 2 | CYP24A1,PPARGC1A |
| SAFB | other | 0.042 | 2 | CNTNAP2,PAPSS2 |
| TSPYL5 | other | 0.00808 | 2 | CNTNAP2,DTD1 |
| 3-nitropropionic acid | chemical toxicant | 0.0406 | 2 | CNR1,TP53 |
| S-nitrosoglutathione | chemical toxicant | 0.0256 | 2 | CNR1,TP53 |
| immethridine | chemical reagent | 0.0103 | 2 | CNR1,SERPINA3 |
| Am 580 | chemical reagent | 0.0406 | 2 | CNR1,RAI14 |
| quinidine | chemical drug | 0.00151 | 2 | CHRM3,TP53 |
| GABBR1 | G-protein coupled receptor | 0.00394 | 2 | CASR,SP7 |
| sesaminol | chemical - endogenous non-mammalian | 0.0279 | 2 | CASQ1,MYL3 |
| miR-199a-3p (and other miRNAs w/seed CAGUAGU) | mature microRNA | 0.00741 | 2 | CALU,PRDX6 |
| SAMMSON | other | 0.0244 | 2 | ATP5F1B,TP53 |
| NFU1 | other | 0.0267 | 2 | ATP5F1B,SDHA |
| HDL | complex | 0.042 | 2 | ATP5F1B,PPARGC1A |
| carbonyl cyanide p-(trifluoromethoxy)phenylhydrazone | chemical reagent | 0.00394 | 2 | ATP5F1B,PPARGC1A |
| 2,4-dinitrophenol | chemical toxicant | 0.000948 | 2 | ATP5F1B,PPARGC1A |
| hexarelin | chemical toxicant | 0.00444 | 2 | ATP5F1B,PPARGC1A |
| CA9 | enzyme | 0.0181 | 2 | ATP5F1B,ENO1 |
| IND S1 | chemical - kinase inhibitor | 0.00741 | 2 | ATP5F1B,ENO1 |
| MEL T1 | chemical - kinase inhibitor | 0.00394 | 2 | ATP5F1B,ENO1 |
| UBQLN2 | other | 0.0379 | 2 | ATAD1,HPX |
| elovanoid N32 | chemical - endogenous mammalian | 0.00614 | 2 | APOE,TP53 |
| elovanoid N34 | chemical - endogenous mammalian | 0.00676 | 2 | APOE,TP53 |
| COL5A1 | other | 0.0191 | 2 | APOE,TF |
| sucrose | chemical - endogenous mammalian | 0.0223 | 2 | APOE,SNAP25 |
| CYP27A1 | enzyme | 0.00741 | 2 | APOE,SLC2A4 |
| baicalein | chemical drug | 0.0291 | 2 | APOE,PRDX6 |
| PLIN2 | other | 0.00151 | 2 | APOE,PCK1 |
| MBD1 | transcription regulator | 0.00808 | 2 | APOE,EFNA5 |
| NR2C2 | ligand-dependent nuclear receptor | 0.0244 | 2 | APOE,CYP24A1 |
| carbamylcholine | chemical drug | 0.0256 | 2 | APOE,CHRM3 |
| selenium | chemical drug | 0.0153 | 2 | APOA1,TXNRD1 |
| TXN | enzyme | 0.034 | 2 | APOA1,TP53 |
| glycyrrhizic acid | chemical drug | 0.0144 | 2 | APOA1,TP53 |
| lipid | chemical - endogenous mammalian | 0.0267 | 2 | APOA1,SLC2A4 |
| salicylic acid | chemical drug | 0.042 | 2 | APOA1,SLC2A4 |
| beta-carotene | chemical - endogenous mammalian | 0.0291 | 2 | APOA1,SLC2A4 |
| phorbol 12,13-dibutyrate | chemical - endogenous non-mammalian | 0.0191 | 2 | APOA1,PRKCB |
| SAR1B | enzyme | 0.00394 | 2 | APOA1,PPARGC1A |
| L 663536 | chemical reagent | 0.0103 | 2 | APOA1,PPARD |
| obeticholic acid | chemical drug | 0.034 | 2 | APOA1,PCK1 |
| NR2F1 | ligand-dependent nuclear receptor | 0.0353 | 2 | APOA1,PCK1 |
| Ppar | group | 0.0153 | 2 | APOA1,MSTN |
| H2AZ1 | other | 0.0392 | 2 | APOA1,GALNTL6 |
| 20-hydroxyeicosatetraenoic acid | chemical - endogenous mammalian | 0.000512 | 2 | APOA1,EPHX2 |
| SCARB1 | transporter | 0.034 | 2 | APOA1,APOE |
| XRCC6 | enzyme | 0.00498 | 2 | APOA1,APOE |
| CETP | enzyme | 0.00808 | 2 | APOA1,APOE |
| cetrorelix | biologic drug | 0.00301 | 2 | ANXA5,TP53 |
| zVAD-FMK | chemical - protease inhibitor | 0.0353 | 2 | ANXA5,TP53 |
| SOX2-OT | other | 0.0022 | 2 | ALDH1A1,VIM |
| Sox2ot | other | 0.00498 | 2 | ALDH1A1,VIM |
| FZD4 | G-protein coupled receptor | 0.00151 | 2 | ALDH1A1,VIM |
| FTO | enzyme | 0.0191 | 2 | ALDH1A1,TP53 |
| ALDH1A2 | enzyme | 0.0153 | 2 | ALDH1A1,SPARC |
| PXR ligand-PXR-Retinoic acid-RXRα | complex | 0.0267 | 2 | ALDH1A1,PAPSS2 |
| myclobutanil | chemical toxicant | 0.0267 | 2 | ALDH1A1,GABRG3 |
| HOXA13 | transcription regulator | 0.00878 | 2 | ALDH1A1,ENO1 |
| IL20 | cytokine | 0.0202 | 2 | AHSG,APOA1 |
| LGR4 | transmembrane receptor | 0.034 | 2 | AHR,VIM |
| AIP | transcription regulator | 0.0434 | 2 | AHR,TP53 |
| PPP5C | phosphatase | 0.00259 | 2 | AHR,TP53 |
| PD 150606 | chemical - protease inhibitor | 0.000713 | 2 | AHR,TP53 |
| 9,10-dimethyl-1,2-benzanthracene | chemical toxicant | 0.0462 | 2 | AHR,TP53 |
| MDL 28170 | chemical toxicant | 0.000948 | 2 | AHR,TP53 |
| IL23A | cytokine | 0.0233 | 2 | AHR,HPX |
| PRKDC | kinase | 0.0127 | 2 | AHR,APOE |
| GLIS1 | transcription regulator | 0.0191 | 2 | ADAMTS10,SPARC |
| SORT1 | G-protein coupled receptor | 0.0022 | 2 | ACTN2,SLC2A4 |
| CDON | other | 0.0022 | 2 | ACE,TUBB3 |
| PROC | peptidase | 0.0327 | 2 | ACE,TP53 |
| MAP3K5 | kinase | 0.00741 | 2 | ACE,TP53 |
| oltipraz | chemical drug | 0.0144 | 2 | ACE,TP53 |
| baicalin | chemical - endogenous non-mammalian | 0.034 | 2 | ACE,SPARC |
| NPFF | other | 0.00301 | 2 | ACE,SLC2A4 |
| eugenol | chemical - endogenous non-mammalian | 0.000713 | 2 | ACE,SLC2A4 |
| 27-hydroxycholesterol | chemical - endogenous mammalian | 0.00614 | 2 | ACE,SLC2A4 |
| propranolol | chemical drug | 0.0256 | 2 | ACE,PPARGC1A |
| Nppb | other | 0.00951 | 2 | ACE,MYL3 |
| CUL3 | enzyme | 0.0448 | 2 | ACE,MYL3 |
| ACE | peptidase | 0.0118 | 2 | ACE,ATP5F1B |
| 24-hydroxycholesterol | chemical - endogenous mammalian | 0.00346 | 2 | ACE,APOE |
| NOSTRIN | transcription regulator | 0.0153 | 2 | ACE,ANXA5 |
| WNT10B | other | 0.00184 | 2 | ACADVL,SP7 |
| OMA1 | peptidase | 0.00498 | 2 | ACADVL,PPARGC1A |
| SOST | other | 0.0181 | 2 | ACADVL,PPARGC1A |
| ABCC8 | transporter | 0.00444 | 2 | ACADVL,PPARGC1A |
| KCNJ11 | ion channel | 0.00151 | 2 | ACADVL,PPARGC1A |
| alpha-keto-beta-methylvaleric acid | chemical - endogenous mammalian | 0.000713 | 2 | ACADVL,PPARGC1A |
| pyrrolidonecarboxylic acid | chemical - endogenous mammalian | 0.000713 | 2 | ACADVL,PPARGC1A |
| RYR1 | ion channel | 0.0181 | 2 | ACADVL,ETFDH |
| P5091 | chemical reagent | 0.0176 | 1 | VIM |
| MS049 | chemical reagent | 0.0118 | 1 | VIM |
| NR2F1-AS1 | other | 0.0234 | 1 | VIM |
| LINC00842 | other | 0.0349 | 1 | VIM |
| FOXQ1 | transcription regulator | 0.0349 | 1 | VIM |
| NSD3 | enzyme | 0.0349 | 1 | VIM |
| ST6GALNAC1 | enzyme | 0.0406 | 1 | VIM |
| ZNF652 | other | 0.0406 | 1 | VIM |
| HOXA-AS3 | other | 0.0234 | 1 | VIM |
| NSRP1 | other | 0.0292 | 1 | VIM |
| MIR9-1HG | other | 0.0406 | 1 | VIM |
| GLIPR2 | other | 0.0234 | 1 | VIM |
| BCAR4 | other | 0.0406 | 1 | VIM |
| CHD5 | enzyme | 0.0292 | 1 | VIM |
| SPRED2 | cytokine | 0.0176 | 1 | VIM |
| LINC00887 | other | 0.0234 | 1 | VIM |
| NBR2 | other | 0.0234 | 1 | VIM |
| ZDHHC2 | enzyme | 0.0349 | 1 | VIM |
| RHBDF1 | other | 0.0118 | 1 | VIM |
| RADIL | other | 0.0176 | 1 | VIM |
| PTPN23 | phosphatase | 0.0176 | 1 | VIM |
| MAP3K21 | kinase | 0.0463 | 1 | VIM |
| lumefantrine | chemical drug | 0.0349 | 1 | VIM |
| PDLIM1 | transcription regulator | 0.0234 | 1 | VIM |
| hydrogel | chemical drug | 0.0463 | 1 | VIM |
| PF-4691502 | chemical drug | 0.0234 | 1 | VIM |
| FAF1 | other | 0.0406 | 1 | VIM |
| CLDN1 | other | 0.0349 | 1 | VIM |
| CBR3-AS1 | other | 0.0406 | 1 | VIM |
| FGD5-AS1 | other | 0.0234 | 1 | VIM |
| PLCD1 | enzyme | 0.0463 | 1 | VIM |
| mir-190 | microRNA | 0.0463 | 1 | VIM |
| miR-382-5p (miRNAs w/seed AAGUUGU) | mature microRNA | 0.0292 | 1 | VIM |
| miR-508-3p (miRNAs w/seed GAUUGUA) | mature microRNA | 0.0292 | 1 | VIM |
| miR-320b (and other miRNAs w/seed AAAGCUG) | mature microRNA | 0.0463 | 1 | VIM |
| mir-612 | microRNA | 0.0292 | 1 | VIM |
| mir-138 | microRNA | 0.0234 | 1 | VIM |
| miR-142-5p (and other miRNAs w/seed AUAAAGU) | mature microRNA | 0.0234 | 1 | VIM |
| mir-944 | microRNA | 0.0118 | 1 | VIM |
| STRN | other | 0.0463 | 1 | VIM |
| APAF1 | other | 0.0349 | 1 | VIM |
| MIR5590 | microRNA | 0.0118 | 1 | VIM |
| EEF1G | translation regulator | 0.00591 | 1 | VIM |
| galunisertib | chemical drug | 0.0349 | 1 | VIM |
| EFEMP1 | enzyme | 0.0463 | 1 | VIM |
| DLGAP1 | other | 0.0292 | 1 | VIM |
| NME3 | kinase | 0.0292 | 1 | VIM |
| TIAM2 | enzyme | 0.0118 | 1 | VIM |
| LASP1 | transporter | 0.0463 | 1 | VIM |
| SLC1A5 | transporter | 0.0234 | 1 | VIM |
| SLC12A6 | transporter | 0.0176 | 1 | VIM |
| MAD2L2 | enzyme | 0.0463 | 1 | VIM |
| DKC1 | enzyme | 0.0463 | 1 | VIM |
| LDN-193189 | chemical drug | 0.0234 | 1 | VIM |
| RAMP3 | G-protein coupled receptor | 0.0118 | 1 | VIM |
| PPID | enzyme | 0.0349 | 1 | VIM |
| EEF2K | kinase | 0.0349 | 1 | VIM |
| BLACAT1 | other | 0.0463 | 1 | VIM |
| PROKR1 | G-protein coupled receptor | 0.0463 | 1 | VIM |
| RAB7A | enzyme | 0.0349 | 1 | VIM |
| TRMT13 | other | 0.0176 | 1 | VIM |
| RAB11FIP1 | other | 0.0176 | 1 | VIM |
| ACTA2 | other | 0.0463 | 1 | VIM |
| SRGN | other | 0.0349 | 1 | VIM |
| PODXL | kinase | 0.0234 | 1 | VIM |
| RPS4Y1 | other | 0.0118 | 1 | VIM |
| capivasertib | chemical drug | 0.0234 | 1 | VIM |
| DPYSL2 | enzyme | 0.0234 | 1 | VIM |
| RPL9 | other | 0.00591 | 1 | VIM |
| crenigacestat | chemical drug | 0.0118 | 1 | VIM |
| butylidenephthalide | chemical reagent | 0.0406 | 1 | VIM |
| 4-aminophenol | chemical - endogenous mammalian | 0.0349 | 1 | VIM |
| miR-508-3p inhibitor | chemical reagent | 0.0406 | 1 | VIM |
| alpha-hydroxyglutarate | chemical - endogenous mammalian | 0.0349 | 1 | VIM |
| trehalose | chemical - endogenous mammalian | 0.0349 | 1 | VIM |
| PXN | other | 0.0406 | 1 | VCL |
| PIWIL1 | enzyme | 0.0349 | 1 | VCL |
| PHIP | other | 0.0406 | 1 | VCL |
| CTNNA1 | other | 0.00591 | 1 | VCL |
| Macf1 | other | 0.0234 | 1 | VCL |
| PTP4A2 | phosphatase | 0.0234 | 1 | VCL |
| adenosine dialdehyde | chemical reagent | 0.0349 | 1 | TXNRD1 |
| SLC6A12 | transporter | 0.0176 | 1 | TUBB3 |
| SOX12 | transcription regulator | 0.0234 | 1 | TUBB3 |
| CHKA | kinase | 0.0292 | 1 | TUBB3 |
| DCX | other | 0.0292 | 1 | TUBB3 |
| ELAVL3 | other | 0.0176 | 1 | TUBB3 |
| dinophysistoxin 1 | chemical toxicant | 0.0118 | 1 | TUBB3 |
| octanal | chemical - endogenous mammalian | 0.0118 | 1 | TUBB3 |
| SLC6A2 | transporter | 0.0406 | 1 | TPH2 |
| citalopram | chemical drug | 0.0463 | 1 | TPH2 |
| GY1-22 | chemical reagent | 0.0176 | 1 | TP53 |
| CGP 74514A | chemical drug | 0.0234 | 1 | TP53 |
| ganciclovir | chemical drug | 0.0349 | 1 | TP53 |
| teniposide | chemical drug | 0.0292 | 1 | TP53 |
| nitrite | chemical - endogenous mammalian | 0.0349 | 1 | TP53 |
| TRE-TTC2-1/2 | group | 0.0118 | 1 | TP53 |
| extremely high molecular weight hyaluronic acid | chemical reagent | 0.0176 | 1 | TP53 |
| TPPP2 | other | 0.0292 | 1 | TP53 |
| (diaminocyclohexane)(diacetato)(dichloro)platinum | chemical reagent | 0.0176 | 1 | TP53 |
| 2,5-bis(5-hydroxymethyl-2-thienyl)furan | chemical reagent | 0.0463 | 1 | TP53 |
| (6)-gingerol | chemical - endogenous non-mammalian | 0.0406 | 1 | TP53 |
| dibutylnitrosamine | chemical toxicant | 0.0118 | 1 | TP53 |
| tris(2,3-dibromopropyl)phosphate | chemical toxicant | 0.00591 | 1 | TP53 |
| asparaginase | group | 0.0406 | 1 | TP53 |
| SNHG1 | other | 0.0234 | 1 | TP53 |
| tozasertib | chemical drug | 0.0234 | 1 | TP53 |
| citrinin | chemical toxicant | 0.0463 | 1 | TP53 |
| diosmin | chemical drug | 0.0176 | 1 | TP53 |
| omethoate | chemical toxicant | 0.0118 | 1 | TP53 |
| nitrosomethylurethane | chemical reagent | 0.0118 | 1 | TP53 |
| piracetam | chemical drug | 0.0118 | 1 | TP53 |
| silipide | chemical drug | 0.0118 | 1 | TP53 |
| givinostat | chemical drug | 0.0349 | 1 | TP53 |
| CYGB | transporter | 0.0349 | 1 | TP53 |
| SLC39A9 | transporter | 0.0349 | 1 | TP53 |
| NSA2 | other | 0.0118 | 1 | TP53 |
| THG1L | enzyme | 0.00591 | 1 | TP53 |
| MELK | kinase | 0.0292 | 1 | TP53 |
| RFFL | enzyme | 0.0176 | 1 | TP53 |
| DDX53 | other | 0.00591 | 1 | TP53 |
| ETHE1 | enzyme | 0.0292 | 1 | TP53 |
| TTC5 | other | 0.0176 | 1 | TP53 |
| USP28 | peptidase | 0.0406 | 1 | TP53 |
| KMT2E | enzyme | 0.0349 | 1 | TP53 |
| RIOK1 | kinase | 0.00591 | 1 | TP53 |
| BCAS2 | other | 0.0406 | 1 | TP53 |
| BIRC6 | enzyme | 0.0176 | 1 | TP53 |
| OR51E1 | G-protein coupled receptor | 0.0176 | 1 | TP53 |
| RASSF3 | other | 0.0349 | 1 | TP53 |
| SLC5A8 | transporter | 0.0406 | 1 | TP53 |
| EEF1E1 | translation regulator | 0.00591 | 1 | TP53 |
| MDC1 | other | 0.0118 | 1 | TP53 |
| LINC00337 | other | 0.0292 | 1 | TP53 |
| PYHIN1 | other | 0.0292 | 1 | TP53 |
| DDIAS | other | 0.00591 | 1 | TP53 |
| ADNP | transcription regulator | 0.0118 | 1 | TP53 |
| CDKN2AIP | transcription regulator | 0.00591 | 1 | TP53 |
| Ribosomal Subunit 80S | complex | 0.0176 | 1 | TP53 |
| TRIM39 | other | 0.00591 | 1 | TP53 |
| REV3L | enzyme | 0.00591 | 1 | TP53 |
| CTBP1-DT | other | 0.0176 | 1 | TP53 |
| SNRPA1 | other | 0.0118 | 1 | TP53 |
| antisense oligonucleotide | biologic drug | 0.0463 | 1 | TP53 |
| TRIM67 | other | 0.0176 | 1 | TP53 |
| K Channel | complex | 0.0234 | 1 | TP53 |
| Importin alpha | group | 0.0118 | 1 | TP53 |
| xenon | chemical drug | 0.0176 | 1 | TP53 |
| Magea3 (includes others) | other | 0.00591 | 1 | TP53 |
| CLK | group | 0.0118 | 1 | TP53 |
| TAF9 | transcription regulator | 0.0292 | 1 | TP53 |
| PCM1 | other | 0.0406 | 1 | TP53 |
| pertuzumab | biologic drug | 0.0234 | 1 | TP53 |
| Pik3r | group | 0.0349 | 1 | TP53 |
| JDP2 | transcription regulator | 0.0406 | 1 | TP53 |
| UNG | enzyme | 0.0234 | 1 | TP53 |
| ZMAT3 | other | 0.0176 | 1 | TP53 |
| YEATS4 | transcription regulator | 0.0176 | 1 | TP53 |
| nomilin | chemical - endogenous non-mammalian | 0.0349 | 1 | TP53 |
| TERF1 | other | 0.0176 | 1 | TP53 |
| PF-4929113 | chemical drug | 0.0118 | 1 | TP53 |
| T-2 toxin | chemical toxicant | 0.0349 | 1 | TP53 |
| TLCD3A | other | 0.0234 | 1 | TP53 |
| SEMA3F | other | 0.0292 | 1 | TP53 |
| DAPK1 | kinase | 0.0292 | 1 | TP53 |
| SNCAIP | transcription regulator | 0.0118 | 1 | TP53 |
| PSMC3 | enzyme | 0.0176 | 1 | TP53 |
| NFE2L3 | transcription regulator | 0.0463 | 1 | TP53 |
| RAD23A | other | 0.00591 | 1 | TP53 |
| CDC25B | phosphatase | 0.0234 | 1 | TP53 |
| XPC | transcription regulator | 0.0349 | 1 | TP53 |
| HINT1 | enzyme | 0.0463 | 1 | TP53 |
| DUSP10 | phosphatase | 0.0406 | 1 | TP53 |
| GLO1 | enzyme | 0.0406 | 1 | TP53 |
| GRK5 | kinase | 0.0463 | 1 | TP53 |
| RPL23 | other | 0.00591 | 1 | TP53 |
| RGS6 | enzyme | 0.0176 | 1 | TP53 |
| MGMT | enzyme | 0.0176 | 1 | TP53 |
| RPL24 | other | 0.00591 | 1 | TP53 |
| RALB | enzyme | 0.0349 | 1 | TP53 |
| NEK10 | kinase | 0.0349 | 1 | TP53 |
| AGAP2 | enzyme | 0.0176 | 1 | TP53 |
| MCRS1 | other | 0.0176 | 1 | TP53 |
| SCGN | other | 0.0118 | 1 | TP53 |
| TYMS | enzyme | 0.0292 | 1 | TP53 |
| RNF34 | enzyme | 0.0292 | 1 | TP53 |
| SNCB | other | 0.0406 | 1 | TP53 |
| RAD51 | enzyme | 0.0176 | 1 | TP53 |
| miR-542-3p (miRNAs w/seed GUGACAG) | mature microRNA | 0.0406 | 1 | TP53 |
| mir-504 | microRNA | 0.0118 | 1 | TP53 |
| miR-758-5p (and other miRNAs w/seed GGUUGAC) | mature microRNA | 0.0118 | 1 | TP53 |
| miR-1204 (miRNAs w/seed CGUGGCC) | mature microRNA | 0.00591 | 1 | TP53 |
| SMARCA1 | transcription regulator | 0.0463 | 1 | TP53 |
| PDE3A | enzyme | 0.0406 | 1 | TP53 |
| EIF5A | translation regulator | 0.0406 | 1 | TP53 |
| RPS14 | translation regulator | 0.0176 | 1 | TP53 |
| CCNG1 | other | 0.0118 | 1 | TP53 |
| KCND2 | ion channel | 0.0292 | 1 | TP53 |
| ARHGAP1 | other | 0.0176 | 1 | TP53 |
| CALCR | G-protein coupled receptor | 0.0234 | 1 | TP53 |
| TOM1L1 | other | 0.00591 | 1 | TP53 |
| TSC22D1 | transcription regulator | 0.0406 | 1 | TP53 |
| TOP3A | enzyme | 0.0234 | 1 | TP53 |
| STIP1 | other | 0.0176 | 1 | TP53 |
| CDC7 | kinase | 0.0118 | 1 | TP53 |
| MAGEA4 | other | 0.0118 | 1 | TP53 |
| SPRR2B | other | 0.00591 | 1 | TP53 |
| MSLN | other | 0.0349 | 1 | TP53 |
| PPM1B | phosphatase | 0.0463 | 1 | TP53 |
| KSR1 | kinase | 0.0463 | 1 | TP53 |
| UBE4B | enzyme | 0.0118 | 1 | TP53 |
| TSPYL2 | other | 0.0118 | 1 | TP53 |
| PEBP4 | other | 0.0234 | 1 | TP53 |
| TP53BP1 | transcription regulator | 0.0118 | 1 | TP53 |
| CNOT8 | transcription regulator | 0.00591 | 1 | TP53 |
| USP11 | peptidase | 0.0463 | 1 | TP53 |
| BRCA2 | transcription regulator | 0.0292 | 1 | TP53 |
| SPN | transmembrane receptor | 0.0463 | 1 | TP53 |
| RBM38 | other | 0.0463 | 1 | TP53 |
| RPS6 | other | 0.0234 | 1 | TP53 |
| SDC2 | other | 0.0463 | 1 | TP53 |
| CEL | enzyme | 0.0292 | 1 | TP53 |
| BCLAF1 | transcription regulator | 0.0118 | 1 | TP53 |
| GNL1 | other | 0.0406 | 1 | TP53 |
| DNAJA1 | other | 0.0349 | 1 | TP53 |
| CELF2 | other | 0.0234 | 1 | TP53 |
| CUL1 | enzyme | 0.0463 | 1 | TP53 |
| NINJ1 | other | 0.00591 | 1 | TP53 |
| PDZD2 | other | 0.0176 | 1 | TP53 |
| DDX19B | enzyme | 0.0234 | 1 | TP53 |
| DMTF1 | transcription regulator | 0.0406 | 1 | TP53 |
| APLP2 | other | 0.0234 | 1 | TP53 |
| MVK | kinase | 0.0118 | 1 | TP53 |
| TOP2B | enzyme | 0.0234 | 1 | TP53 |
| BRAT1 | other | 0.00591 | 1 | TP53 |
| SVIL | other | 0.0118 | 1 | TP53 |
| CCNL2 | other | 0.0176 | 1 | TP53 |
| BLM | enzyme | 0.0463 | 1 | TP53 |
| CENPJ | transcription regulator | 0.0176 | 1 | TP53 |
| RCHY1 | enzyme | 0.0292 | 1 | TP53 |
| RPS7 | other | 0.0176 | 1 | TP53 |
| TERF2 | transcription regulator | 0.0463 | 1 | TP53 |
| RAPGEF4 | other | 0.0406 | 1 | TP53 |
| WT1-AS | other | 0.00591 | 1 | TP53 |
| RPS25 | other | 0.0118 | 1 | TP53 |
| TAGLN | other | 0.0118 | 1 | TP53 |
| PCBP4 | other | 0.0118 | 1 | TP53 |
| PLAC8 | other | 0.0406 | 1 | TP53 |
| CREG1 | transcription regulator | 0.0349 | 1 | TP53 |
| WEE1 | kinase | 0.00591 | 1 | TP53 |
| BARD1 | transcription regulator | 0.0349 | 1 | TP53 |
| GNL2 | enzyme | 0.0292 | 1 | TP53 |
| brequinar | chemical drug | 0.0234 | 1 | TP53 |
| DAPK2 | kinase | 0.00591 | 1 | TP53 |
| DDT | enzyme | 0.0292 | 1 | TP53 |
| BUB3 | other | 0.0176 | 1 | TP53 |
| DDX20 | transcription regulator | 0.0118 | 1 | TP53 |
| glyceraldehyde-BSA | chemical reagent | 0.0406 | 1 | TP53 |
| methylglyoxal-BSA | chemical reagent | 0.0406 | 1 | TP53 |
| CDC6 | other | 0.0176 | 1 | TP53 |
| RPS27 | other | 0.00591 | 1 | TP53 |
| Zfp871 | other | 0.00591 | 1 | TP53 |
| RPS27L | translation regulator | 0.00591 | 1 | TP53 |
| Ifna4 | other | 0.0349 | 1 | TP53 |
| Nedd4 | enzyme | 0.0118 | 1 | TP53 |
| SPRR3 | other | 0.0234 | 1 | TP53 |
| APH1B | peptidase | 0.0118 | 1 | TP53 |
| RPL5 | other | 0.0234 | 1 | TP53 |
| (+)-R-3-(4-chlorophenyl)-3-(1-hydroxymethylcyclopropylmethoxy)-2-(4-nitrobenzyl)-2,3-dihydroisoindol-1-one | chemical reagent | 0.0176 | 1 | TP53 |
| VNN1 | enzyme | 0.0118 | 1 | TP53 |
| MI-63 | chemical reagent | 0.0118 | 1 | TP53 |
| DUOX2 | enzyme | 0.0176 | 1 | TP53 |
| GTSE1 | other | 0.00591 | 1 | TP53 |
| PSENEN | peptidase | 0.0406 | 1 | TP53 |
| RPL7A | other | 0.00591 | 1 | TP53 |
| JV-1-65 hGHRH(1-29)NH2 | chemical reagent | 0.0292 | 1 | TP53 |
| JV-1-63 hGHRH(1-29)NH2 | chemical reagent | 0.00591 | 1 | TP53 |
| MI-773 | chemical drug | 0.0292 | 1 | TP53 |
| azurin 50-77 | biologic drug | 0.0234 | 1 | TP53 |
| 5-fluorouracil/thymidine | chemical drug | 0.00591 | 1 | TP53 |
| 2-ethylestrone-3-O-sulfamate | chemical reagent | 0.00591 | 1 | TP53 |
| O-(chloroacetylcarbamoyl)fumagillol | chemical drug | 0.0176 | 1 | TP53 |
| dacarbazine | chemical drug | 0.0406 | 1 | TP53 |
| meclofenamic acid | chemical drug | 0.0118 | 1 | TP53 |
| EGTA acetoxymethyl ester | chemical reagent | 0.0349 | 1 | TP53 |
| mafosfamide | chemical drug | 0.0234 | 1 | TP53 |
| tirapazamine | chemical drug | 0.0234 | 1 | TP53 |
| FK 409 | chemical drug | 0.0176 | 1 | TP53 |
| plevitrexed | chemical drug | 0.0292 | 1 | TP53 |
| zeocin | chemical reagent | 0.0118 | 1 | TP53 |
| 2-aminofluorene | chemical toxicant | 0.00591 | 1 | TP53 |
| bilobalide | chemical - endogenous non-mammalian | 0.0234 | 1 | TP53 |
| ABT 702 | chemical drug | 0.00591 | 1 | TP53 |
| NU 1025 | chemical toxicant | 0.0176 | 1 | TP53 |
| talipexole | chemical drug | 0.0118 | 1 | TP53 |
| vandetanib | chemical drug | 0.0463 | 1 | TP53 |
| cidofovir | chemical drug | 0.0118 | 1 | TP53 |
| propafenone | chemical drug | 0.0234 | 1 | TP53 |
| AN-238 | chemical toxicant | 0.00591 | 1 | TP53 |
| lasofoxifene | chemical drug | 0.00591 | 1 | TP53 |
| hydroxyacetylaminofluorene | chemical toxicant | 0.00591 | 1 | TP53 |
| fluoranthene | chemical toxicant | 0.00591 | 1 | TP53 |
| carnosol | chemical - endogenous non-mammalian | 0.0463 | 1 | TP53 |
| adozelesin | chemical drug | 0.00591 | 1 | TP53 |
| 10-decarbamoylmitomycin C | chemical toxicant | 0.0176 | 1 | TP53 |
| propyl gallate | chemical toxicant | 0.0349 | 1 | TP53 |
| NPM1-RARA | fusion gene/product | 0.0118 | 1 | TP53 |
| tilimycin | chemical - endogenous non-mammalian | 0.00591 | 1 | TP53 |
| TFMB-(R)-2-HG | chemical reagent | 0.00591 | 1 | TP53 |
| FT671 | chemical - protease inhibitor | 0.0292 | 1 | TP53 |
| HL001 | chemical reagent | 0.0118 | 1 | TP53 |
| 1-Naphthalenesulfonyl-IW-CHO | chemical - protease inhibitor | 0.0292 | 1 | TP53 |
| ATSP-7041 | chemical reagent | 0.00591 | 1 | TP53 |
| calpain inhibitor 2 | chemical - protease inhibitor | 0.0406 | 1 | TP53 |
| Z-Leu-Leu-Leu-B(OH)2 | chemical reagent | 0.0349 | 1 | TP53 |
| MW167 | chemical - protease inhibitor | 0.0118 | 1 | TP53 |
| NRP2-PLXNA1 | complex | 0.00591 | 1 | TP53 |
| dihematoporphyrin ether | chemical drug | 0.0463 | 1 | TP53 |
| ampelopsin | chemical drug | 0.0292 | 1 | TP53 |
| vinflunine | chemical drug | 0.0118 | 1 | TP53 |
| pergolide | chemical drug | 0.0292 | 1 | TP53 |
| metal ion | chemical - other | 0.00591 | 1 | TP53 |
| ISA-2011B | chemical drug | 0.00591 | 1 | TP53 |
| AVI-4126 | biologic drug | 0.0176 | 1 | TP53 |
| NADPH | chemical - endogenous mammalian | 0.0292 | 1 | TP53 |
| floxuridine | chemical drug | 0.0292 | 1 | TP53 |
| sparfosic acid | chemical drug | 0.0349 | 1 | TP53 |
| pemetrexed | chemical drug | 0.0292 | 1 | TP53 |
| CM101 | chemical drug | 0.0234 | 1 | TP53 |
| AZD1480 | chemical drug | 0.0292 | 1 | TF |
| CDR2 | other | 0.0292 | 1 | TF |
| 8-methyl-pyridoxatin | chemical reagent | 0.0118 | 1 | TF |
| ciclopirox olamine | chemical drug | 0.0463 | 1 | TF |
| NRXN2 | transporter | 0.00591 | 1 | TAFA1 |
| Nrxn3 | other | 0.00591 | 1 | TAFA1 |
| Ho | group | 0.0292 | 1 | SPARC |
| VIPAS39 | other | 0.0406 | 1 | SPARC |
| 2,3-bis(3'-hydroxybenzyl)butane-1,4-diol | chemical - endogenous mammalian | 0.0292 | 1 | SPARC |
| MIA | other | 0.0463 | 1 | SPARC |
| TP53INP1 | other | 0.0406 | 1 | SPARC |
| CXXC4 | other | 0.0349 | 1 | SPARC |
| synthetic peptide | chemical reagent | 0.0118 | 1 | SPARC |
| CBFB-MYH11 | fusion gene/product | 0.0292 | 1 | SPARC |
| advanced glycation end product 3 | chemical - endogenous mammalian | 0.0292 | 1 | SP7 |
| DIPQUO | chemical reagent | 0.0292 | 1 | SP7 |
| AMELY | growth factor | 0.0349 | 1 | SP7 |
| RIOX1 | enzyme | 0.0406 | 1 | SP7 |
| PDLIM7 | other | 0.0463 | 1 | SP7 |
| EZH1 | enzyme | 0.0406 | 1 | SP7 |
| miR-637 (and other miRNAs w/seed CUGGGGG) | mature microRNA | 0.0118 | 1 | SP7 |
| OGN | growth factor | 0.0406 | 1 | SP7 |
| G3BP1 | enzyme | 0.0463 | 1 | SP7 |
| HTR4 | G-protein coupled receptor | 0.0349 | 1 | SP7 |
| DSPP | other | 0.0349 | 1 | SP7 |
| HIVEP2 | transcription regulator | 0.0349 | 1 | SP7 |
| FZD2 | G-protein coupled receptor | 0.0406 | 1 | SP7 |
| FZD6 | G-protein coupled receptor | 0.0349 | 1 | SP7 |
| ASARM-PO4 | chemical reagent | 0.0463 | 1 | SP7 |
| phenylamil | chemical reagent | 0.0349 | 1 | SP7 |
| chitosan | chemical - endogenous mammalian | 0.0176 | 1 | SP7 |
| CLASP2 | other | 0.0234 | 1 | SNAP25 |
| ZFP36L2 | transcription regulator | 0.0406 | 1 | SNAP25 |
| forchlorfenuron | chemical reagent | 0.00591 | 1 | SNAP25 |
| DNAJC5 | other | 0.00591 | 1 | SNAP25 |
| APPL1 | other | 0.0234 | 1 | SNAP25 |
| RAB3B | enzyme | 0.0176 | 1 | SNAP25 |
| Fudan-Yueyang-Ganoderma lucidum | chemical - endogenous non-mammalian | 0.0118 | 1 | SLC2A4 |
| diazinon | chemical toxicant | 0.0176 | 1 | SLC2A4 |
| gliquidone | chemical drug | 0.0176 | 1 | SLC2A4 |
| 1,3-bis(4-hydroxyphenyl)-4-methyl-5-[4-(2-piperidinylethoxy)phenol]-1H-pyrazole | chemical reagent | 0.0463 | 1 | SLC2A4 |
| GPAT3 | enzyme | 0.0463 | 1 | SLC2A4 |
| ZNF407 | transcription regulator | 0.00591 | 1 | SLC2A4 |
| WDFY2 | other | 0.0406 | 1 | SLC2A4 |
| SNED1 | other | 0.0406 | 1 | SLC2A4 |
| SERPINA12 | other | 0.0349 | 1 | SLC2A4 |
| NSF | transporter | 0.0118 | 1 | SLC2A4 |
| VAMP7 | transporter | 0.0176 | 1 | SLC2A4 |
| SLC2A4RG | transcription regulator | 0.00591 | 1 | SLC2A4 |
| VPS45 | transporter | 0.00591 | 1 | SLC2A4 |
| SDHA | enzyme | 0.0406 | 1 | SLC2A4 |
| INPPL1 | phosphatase | 0.0349 | 1 | SLC2A4 |
| TBC1D4 | other | 0.0234 | 1 | SLC2A4 |
| TNKS | enzyme | 0.0406 | 1 | SLC2A4 |
| COL14A1 | other | 0.00591 | 1 | SLC2A4 |
| LNPEP | peptidase | 0.0176 | 1 | SLC2A4 |
| PF-8380 | chemical reagent | 0.0234 | 1 | SLC2A4 |
| 8-aminoadenosine | chemical reagent | 0.0118 | 1 | SLC2A4 |
| isoferulic acid | chemical - endogenous mammalian | 0.00591 | 1 | SLC2A4 |
| sinapinic acid | chemical - endogenous non-mammalian | 0.0118 | 1 | SLC2A4 |
| potassium perchlorate | chemical reagent | 0.0292 | 1 | SLC2A4 |
| DIM-C-pPhOH-3-Cl | chemical reagent | 0.0349 | 1 | SLC2A4 |
| DIM-C-pPhOH-3,5-Br2 | chemical reagent | 0.0406 | 1 | SLC2A4 |
| duvoglustat | chemical drug | 0.0234 | 1 | SLC2A4 |
| PPP2R2A | phosphatase | 0.0349 | 1 | SLC22A4 |
| PPP2R5B | phosphatase | 0.0292 | 1 | SLC22A4 |
| SLC25A12 | transporter | 0.0292 | 1 | SLC18A2 |
| lobeline | chemical drug | 0.0176 | 1 | SLC18A2 |
| CD2AP | other | 0.0406 | 1 | SH3KBP1 |
| CDHR1 | other | 0.0349 | 1 | SERPINA3 |
| RNF17 | other | 0.0349 | 1 | SERPINA3 |
| zimelidine | chemical drug | 0.0406 | 1 | SERPINA3 |
| FBLN5 | other | 0.0406 | 1 | SERPINA1 |
| ADH5 | enzyme | 0.0118 | 1 | SDHA |
| raclopride | chemical drug | 0.0463 | 1 | SDHA |
| spiperone | chemical drug | 0.0292 | 1 | SDHA |
| BMS-214662 | chemical drug | 0.0118 | 1 | PRKCB |
| nardostachys chinensis extract | chemical reagent | 0.00591 | 1 | PRKCB |
| 3,4-dideoxyglucosone-3-ene | chemical - endogenous mammalian | 0.00591 | 1 | PRKCB |
| enzastaurin | chemical drug | 0.0406 | 1 | PRKCB |
| epalrestat | chemical drug | 0.0176 | 1 | PRKCB |
| cicletanine | chemical drug | 0.00591 | 1 | PRKCB |
| ingenol mebutate | chemical drug | 0.0406 | 1 | PRKCB |
| miR-377-5p (and other miRNAs w/seed GAGGUUG) | mature microRNA | 0.00591 | 1 | PRDX6 |
| 4-CMTB | chemical reagent | 0.0176 | 1 | PPARGC1A |
| Katp Channel (family) | group | 0.00591 | 1 | PPARGC1A |
| Nfatc | group | 0.0292 | 1 | PPARGC1A |
| GP7 | chemical reagent | 0.0349 | 1 | PPARGC1A |
| KCP | other | 0.0118 | 1 | PPARGC1A |
| (S)-3-hydroxy-2-methylpropanoic acid | chemical - endogenous mammalian | 0.0118 | 1 | PPARGC1A |
| AMG-9810 | chemical reagent | 0.00591 | 1 | PPARGC1A |
| Mitochondrial complex 1 | complex | 0.0118 | 1 | PPARGC1A |
| Tug1 | other | 0.00591 | 1 | PPARGC1A |
| CHCHD5 | other | 0.0463 | 1 | PPARGC1A |
| KLF14 | transcription regulator | 0.0176 | 1 | PPARGC1A |
| YTHDF2 | other | 0.0349 | 1 | PPARGC1A |
| SLC25A33 | transporter | 0.0118 | 1 | PPARGC1A |
| FNIP1 | other | 0.0406 | 1 | PPARGC1A |
| ACOT13 | enzyme | 0.0349 | 1 | PPARGC1A |
| TXLNG | other | 0.0292 | 1 | PPARGC1A |
| ACOT11 | enzyme | 0.0234 | 1 | PPARGC1A |
| alisporivir | biologic drug | 0.0118 | 1 | PPARGC1A |
| OSTN | other | 0.00591 | 1 | PPARGC1A |
| HAO1 | enzyme | 0.0292 | 1 | PPARGC1A |
| miR-217-5p (and other miRNAs w/seed ACUGCAU) | mature microRNA | 0.0463 | 1 | PPARGC1A |
| GW 6471 | chemical reagent | 0.0292 | 1 | PPARGC1A |
| SLN | other | 0.0176 | 1 | PPARGC1A |
| ABCC9 | ion channel | 0.0234 | 1 | PPARGC1A |
| CAY10594 | chemical reagent | 0.0463 | 1 | PPARGC1A |
| cyclic des-acyl ghrelin (6-13) | biologic drug | 0.0118 | 1 | PPARGC1A |
| Cyp2a12/Cyp2a22 | enzyme | 0.0234 | 1 | PPARGC1A |
| oligomycin A | chemical - endogenous non-mammalian | 0.0118 | 1 | PPARGC1A |
| pterosin B | chemical reagent | 0.0234 | 1 | PPARGC1A |
| CGP 12177 | chemical reagent | 0.0176 | 1 | PPARGC1A |
| arginine methyltransferase inhibitor-1 | chemical reagent | 0.0349 | 1 | PPARGC1A |
| BRL 37344 | chemical reagent | 0.0349 | 1 | PPARGC1A |
| PQQ cofactor | chemical - endogenous non-mammalian | 0.0463 | 1 | PPARGC1A |
| trimetazidine | chemical drug | 0.0234 | 1 | PPARGC1A |
| cAMP-dependent protein kinase | complex | 0.0118 | 1 | PPARGC1A |
| K ATP Channel | complex | 0.0176 | 1 | PPARGC1A |
| FGFR3-TACC3 | fusion gene/product | 0.00591 | 1 | PPARGC1A |
| CRTC1-MAML2 | fusion gene/product | 0.0176 | 1 | PPARGC1A |
| adiporon | chemical reagent | 0.0406 | 1 | PPARGC1A |
| GSK3235025 | chemical reagent | 0.0234 | 1 | PPARGC1A |
| 10,12-tricosadiynoic acid | chemical reagent | 0.0349 | 1 | PPARGC1A |
| MHY1485 | chemical reagent | 0.0118 | 1 | PPARGC1A |
| RSAD2 | enzyme | 0.0349 | 1 | PPARD |
| afamelanotide | biologic drug | 0.0234 | 1 | PPARD |
| d18:1/48:2 omega-O-linoleoyl-ceramide | chemical - endogenous mammalian | 0.0176 | 1 | PPARD |
| IVNS1ABP | other | 0.0176 | 1 | PML |
| ZRL5P4 | chemical reagent | 0.0118 | 1 | PML |
| PISD | enzyme | 0.00591 | 1 | PCYT2 |
| xylitol | chemical - endogenous mammalian | 0.0349 | 1 | PCK1 |
| Li+ | chemical reagent | 0.0463 | 1 | PCK1 |
| ARRDC3 | other | 0.0234 | 1 | PCK1 |
| 3-(methylthio)propionic acid | chemical - endogenous mammalian | 0.0118 | 1 | PCK1 |
| SHP | group | 0.0234 | 1 | PCK1 |
| Gm10768 | other | 0.0176 | 1 | PCK1 |
| benfluorex | chemical drug | 0.0349 | 1 | PCK1 |
| hydroxyethylnorfenfluramine | chemical - endogenous mammalian | 0.0349 | 1 | PCK1 |
| propionate derivative | chemical - other | 0.0118 | 1 | PCK1 |
| C1QL3 | other | 0.0234 | 1 | PCK1 |
| DUSP9 | phosphatase | 0.0176 | 1 | PCK1 |
| Ins1 | other | 0.0349 | 1 | PCK1 |
| PKIA | other | 0.0234 | 1 | PCK1 |
| C1QTNF3 | other | 0.0349 | 1 | PCK1 |
| GAL3ST1 | enzyme | 0.0463 | 1 | PCK1 |
| THRSP | other | 0.0463 | 1 | PCK1 |
| isonicotinamide | chemical reagent | 0.00591 | 1 | PCK1 |
| DOCK5 | other | 0.0234 | 1 | PCK1 |
| ITPR3 | ion channel | 0.0118 | 1 | PCK1 |
| SLC2A2 | transporter | 0.0463 | 1 | PCK1 |
| CARHSP1 | transcription regulator | 0.0176 | 1 | PCK1 |
| aminooxyacetic acid | chemical reagent | 0.0234 | 1 | PCK1 |
| REC2923 | chemical reagent | 0.0176 | 1 | PCK1 |
| decanoic acid | chemical - endogenous mammalian | 0.0292 | 1 | PCK1 |
| NUBPL | other | 0.0292 | 1 | NDUFS3 |
| PCBP1 | translation regulator | 0.0463 | 1 | MYH10 |
| ZHX2 | transcription regulator | 0.0406 | 1 | Mup1 (includes others) |
| PNPLA8 | enzyme | 0.0463 | 1 | Mt1 |
| SLC39A4 | transporter | 0.0234 | 1 | Mt1 |
| cadmium sulfate | chemical toxicant | 0.0234 | 1 | Mt1 |
| N-methyl-3,4-methylenedioxyamphetamine | chemical drug | 0.0463 | 1 | Mt1 |
| Cr6+ | chemical reagent | 0.0292 | 1 | Mt1 |
| mercury | chemical toxicant | 0.0406 | 1 | Mt1 |
| NUMB/NUMBL | group | 0.00591 | 1 | MSTN |
| ostarine | chemical drug | 0.0176 | 1 | MSTN |
| SENP2 | peptidase | 0.0406 | 1 | MSTN |
| olomoucine | chemical - kinase inhibitor | 0.0406 | 1 | MAP2 |
| pregnanolone | chemical - endogenous mammalian | 0.00591 | 1 | MAP2 |
| coconut oil | chemical drug | 0.0406 | 1 | MAP2 |
| tri-o-cresyl phosphate | chemical reagent | 0.0234 | 1 | MAP2 |
| 1-methyl-D-tryptophan | chemical drug | 0.00591 | 1 | MAP2 |
| GALNS | enzyme | 0.0234 | 1 | MAP2 |
| Dst | other | 0.0118 | 1 | MAP2 |
| opaganib | chemical drug | 0.0176 | 1 | MAP2 |
| MME | peptidase | 0.0234 | 1 | MAP2 |
| betel quid extract | chemical reagent | 0.00591 | 1 | MAP2 |
| SKF-83959 | chemical reagent | 0.0176 | 1 | MAP2 |
| 3beta-methoxypregnenolone | chemical drug | 0.00591 | 1 | MAP2 |
| CYM-5520 | chemical reagent | 0.0234 | 1 | MAP2 |
| ergothioneine | chemical - endogenous non-mammalian | 0.00591 | 1 | MAP2 |
| E-64c | chemical - protease inhibitor | 0.00591 | 1 | MAP2 |
| FMOD | other | 0.00591 | 1 | LUM |
| PCAT6 | other | 0.00591 | 1 | KLHL12 |
| CpG ODN 1555 | chemical reagent | 0.0292 | 1 | IL15RA |
| bis(3',5')-cyclic diguanylic acid | chemical - endogenous non-mammalian | 0.0406 | 1 | IL15RA |
| nandrolone decanoate | chemical drug | 0.0234 | 1 | GABRA5 |
| testosterone cypionate | chemical drug | 0.0234 | 1 | GABRA5 |
| methyltestosterone | chemical drug | 0.0292 | 1 | GABRA5 |
| USP49 | peptidase | 0.00591 | 1 | FKBP4 |
| AKR1C3 | enzyme | 0.0234 | 1 | FKBP4 |
| 14,15-DHET | chemical - endogenous mammalian | 0.0176 | 1 | EPHX2 |
| CENPN | other | 0.0463 | 1 | ENO1 |
| INSL5 | other | 0.0292 | 1 | ENO1 |
| CACNA1C | ion channel | 0.0463 | 1 | ENO1 |
| ethyl protocatechuate | chemical reagent | 0.0292 | 1 | ENO1 |
| 1-alpha,24(R),25-trihydroxyvitamin D3 | chemical - endogenous mammalian | 0.0118 | 1 | CYP24A1 |
| 24R,25-dihydroxyvitamin D3 | chemical - endogenous mammalian | 0.0118 | 1 | CYP24A1 |
| 25-hydroxyvitamin D | chemical drug | 0.0118 | 1 | CYP24A1 |
| GC | transporter | 0.0176 | 1 | CYP24A1 |
| phosphorus | chemical reagent | 0.0234 | 1 | CYP24A1 |
| DP-001 | chemical drug | 0.0234 | 1 | CYP24A1 |
| 1beta,25-dihydroxyvitamin D3 | chemical drug | 0.0118 | 1 | CYP24A1 |
| 2-hydroxy-imino phenylpyruvic acid | chemical reagent | 0.0118 | 1 | CTBP2 |
| 2-keto-4-methylthiobutyric acid | chemical - endogenous mammalian | 0.0118 | 1 | CTBP2 |
| Becn2 | other | 0.00591 | 1 | CNR1 |
| lorglumide | chemical reagent | 0.0234 | 1 | CNR1 |
| GPRASP1 | transporter | 0.00591 | 1 | CNR1 |
| cannabinoid | chemical drug | 0.0292 | 1 | CNR1 |
| 4-diphenylacetoxy-1,1-dimethylpiperidinium | chemical reagent | 0.00591 | 1 | CHRM3 |
| thioperamide | chemical reagent | 0.00591 | 1 | CHRM3 |
| Astra 1397 | chemical reagent | 0.00591 | 1 | CHRM3 |
| quinuclidinyl benzilate | chemical reagent | 0.00591 | 1 | CHRM3 |
| atropine | chemical drug | 0.0406 | 1 | CHRM3 |
| N-methylscopolamine | chemical drug | 0.0118 | 1 | CHRM3 |
| 2-amino-5-azotoluene | chemical toxicant | 0.0463 | 1 | CASR |
| GABBR2 | G-protein coupled receptor | 0.0406 | 1 | CASR |
| RNF19A | enzyme | 0.00591 | 1 | CASR |
| cinacalcet | chemical drug | 0.0118 | 1 | CASR |
| TRDN | other | 0.0234 | 1 | CASQ1 |
| miR-9-3p (and other miRNAs w/seed UAAAGCU) | mature microRNA | 0.0349 | 1 | CAMTA1 |
| LGH447 | chemical drug | 0.0349 | 1 | ATP6V1A |
| LDHB | enzyme | 0.0406 | 1 | ATP5F1B |
| 4-aminopyrazolo(3,4-d)pyrimidine | chemical reagent | 0.0118 | 1 | APOE |
| hyodeoxycholic acid | chemical - endogenous mammalian | 0.0176 | 1 | APOE |
| 24(S),25-epoxycholesterol | chemical - endogenous mammalian | 0.0406 | 1 | APOE |
| triamcinolone | chemical drug | 0.0406 | 1 | APOE |
| DNAJA4 | other | 0.00591 | 1 | APOE |
| AFM | transporter | 0.0118 | 1 | APOE |
| Pzp | other | 0.0349 | 1 | APOE |
| lynestrenol | chemical drug | 0.0118 | 1 | APOE |
| 2-[[4-[(e)-styryl]phenoxy]methyl]oxirane | chemical reagent | 0.0234 | 1 | APOE |
| ALYREF | transcription regulator | 0.0118 | 1 | APOE |
| non-esterified fatty acid | chemical - endogenous mammalian | 0.0406 | 1 | APOE |
| gemcabene | chemical drug | 0.0234 | 1 | APOE |
| 20alpha-hydroxycholesterol | chemical - endogenous mammalian | 0.0406 | 1 | APOE |
| hydrocortisone phosphate | chemical drug | 0.00591 | 1 | APOE |
| 22-hydroxycholesterol | chemical - endogenous mammalian | 0.0118 | 1 | APOE |
| endocannabinoid | chemical - endogenous mammalian | 0.00591 | 1 | APOA1 |
| SGMS2 | enzyme | 0.0118 | 1 | APOA1 |
| LY 518674 | chemical drug | 0.0176 | 1 | APOA1 |
| LIPC | enzyme | 0.0463 | 1 | APOA1 |
| CFH | other | 0.0349 | 1 | APOA1 |
| IGHMBP2 | enzyme | 0.00591 | 1 | APOA1 |
| nevirapine | chemical drug | 0.0234 | 1 | APOA1 |
| bisperoxovanadium 1,10-phenanthroline | chemical reagent | 0.0463 | 1 | APOA1 |
| TUBB3 | other | 0.00591 | 1 | ANXA5 |
| buserelin | biologic drug | 0.0463 | 1 | ANXA5 |
| bacitracin | biologic drug | 0.0176 | 1 | ANXA5 |
| fertirelin | chemical reagent | 0.0118 | 1 | ANXA5 |
| marinobufagenin | chemical - endogenous mammalian | 0.0292 | 1 | ANXA5 |
| PLAA | other | 0.0292 | 1 | ANXA4 |
| SLCO1C1 | transporter | 0.0234 | 1 | ALDH1A1 |
| ARTN | growth factor | 0.0234 | 1 | ALDH1A1 |
| HPGD | enzyme | 0.0176 | 1 | ALDH1A1 |
| PDE1C | enzyme | 0.0234 | 1 | ALDH1A1 |
| ribociclib | chemical drug | 0.0292 | 1 | ALDH1A1 |
| CHS-828 | chemical drug | 0.0463 | 1 | ALDH1A1 |
| muscarine | chemical toxicant | 0.0406 | 1 | ALDH1A1 |
| CIL56 | chemical reagent | 0.0234 | 1 | ALDH1A1 |
| Tcf 1/3/4 | group | 0.0349 | 1 | AHSG |
| GW 9508 | chemical reagent | 0.0176 | 1 | AHSG |
| TDO2 | enzyme | 0.0406 | 1 | AHR |
| 2,2-(2-chlorophenyl-4'-chlorophenyl)-1,1-dichloroethene | chemical - endogenous mammalian | 0.0118 | 1 | AHR |
| darusentan | chemical drug | 0.0292 | 1 | ACE |
| PITHD1 | other | 0.0406 | 1 | ACE |
| CYP4A11 | enzyme | 0.0463 | 1 | ACE |
| temocapril | chemical reagent | 0.0176 | 1 | ACE |
| olmesartan medoxomil | chemical drug | 0.0349 | 1 | ACE |
| quinapril | chemical drug | 0.0349 | 1 | ACE |
| trigonelline | chemical - endogenous mammalian | 0.0349 | 1 | ACE |
| D,L-propargylglycine | chemical reagent | 0.0406 | 1 | ACE |
| fosinopril | chemical drug | 0.0176 | 1 | ACE |
| WNT16 | other | 0.0406 | 1 | ACADVL |
| WNT2B | other | 0.0292 | 1 | ACADVL |
| BML-284 | chemical reagent | 0.0349 | 1 | ACADVL |
| SKL-2001 | chemical reagent | 0.0234 | 1 | ACADVL |
